# Ancient co-option of LTR retrotransposons as yeast centromeres

**DOI:** 10.1101/2025.04.25.647736

**Authors:** Max A. B. Haase, Luciana Lazar-Stefanita, Lyam Baudry, Aleksandra Wudzinska, Xiaofan Zhou, Antonis Rokas, Chris Todd Hittinger, Andrea Musacchio, Jef D. Boeke

## Abstract

The evolutionary origins of the genetic point centromere in the brewer’s yeast *Saccharomyces cerevisiae,* a member of the order Saccharomycetales, are still unknown. Competing hypotheses suggest that the point centromere tripartite genetic centromere DNA elements (CDEs) either evolved from ancestral epigenetic centromeres by descent with modification or were gained through horizontal transfer from selfish DNA plasmids.^1,2^ Here, we identified centromeres in the sister order Saccharomycodales and termed them “proto-point centromeres” due to sequence features that bridge the evolutionary gap between point centromeres and ancestral centromeres types. Comparative genomic analyses across multiple yeast orders showed an unexpected evolutionary link between point and proto-point centromeres to the long terminal repeats (LTRs) of Ty5 retrotransposons. Strikingly, one Saccharomycodales species, *Saccharomycodes ludwigii*, harbors compact Ty5-based centromeres, where its CDEII elements are divergent AT-rich Ty5 LTRs. These living fossil centromeres show how retrotransposon cis-regulation was likely co-opted for genetic centromere specification. These insights show that point centromeres are direct descendants of retrotransposons and have evolved by descent with modification. Ultimately, the many diverse centromere types across the yeast subphylum may share a common ancestry rooted in retrotransposon activity.

## Main

Centromeric DNA was first described in the brewer’s yeast *S. cerevisiae* over 40 years ago.^3,4^ Since then, discovery of centromeres across many species have shown remarkable diversity in sequence, size, and specification mechanisms.^5^ Most centromeres are epigenetically determined via the histone variant CENP-A (Cse4 in yeasts)^6^, but in the yeast subphylum Saccharomycotina, a sequence-dependent mechanism replaced this ancestral epigenetic mode.^3,7,8^ The aptly named "point centromeres" are genetically defined by a short tripartite sequence organization: an essential AT-rich CDEII (∼50 to 200 bp, depending on the species examined^9^), flanked by the non-essential CDEI and the essential CDEIII, which bind Cbf1c and CBF3c, respectively.^10–15^ These elements nucleate assembly of a single inner kinetochore and enable a single chromosome-to-microtubule connection.^12^ These features are thought to be restricted to a single order of Saccharomycotina yeasts, the Saccharomycetales.^1^

Beyond Saccharomycetales, Saccharomycotina centromeres are diverse. In Pichiales, they include centromeres organized within Ty5 clusters (*Ogataea polymorpha*) or inverted repeats (*Komagataella phaffii*), or lack any obvious conserved structure (*Kuraishia capsulata*).^16–18^ Serinales species also show diverse centromeric sequences. The best studied of these are the centromeres of the human commensal and major pathogen *Candida albicans* and its close relative, *Candida dubliniensis*, which have unique 4–18 kb centromeres enriched with Cse4 and function epigenetically.^19–23^ However, the centromeres of *C. albicans* appear to be recently evolutionary derived and are not ancestral.^24^ Other Serinales species, like *Debaryomyces hansenii* and *Scheffersomyces stipitis*, have centromeres marked by Ty5 clusters^25,26^, while *Candida tropicalis* centromeres are found within inverted repeats.^27^ Additionally, *Saccharomycopsis fibuligera*, a species within the order Ascoideales, also harbors putative centromeres defined by Ty5 clusters.^28^ The evolutionary relationship between point centromeres and these other centromeres remains unclear (Figure S1a).

Previous evolutionary models, based on fewer than ten genomes, suggested two possible origins for point centromeres (1) descent from ancestral epigenetic centromeres or (2) evolution from the selfish 2µ plasmid. The latter hypothesis has been widely supported by evidence demonstrating similar biochemical properties of the 2µ partitioning locus and the point centromere.^1,2,29–33^ However, no direct molecular evidence of shared ancestry linking 2µ plasmids to point centromeres has been uncovered. Since 2µ is exclusive to Saccharomycetales^34–40^, the discovery of a non-Saccharomycetales species with transitional centromeres—blending characteristics of both point and other yeast centromeres—would fundamentally challenge this model. Species with such centromeres would likely encode the CDEIII-binding CBF3 complex with centromere sequences showing affinity to centromeres of species with point and non-point centromeres, providing compelling evidence for an evolutionary bridge between centromere types.

Recent studies revealed extensive genomic diversity within and between the 12 orders Saccharomycotina.^41^ Of particular interest is Saccharomycodales, the sister order to Saccharomycetales, which includes *Saccharomycodes ludwigii* and the *Hanseniaspora* (divided into a Slower-Evolving Lineage or SEL and a Faster-Evolving Lineage or FEL^42^). To shed light on potential transitional centromeres, we identified the centromeres of Saccharomycodales. Comparative synteny studies and evolutionary analysis spanning 400 million years, revealed an unexpected ancestry of Saccharomycodales point centromeres to LTR retrotransposons. We then mapped genes associated with epigenetic or genetic centromeres across 1,090 yeast species and inferred a plausible trajectory of centromere evolution in Saccharomycotina.

### Mapping centromeres of Saccharomycodales yeast

To reconstruct the evolutionary history of point centromeres we studied centromere structure in the sister order to Saccharomycetales yeasts, the Saccharomycodales (Figure 1a). Prior work in *Sa*. *ludwigii* indicated that they have point centromeres with non-canonical CDEIII sequences, but biochemical validation was lacking^43^. We therefore studied centromere structure in additional Saccharomycodales species from the genus *Hanseniaspora* (two SEL species, *H. occidentalis* and *H*. *vineae*, and one FEL species, *H. uvarum*). We generated *de novo* genome assemblies from Oxford Nanopore long read sequences that were then scaffolded to chromosome-level assemblies with chromatin conformation sequencing (Figure 1b, S1b–f, Table S1, for details see supplemental note 1). Using HiC data to map the interacting pericentromeric DNAs, we were able to confidently map the putative centromeres to small, un-transcribed, intergenic regions (<2.5 Kb; Figure S1g–i). Within each of these putative centromeric regions we identified short (<300 bp) regions of AT-rich DNA, reminiscent of the CDEII sequences of *S. cerevisiae* and suggesting functional centromeres (Figure S1h–i). Analysis of gene synteny near these putative centromeric indicated they are conserved among *Hanseniaspora* species and their close relative *Sa*. *ludwigii* (Figure S1j).

**Figure 1.**
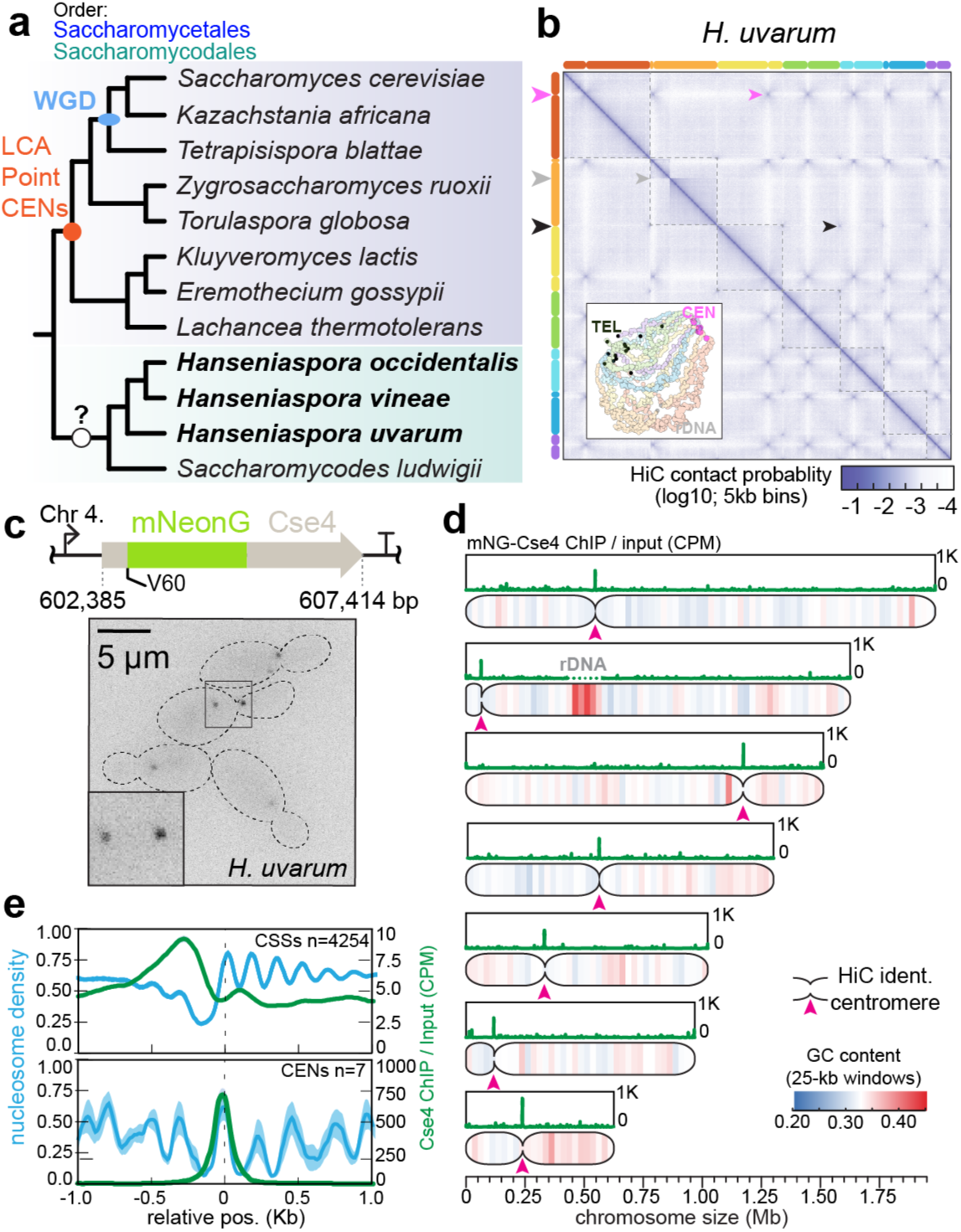
Mapping centromeric DNAs of Saccharomycodales yeasts. (**a**) Cladogram of Saccharomycodales and Saccharomycetales yeasts (topology from Opulente et al.) with labeled Whole Genome Duplication (WGD) node and “Point CENs” node indicating species with canonical point centromeres. (**b**) Hi-C-based chromosome assembly of *H. uvarum*, showing interacting regions: pericentromeric (pink arrow), telomeric (black arrow), and rDNA locus (gray arrow). (**c**) mNeonGreen (mNG)-tagged Cse4^CENP-A^ variant (inserted at Val 60) schematic and cellular localization micrograph. (**d**) MNase-ChIP-seq of mNG-Cse4^CENP-A^ in *H. uvarum*. Hi-C-identified centromeric regions marked with pink arrowheads; GC content (25 kb windows) plotted alongside. (**e**) Composite plots: nucleosome density (MNase-seq, left axis) relative to coding sequence start sites (CCS) and CDEII midpoints, with mNG-Cse4^CENP-A^ signal (right axis). Cse4^CENP-A^ signal is ∼100-fold weaker near CCSs than CDEII midpoints. Shaded blue regions represent the standard deviation of averaged MNase signals (for *CENs* = 7 and for CSS = 4254).

To verify the accuracy of our centromere mapping by HiC, we used the genetically tractable species *H. uvarum*^44,45^, where we inserted an internal mNeonGreen-tagged Cse4^CENP-A^ to map its centromeres by chromatin immunoprecipitation and sequencing (Figure 1c). We observed either one (G1) or two (G2) mNG::Cse4 foci per cell, the latter showing a characteristic G2/M separation of two foci across the mother-bud neck (Figure 1c). We then performed a cross-linked MNAse ChIP-seq experiment to map Cse4-containing mononucleosomes genome-wide (Figure 1d; Figure S2). Each chromosome had a single major peak of Cse4 that precisely intersected with the putative centromeric regions we identified using HiC (Figure 1d). Enrichment of Cse4 at putative centromeres was >100 fold greater than the -1 nucleosomes near coding sequence start sites, a region where Cse4 is also found in *S. cerevisiae* (Figure 1e).^46^ Lastly, inference of Cse4-nucleosome dyads supported the conclusion that Cse4 is present at a single nucleosome-sized AT-rich region of DNA per centromere (Figure 1e; Figure S2a). By mapping mono-nucleosomes showed that the Cse4-dyads are indeed contained within a single mono-nucleosome sized region (Figure S2a–c). This is consistent with the conclusion that the centromeres of *H. uvarum* wrap a single Cse4 nucleosome or nucleosome-like structure.

### Conserved and non-conserved sequence features between Saccharomycodales and Saccharomycetales centromeres

We next analyzed Saccharomycodales centromeres for enrichment of canonical CDEI/CDEIII motifs. While *H. vineae* and *H. occidentalis* centromeres contained multiple CDEI/CDEIII-like sequences, their arrangements were variable, unlike the conserved structure in Saccharomycetales (Figure 2a–b, S3a–b). In contrast, *Sa. ludwigii* and H*. uvarum* lacked CDEI/CDEIII but featured unique "GCG"-containing motifs flanking their CDEIIs (termed centromere-associated motifs, CAMs; see supplemental note 2 for a discussion on CAMs in relation to CDEIII; Figure 2c, S3c). CAMs in *H. uvarum* appeared near CDEII and at distal sites (Figure 2c, S2a), while *Sa. ludwigii* CAMs flanked CDEII in inverted orientations (Figure S3c).

**Figure 2.**
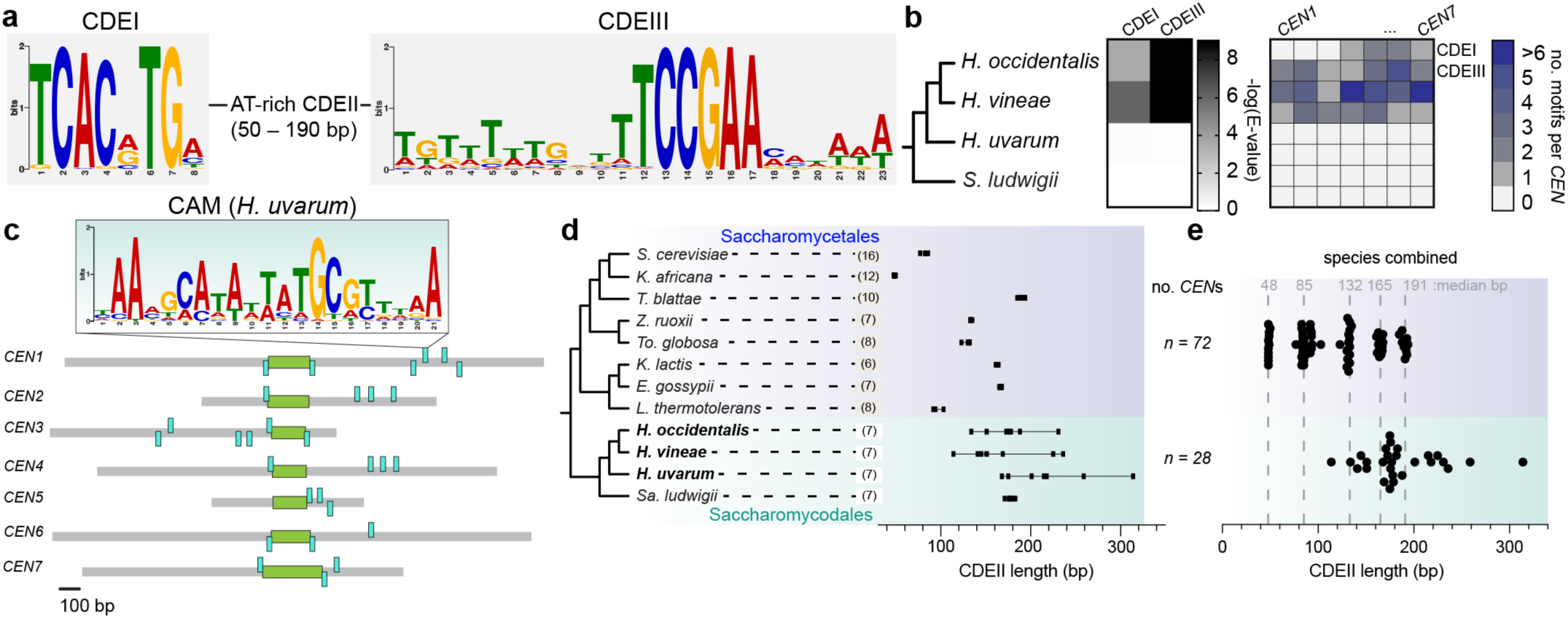
Centromeres of Saccharomycodales are sequence-flexible (**a**) Point centromere structure of Saccharomycetales yeasts, showing CDEI and CDEIII motifs derived from 172 centromeres (Yeast Gene Order Browser, YGOB). (**b**) Motif enrichment analysis of Saccharomycodales centromeric regions: left, log e-values for CDEI/CDEIII enrichment, average value of all centromeric regions; right, motif matches per centromere (for each species: CDEI: top row, CDEIII: bottom row). (**c**) *H. uvarum* centromere structure: CDEII regions (mNG-Cse4^CENP-A^ enriched, green) and DNA sequences matching conserved motifs (CAM, cyan). (**d**, **e**) CDEII length distributions: (d) across species; (e) by order (Saccharomycodales: n = 28, cyan, Saccharomycetales n = 72: blue), intervals of Saccharomycetales CDEII lengths are shown.

Saccharomycodales centromeres exhibited striking sequence flexibility. *Hanseniaspora* centromeres lacked unified ordering of sequences (CDEI, CDEIII, or CAM), with sequences appearing in variable orientations, copy numbers, and positions (Figure 2c, S3a–c). A most striking observation was that CDEII length also varied significantly in *Hanseniaspora* species (±41 bp variance vs. ±2 bp in Saccharomycetales; Figure 2d). *Sa. ludwigii* CDEII lengths were more uniform (177 ±4 bp), resembling the longest Saccharomycetales CDEIIs (e.g., *K. lactis*; Figure 2d– e). Notably, Saccharomycetales CDEIIs cluster at ∼30–40 bp intervals (e.g., ∼50 bp, ∼90 bp, ∼130 bp, 160 bp, or 190 bp), a pattern recently reported independently^9^, whereas Saccharomycodales CDEIIs are normally distributed around ∼185 bp (Figure 2e). This flexibility—in CDEII length, motif placement, and copy number— defines Saccharomycodales centromeres as distinct from Saccharomycetales.

### CDEII is necessary and sufficient for centromere function in *H. uvarum*

We next investigated the determinants of centromere function in *H. uvarum*. Classically, point centromeres subcloned into an episomal vector stabilize the vector during mitosis^3^. Unexpectedly, we observed an immediate and paradoxical negative impact on cellular fitness upon transformation of *H. uvarum* with its *CEN*s subcloned into an episome (Figure 3a). A plasmid stability assay, in which selection on the *LEU2* gene is relaxed for 6-8 generations, revealed that plasmids carrying *H. uvarum* centromeres were far more rapidly depleted from the population than control plasmids lacking the *CEN* sequences (Figure 3b-c). Two explanations for this paradoxical observation could be i) increased mitotic instability or ii) negative selective pressure on the maintenance of the *CEN* DNA. To disentangle these two, we performed a plasmid copy number analysis by whole genome sequencing (WGS) in cells under *LEU2* selective growth. We observed that the parental plasmid (no *CEN*) was maintained at ∼20 copies per diploid genome, whereas insertion of *CEN1*, *CEN4*, or *CEN7* resulted in single copy maintenance of each plasmid (Figure 3d). Therefore, despite being phenotypically ‘unstable’, the episomal *CEN*s are mitotically stable.

**Figure 3.**
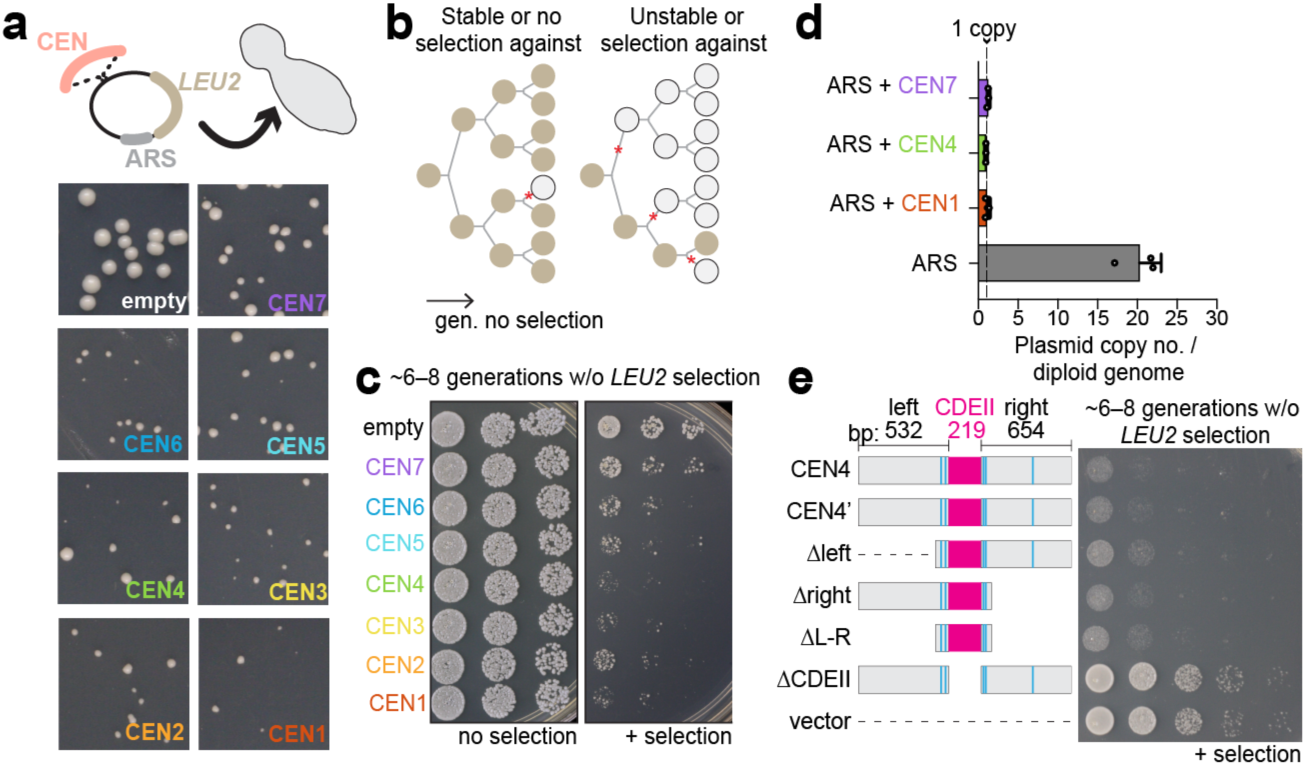
CDEII is necessary and sufficient for centromere function in *H. uvarum*. (**a**) Cloning *H. uvarum* centromeric regions into an episomal ARS vector caused growth defects in transformed strains (assayed on YPD agar at 30°C for 3 days). (**b**, **c**) Plasmid stability assay: plasmid loss by growth in rich medium (6–8 generations) followed by regrowth on selective Leucine dropout medium. (**d**) Centromere insertion mitotically stabilized the ARS plasmid (copy number analysis: average ± SD, n = 3). (**e**) Centromere deletion mutants confirm that CDEII is both necessary and sufficient for centromere activity. A *CEN4* plasmid where the ARS is inverted relative to the centromere is denoted by “*CEN4’* ”.

Using episomal *CEN*-induced cellular arrest as a proxy for centromere activity (for details on phenotypic and genetic characterization of growth arrest see supplemental note 3), we next asked whether the CAM and CDEII elements were necessary for centromere function. We examined a 1.4-kb region containing *CEN4* with flanking DNA on either side of the CDEII element (Figure 3e). Only upon deletion of the CDEII element did we observe relaxation of growth arrest. Importantly, a minimal 450-bp *CEN4* (containing the nearest flanking CAM sequences) was sufficient to elicit growth arrest (Figure 3e). Shuffling the sequence of CDEII, but maintaining its base composition, also resulted in relaxation of growth arrest (Figure S4), suggesting that specific sequence features of CDEII are essential for centromere function, likely the long poly-T and poly-A tracts, are essential for centromere function, as in *S. cerevisiae* CDEII^10,11^. Furthermore, mutating the sequence of flanking CAMs resulted in only a partial rescue of growth arrest (Figure S4). These experiments show that CAM sequences are necessary for optimal centromere function and that CDEII is necessary and sufficient for centromere function in *H. uvarum*.

### Saccharomycodales centromeres share a common evolutionary origin with point centromeres

The centromeres of Saccharomycodales may represent an evolutionary transition state from epigenetic to genetic point centromeres. In order to determine this, we inferred their common ancestry by reconstructing ancestral linkage groups (ALGs) across Saccharomycodales and non-WGD Saccharomycetales species (Figure 4). We identified 286 significant ALGs, with a median number of genes within each ALG of 4, consistent with the rapid decay of genome synteny during yeast evolution^47^.

**Figure 4.**
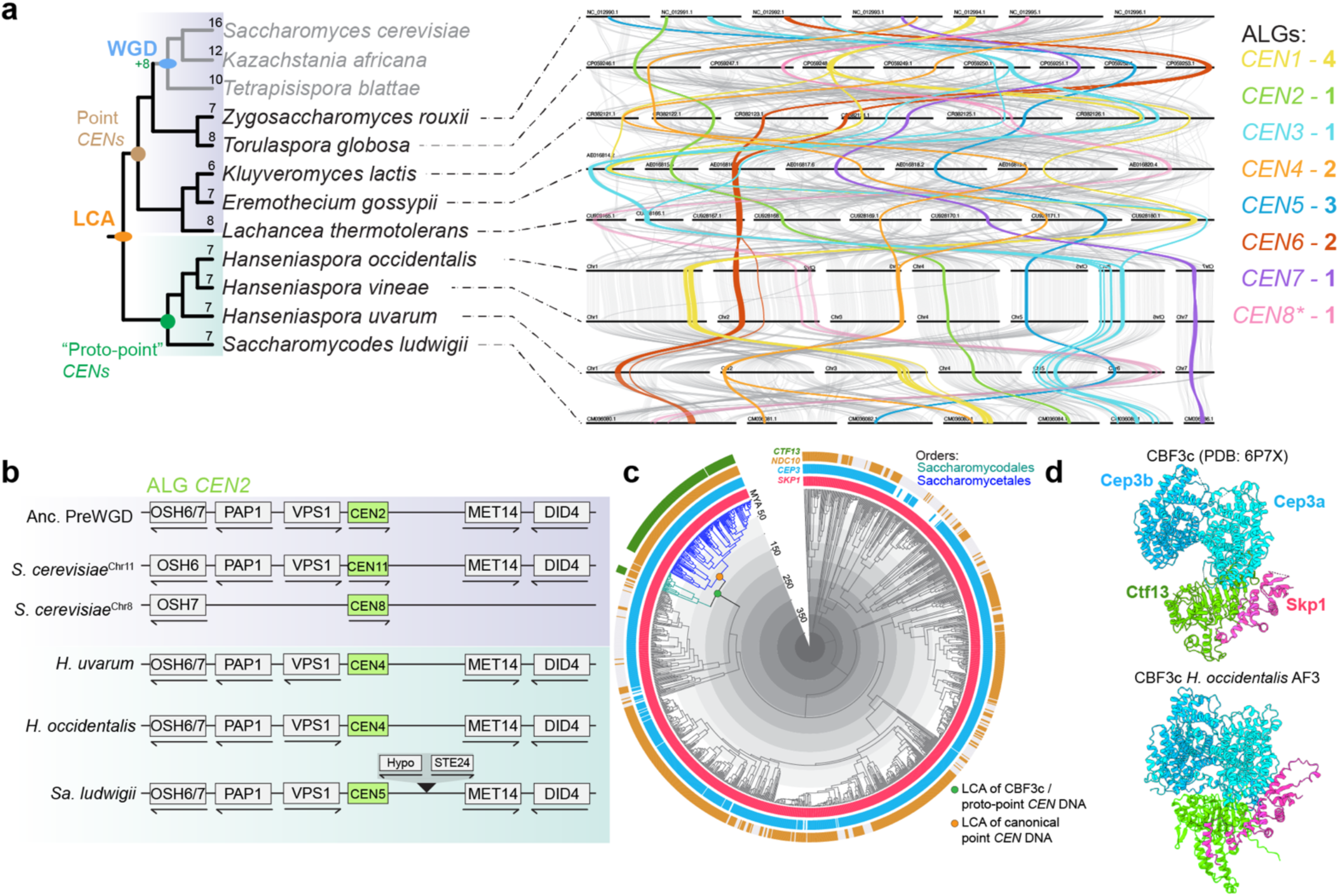
Common origin and evolution of Saccharomycodales and Saccharomycetales centromeres and DNA-adaptor proteins (**a**) Cladogram of Saccharomycodales and Saccharomycetales orders, with species haploid ploidy numbers indicated. Right: Centromere-linked ancestral linkage groups (ALGs) are color-coded by ancestral centromere numbering (YGOB); pairwise links between species chromosomes show ALGs are conserved across the two orders. ALG-*CEN8* is marked with an asterisk (*) as *CEN8* is absent in Saccharomycodales. (**b**) Example gene synteny of ALG-*CEN2* across select species. *Sa. ludwigii* shows two intervening genes from a derived insertion. (**c**) Presence and absence of CBF3c proteins across 1,090 yeast species (1,154 strains). Colored nodes indicate: (1) last common ancestor (LCA) with canonical point centromere DNA (orange); (2) LCA with CBF3c and ancestral centromeres of the two orders (green). Other yeast orders are left uncolored for clarity. (**d**) AlphaFold3 model of *H. occidentalis* CBF3 core complex and atomic model of *S. cerevisiae* CBF3c (PDB: 6P7X).

Centromere-associated ALGs contained more genes than average (Kolmogorov-Smirnov Test: *P* = 0.0161), suggesting selection against translocations or inversions in these regions (Figure 4; Figure S5a). From the perspective of *H. uvarum*, 3/7 centromeres were fully contained within a single ALG, while the remaining 4 spanned multiple non-contiguous ALGs (Figure 4b). Some ALGs were fragmented due to species-specific genome arrangements or annotation inconsistencies (Figure S5b–c, Figure S6, see supplemental note 4). In addition, the centromere of the *CEN8*-ALG was not conserved in Saccharomycodales, possibly due to centromere loss or gain (Figure S5g). Despite these complexities, ALG reconstructions strongly support a common evolutionary origin for the genomic regions containing centromeres between the two orders (Figure S5d–g, S6a–d, Table 1).

**Table 1.** Orthology among point and proto-point centromeres.

| ANCEN* | CEN1 | CEN2 | CEN3 | CEN4 | CEN5 | CEN6 | CEN7 | CEN8 | Order |
| --- | --- | --- | --- | --- | --- | --- | --- | --- | --- |
| *H. uvarum* | 3 | 4 | 6 | 2 | 5 | 1 | 7 | – | Saccharomycodales |
| *H. vineae* | 1 | 4 | 5 | 3 | 6 | 2 | 7 | – | Saccharomycodales |
| *H. occidentalis* | 1 | 4 | 5 | 3 | 6 | 2 | 7 | – | Saccharomycodales |
| *Sa. ludwigii* | 4 | 5 | 3 | 2 | 6 | 1 | 7 | – | Saccharomycodales |
| *S. cerevisiae* | 3 | 11 | 4 | 7 | 12 | 15 | 5 | 6 | Saccharomycetales (WGD) |
|  | 14 | 8 | 2 | 1 | 10 | 13 | 9 | 16 |  |
| *Z. rouxii* | 5 | – | 1 | 7 | 2 | 3 | 4 | 6 | Saccharomycetales (Non-WGD) |
| *T. globosa* | 3 | 1 | 7 | 4 | 5 | 8 | 6 | 2 |  |
| *K. lactis* | 1 | 2 | – | 3 | 6 | – | 4 | 5 | Saccharomycetales (Non-WGD) |
| *E. gossypii* | 1 | 2 | 5 | 6 | – | 3 | 4 | 7 | Saccharomycetales (Non-WGD) |
| *L. thermotolerans* | 8 | 4 | 7 | 5 | 2 | 3 | 6 | 1 | Saccharomycetales (Non-WGD) |

\*AncCEN nomenclature taken from the Yeast Gene Order Browser<sup>56</sup>.

The CBF3 complex (CBF3c: Cep3, Ndc10, Ctf13, Skp1), which binds CDEIII, is a defining hallmark of point centromeres. While Skp1 and Cep3 have homologs beyond Saccharomycotina, Ndc10 and Ctf13’s origins are debated. Prior hypotheses linked them to 2µ plasmid proteins,^1^ but our analysis of 1,154 yeast genomes suggested otherwise (see supplemental note 5). Critically, we observed that Ndc10 is widely conserved in Saccharomycotina and beyond (Figure 4c; Figure S7a–b, Table S4). Surprisingly, Saccharomycodales species encode Ctf13 homologs. These putative Ctf13 homologs from Saccharomycodales are supported as *bona fide* orthologs to Saccharomycetales Ctf13 due to conserved genomic synteny, phylogenetic affinity, and predicted structural similarities (Figure 4d, Figure S8a–g). Phylogenetic analysis of F-box containing proteins indicated that *CTF13* evolved as a paralog of the genes *DAS1* and *YDR131C* (Figure S9a–b). Strikingly, Saccharomycodales’ CBF3c subunits may assemble into a CBF3-like complex, which implies that the evolution of the CBF3c itself predated the evolution of canonical point centromere DNAs (Figure 4c–d; Figure S8f-g). Given that Cep3, Ndc10, and Skp1 are broadly conserved and that Ctf13 is found in at least two yeast orders and is related to conserved yeast genes, we can reject the hypothesis that the CBF3c subunits are descended from a common ancestor shared with 2µ proteins.

The presence of Ctf13 and a predicted CBF3c in Saccharomycodales yeast independently supports a common evolutionary origin for centromeres between the two orders. The Saccharomycodales centromeres exhibit distinct characteristics at the sequence level, demonstrating increased flexibility (Figure 2, Figure S3), yet they share notable similarities with point centromeres. These centromeres are short, approximately 185 base pairs in length, and are AT-rich, likely encompassing a single Cse4-containing structure per chromosome (Figure 1e, S2a). Given these features, which suggest an intermediate state between strictly genetic and non-genetic elements, we propose the term "proto-point" centromeres. This designation reflects their position as an early or transitional form, one potentially evolving towards more defined point centromeres.

### Point and Proto-point centromeres evolved from diminutive Ty5 clusters

Having established the common ancestry of Saccharomycodales and Saccharomycetales centromeres, we explored the deep evolutionary history of centromeres in Saccharomycotina yeast. The key clue came from *Sa. ludwigii* whose putative CDEII regions were flanked by repetitive sequences (Figure S3c, S10a,b), an observation not previously reported.^43^ Homologous repeats were found at non-CDEII positions, mapping to the LTRs of retrotransposons (Figure S10c). Notably, *Sa. ludwigii CEN2*, *CEN4*, and *CEN6* were associated with full-length LTR retrotransposons, complete with LTRs at both termini, but previously annotated as “ORFs” (Figure S10e–f). Phylogenetic analysis confirmed these sequences as Ty5 family members (Figure S10g, S11). Supporting this, *Sa. ludwigii* centromeric Ty elements contain target site duplications, characteristic yeast LTR sequence tip sequences (TGTTG and CAACA), and a Ty5 primer binding site complementary to the *Sa. ludwigii* initiator methionine tRNA half molecule (Figure S10h; Table S5) a feature unique to the Ty5 family of yeast elements. We term these elements *SaCEN*-Ty5 (for *<u>Sa</u>. ludwigii* <u>C</u>entromere <u>T</u>ransposon of <u>y</u>east).

Local DNA alignments confirmed the high repetitiveness of *Sa. ludwigii* centromeres due to Ty5 LTR fragments, with *CEN2* containing 237 intra-centromeric hits (Figure 5a, b; S10b). These LTRs share 62-95% nucleotide identity (Figure 5c), suggesting they are remnants of ancient retrotransposons. However, full-length *SaCEN-Ty5* elements themselves are highly conserved across *Sa. ludwigii* strains (Figure S12a-c), indicating recent transposition events supporting the conjecture that Ty5 elements are active in this species, just as they are in only a discrete subset of Saccharomycetales species.^48^ Interestingly, distant homology was detected between certain CDEII sequences (Figure 5a, c). For example, the CDEII’s of *CEN2* and *CEN3* shared 62% nucleotide identity (Figure 5c, d). Additional significant alignments are shown in Figure 5a and those overlapping CDEII sequences in Table S6. These findings suggest that the centromeres of *Sa. ludwigii* represent a remarkably well-preserved living transitional state between Ty5 cluster centromeres previously identified in many yeast orders (see next section) and the genetic point centromeres.

**Figure 5.**
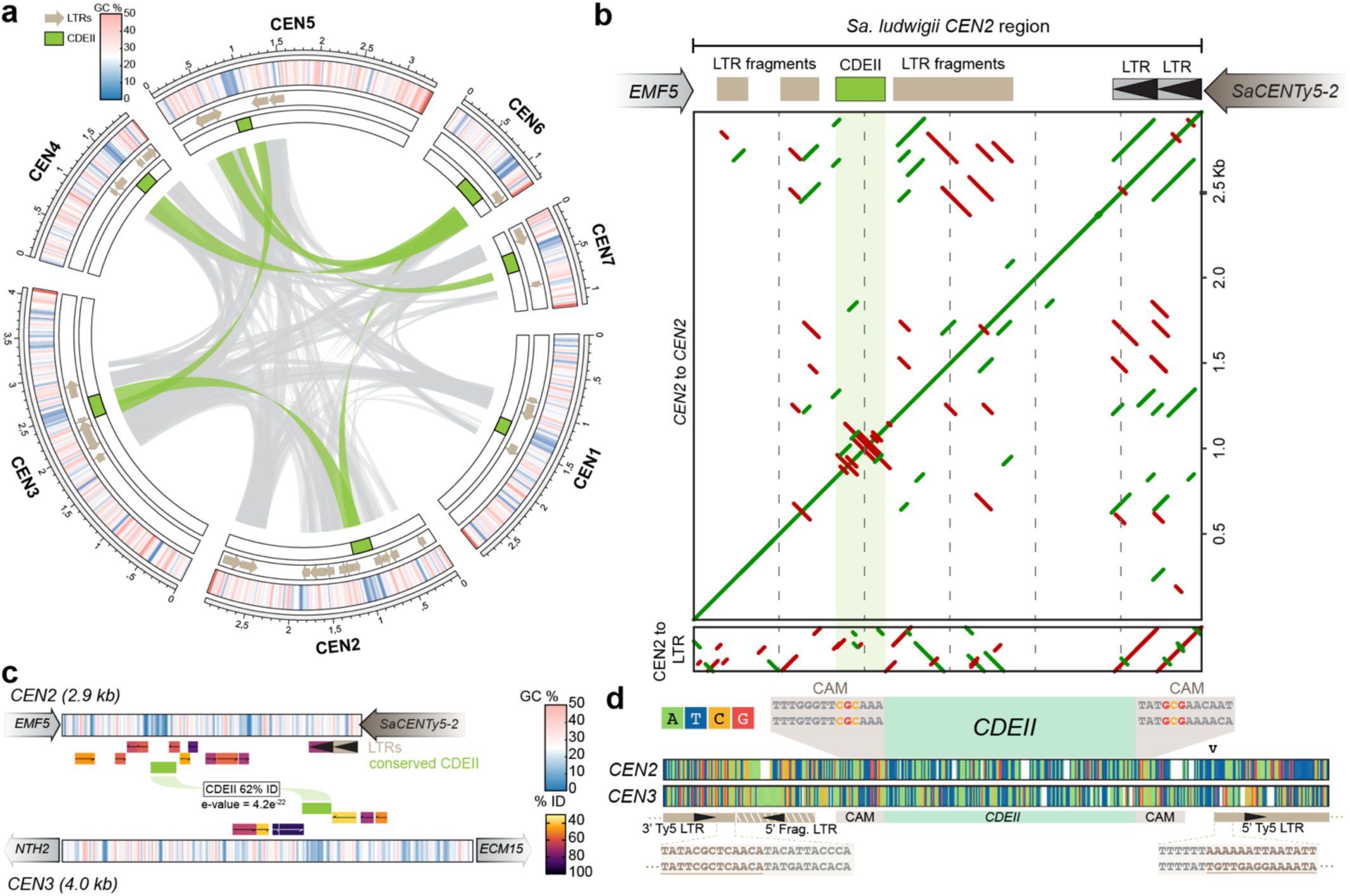
*Saccharomycodes ludwigii* proto-point centromeres are diminutive Ty5 LTRs clusters. (**a**) Circos plot of *Sa. ludwigii* centromere DNA alignments. Inner ring marks AT-rich CDEII regions. Arches connecting centromeres; green highlights alignments overlapping CDEII; gray alignments between LTR DNA. Annotated Ty5 LTRs are shown as beige arrows; GC content is plotted. Alignments were filtered at e-value < 0.01. (**b**) Local alignment of *Sa. ludwigii CEN2* reveals repetitive sequences in its intergenic region containing the putative CDEII. Shown below: alignment of *CEN2* to a consensus *SaCEN-*Ty5 LTR. Red and green line denote forward or reverse DNA alignments. (**c**) Detailed alignment between *Sa. ludwigii CEN2* and *CEN3*. Ty5 LTRs color-coded by percent identity to the most proximal LTR of *SaCEN*-Ty5-2. CDEII alignment is highlighted; and GC content is plotted. (**d**) Alignment of *Sa. ludwigii CEN2* and *CEN3* CDEII regions (*CEN2* is reversed from its genomic orientation). Expanded regions show conserved GCG motif, or its reverse complement CGC, and flanking regions of Ty5 LTR and LTR fragments.

### Molecular evidence of common ancestry between Ty5 LTRs, point, and proto-point centromeres

Ty5 clusters are tightly associated with centromeres in diverse yeast orders, including Pichiales, Serinales, and Ascoideales.^25,26,28^ How these Ty5 clusters relate to those of *Sa. ludwigii* is not clear. Therefore, we analyzed Ty5 distribution in fully assembled genomes, identifying Ty5 elements in 37% of species, which showed one of three enrichment patterns: telomeric, single-locus chromosomal, or nonspecific (Figure S10a, see supplemental note 6). Notably, we observed Ty5 clusters in *D. hansenii*, *O. polymorpha*, and *Sac. fibuligera*, with newly identified clusters in two *Cyberlindnera* species and in *P. terricola* (Figure S10). These Ty5 clusters span a striking size range, from small proto-point centromeres (1.2 kb in *Sa. ludwigii*) to large centromeric domains (131 kb in *Sac. fibuligera*), highlighting the structural diversity of Ty5 cluster centromeres (Figure 6a).

**Figure 6.**
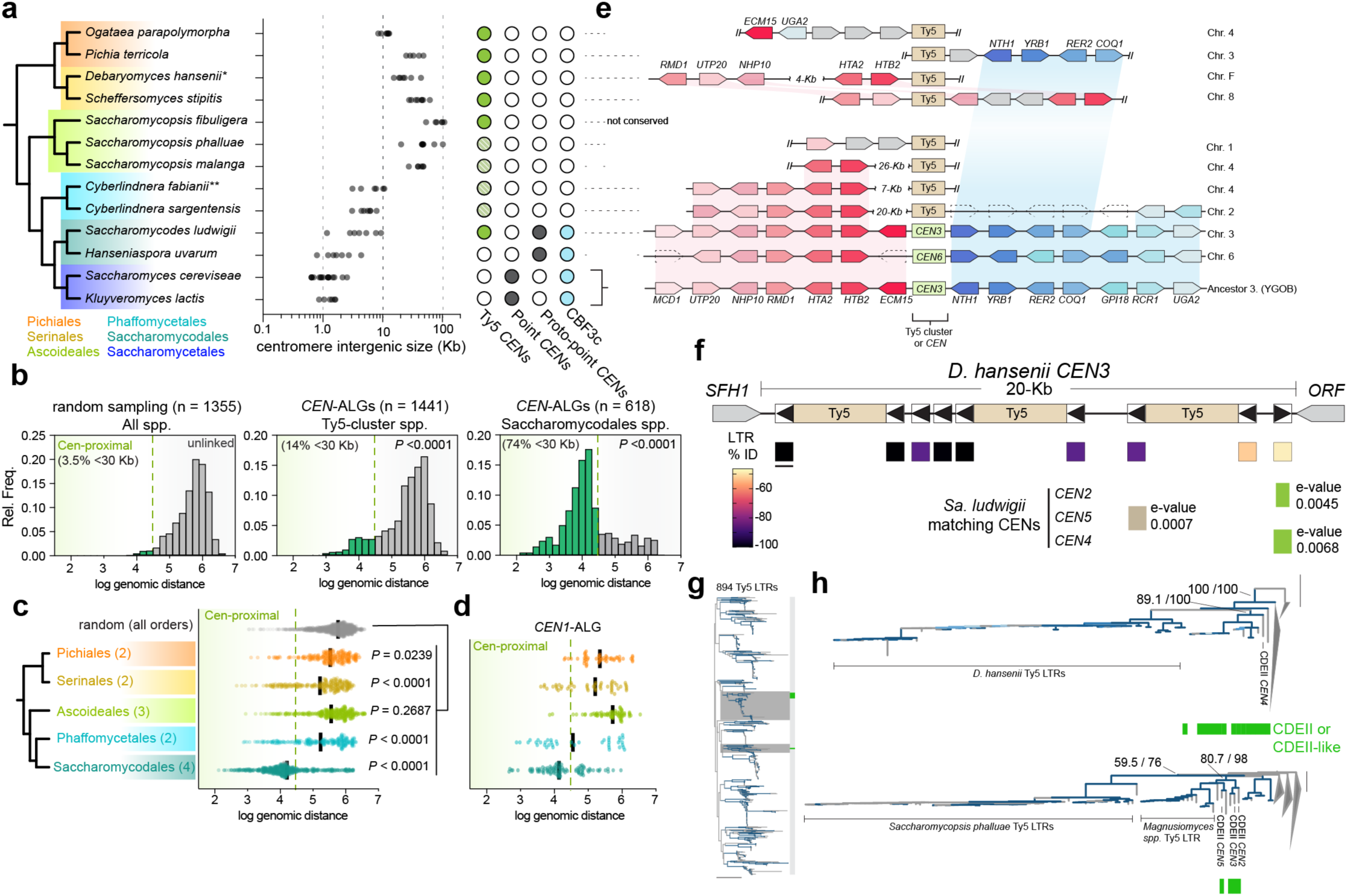
Molecular evidence of common ancestry of point, proto-point, and Ty5 LTR centromeres. (**a**) Cladogram of five Saccharomycotina yeast orders with Ty5 cluster centromeres (Saccharomycetales included; full details in Figure S13). Adjacent panels show centromere intergenic distances (log scale), centromere types, and presence of CBF3c. Filled circles denote experimentally validated or computationally inferred Ty5 cluster centromeres; partially filled circles indicate putative centromeres inferred from presence of Ty5 clusters. (**b**) Gene positioning relative to centromeres in species from panel (a). Left: distribution of distances from centromere of non-centromere-linked genes. Middle: distribution of distances from centromere of *CEN*-ALG-linked genes in Ty5 cluster species. Right: distribution of distances from centromere of *CEN*-ALG-linked genes in Saccharomycodales. Kolmogorov-Smirnov tests: comparisons between the distribution for unlinked genes and the distributions for either Ty5 cluster species or Saccharomycodales species. (**c**, **d**) Gene positioning relative to centromeres by order for all CEN-ALG genes (c) and CEN1-ALG specifically (d). P-values from two-sided t-tests with Tukey’s multiple tests correction. (**e**) Conserved synteny of centromere-linked histone H2A-H2B gene cluster. (**f**) Schematic of DNA alignments between *Sa. ludwigii* centromeres and *D. hansenii CEN3*. Brown/green shading highlights matches to *Sa. ludwigii* LTRs (brown) and CDEII (green); percent identity of *D. hansenii* LTRs on *CEN3* is noted by colored scale. (**g**) Maximum likelihood (ML) phylogeny of *Sa. ludwigii* CDEII sequences with 894 Ty5 LTRs (g). Zoomed subtrees (h) shows clades containing *Sa. ludwigii* CDEII and CDEII-like LTR sequences (scale: 1 substitution/site).

These diverse Ty5 clusters may have evolved independently or share a common ancestry. We found that the centromeric ALG genes identified above showed significant linkage to Ty5 cluster centromeres of other yeast species (Figure 6b–d), with well-preserved micro-syntenies (e.g., histone H2A-H2B and *NTH1*-*UGA2* blocks linked to Ty5 clusters in Serinales, Ascoideales, Pichiales, Phaffomycetales; Figure 6e, S14–15). Presence of these conserved micro-syntenies strongly support a shared evolutionary ancestry of the genomic regions encoding Ty5 cluster centromeres and point and proto-point centromeres (Figure S14–15). Absence of linkage to Ty5 clusters in *Blastobotrys aristatus* (Dipodascales; Figure S15i) placed the divergence of Ty5 clusters, point, and proto-point centromeres at the last common ancestor of Pichiales and Saccharomycetales (∼247–311 MYA).

Strikingly, local DNA alignments between the Ty5 clusters of *Sa. ludwigii* and other species revealed conserved sequences (Figure 6f, S16a). We not only identified significant matches between the species Ty5 LTR sequences, but, remarkably, also significant matches between a subset of *Sa. ludwigii* CDEII elements to newly discovered short CDEII-like sequences from other yeast Ty5 clusters (Figure 6f, S16b). These CDEII-like sequences, were enriched in AT content compared to their neighboring LTRs (Figure S16f), and showed phylogenetic affinity with *Sa. ludwigii* CDEIIs (Figure S16c–g). Notably, *Sa. ludwigii* CDEIIs and the CDEII-like sequences were confidently placed within the yeast Ty5 LTR family, suggesting they are divergent Ty5 LTRs (Figure 6g–h, Figure S16c–i). Thus, based on conserved micro-syntenies and directly identifiable sequence relationship of CDEII to Ty5 LTRs, as well as conservation of the Cbf3 complex that binds yeast centromeres, we conclude that Saccharomycodales and Saccharomycetales centromeres are descended directly from Ty5 cluster centromeres.

Finally, we explored how this evolutionary relationship to Ty5 LTRs might explain the evolution of the CDEI and CDEIII sequences. A plausible mechanism is Ty5 retrotransposition, which would bring Ty5 LTR *cis-*regulatory sequences directly into centromeres. By analyzing 975 yeast Ty5 LTRs, we identified 148 transcription factors with enriched binding sites (Figure S17a–b). Two of the most frequent class of transcription factors were zinc cluster (28%) and basic helix-loop-helix proteins (12%), which include Cep3 (CBF3c) and Cbf1, respectively (Figure S17b). Notably, 14.2% of all yeast Ty5 LTRs had canonical Cep3-binding sites, and 24.2% had canonical Cbf1-binding sites. Moreover, 42.4% of all LTRs were enriched with binding site of the zinc cluster protein, Ume6, which contains a “GCG” trinucleotide (Figure S17b). Co-occurrence of Cep3-binding sites and Cbf1-binding sites across LTRs from multiple yeast orders suggests that the cooccurrence of these *cis* sites may have been ancestral (Figure S17c). Thus, we propose that Ty5 retrotransposition is a reasonable mechanism for the evolution of CDEI, CDEII, and CDEIII at yeast centromeric DNA (Figure 7, Supplemental note 2).

**Figure 7.**
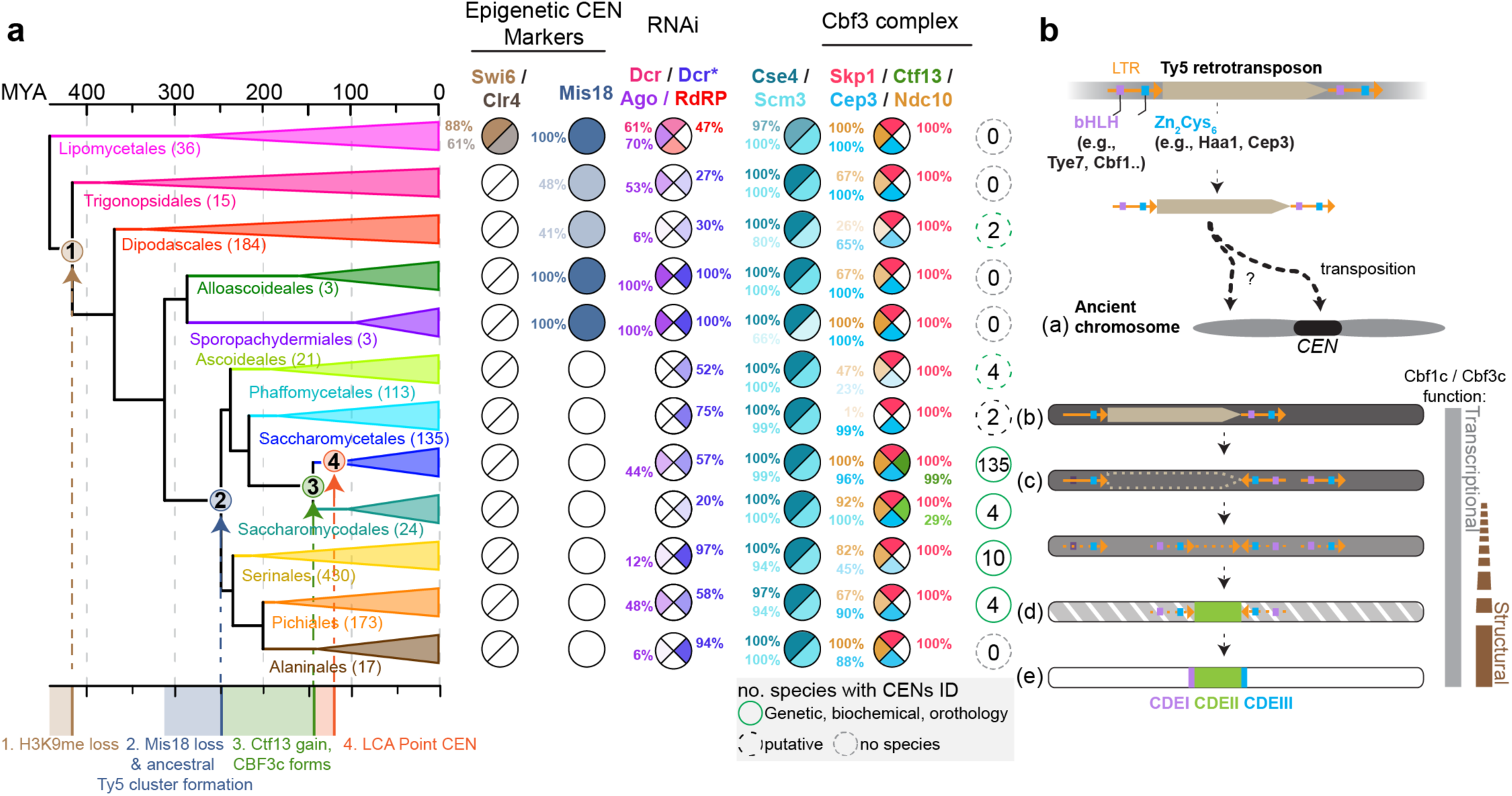
Tempo and mode of centromere evolution in Saccharomycotina yeasts. (**a**) Mapping gene presence on a time-calibrated phylogeny of twelve Saccharomycotina yeast orders (species count in parentheses)^41^ highlights three key centromere-related evolutionary events: (1) early loss of H3K9 methylation (Swi6/Clr4 retained only in Lipomycetales); (2) loss of Mis18, disrupting canonical epigenetic centromere maintenance; and (3) emergence of Ctf13 (via duplication of *DAS1*/*YDR130C*) in the ancestor of Saccharomycetales and Saccharomycodales. Ndc10, Cep3, and Skp1 (DNA-kinetochore adaptors) are deeply conserved, and with Ctf13, all predate point centromere DNA evolution (LCA marked at event #4). Green-dashed circles denote orders with at least one species having experimentally validated centromeres; all Saccharomycetales centromeres have been computationally predicted.^9^ Key innovations are mapped to timelines based on divergence times. (**b**) Model of Ty5 retrotransposon co-option for centromere function. An ancestral Ty5 element, regulated by basic helix-loop-helix (bHLH) protein and zinc cluster (Zn2Cys6) transcription factors, inserted into an epigenetic centromere or nearby. Selective pressures led to retention of Ty5 cis-regulatory elements flanking an AT-rich LTR-derived region enriched for Cse4^CENP-A^. Over time and in a lineage-specific manner, Ty5s were maintained, lost, reshuffled, or mutated – and in the ancestor of the Saccharomycodales and Saccharomycetales – to the point where only the cis-regulatory sites flanking an LTR-derived AT-rich region of Cse4^CENP-A^ enrichment remained. Cbf1 and Cbf3 complexes evolved structural roles in the kinetochore during this period and we predict the key innovation of Ctf13’s emergence that enabled Cbf3c function at centromeres/kinetochores. The proto-point centromeres diverged before the complete loss of flanking Ty5 elements, whereas point centromeres diverged only after the loss of centromeric Ty5 elements and the emergence of the strict spatial constraints of CDEI-II-III. See Supplementary Discussion for more details.

### Tempo and mode of ancestral epigenetic replacement

Ancestral Saccharomycotina centromeres were likely AT-rich, regional, and epigenetically defined by H3K9me2/3 heterochromatin (similar to Pezizomycotina and Taphrinomycotina^18,49,50^). However, the tempo and mode of how ancestral epigenetic centromeres were replaced by Ty5 clusters is not known. We therefore inferred this evolutionary process by mapping the key proteins that define epigenetic or genetic point centromeres across >400 million years of yeast evolution (1,154 genomes). We observed that Lipomycetales, the order sister to all other Saccharomycotina yeasts, retains Mis18 (epigenetic Cse4^CENP-A^ maintenance), Swi6^HP1α^, and Clr4^Suv39^ (H3K9me2/3), suggesting they have heterochromatic epigenetic centromeres (Figure 7a, S18). We inferred that Swi6 and Clr4 were lost early in Saccharomycotina evolution (>400 million years ago; MYA), but Mis18 was instead lost later as it persists in 5 orders (10.7% of species). Mis18 loss coincided with the inferred Ty5 cluster centromere emergence in the LCA of other orders Pichiales and Saccharomycetales (∼250–310 MYA). Therefore, replacement of ancestral heterochromatic epigenetic centromeres preceded first by the loss of H3K9me, followed by loss of Mis18 100 million years later.

RNAi is essential for epigenetic centromeres^51^, but we observed that RNAi is patchily distributed across the subphylum (Figure 7a). Some Lipomycetales species retain ancestral like RNAi systems as they encode genes similar to Dicer and RdRP, again supporting the idea they encode heterochromatic epigenetic centromeres (Figure 7a, S18b). The remaining orders lack RdRP and Dicer completely (Figure 7a). However certain species encode Rnt1-derived Dicer-like homologs.^52^ Argonaute is also completely absent in multiple orders and is repeatedly lost elsewhere (e.g., 12% of Serinales retain it). In total we observed the majority of species are predicted to lack functional RNAi (strains without Dicer or Dicer-like and Argonaute 906/1154). These results suggest that in Saccharomycotina yeasts RNAi does not correlate to centromeres type (e.g., point, proto-point, or Ty5 clusters), suggesting the dependence of RNAi for centromere specification^53^ was bypassed early on in Saccharomycotina evolution.

Lastly, we infer that emergence of Ctf13 predates canonical point centromeres, emerging 140–250 MYA (Figure 7a). We infer that point centromeric DNA evolved 110–140 MYA in Saccharomycetales’ LCA. Critically, we failed to detect any homologs of 2µ proteins outside of the Saccharomycetales (Figure S18c and methods), consistent with expansive surveys of plasmid diversity in yeasts.^34–40^ Thus, we infer that 2µ plasmids invaded the Saccharomycetales post-divergence of Saccharomycetales and Saccharomycodales, critically after CBF3c and proto-point centromere evolution. These facts argue strongly against the model of descent of point centromeres from 2µ plasmids. Centromere evolution in Saccharomycotina thus reflects a gradual transition over >400 million years from heterochromatic epigenetic centromeres, to Ty5 cluster centromeres, and finally to point centromeres (Fig. 7a, b).

## Discussion

The evolution of point centromeres in Saccharomycotina yeasts represents an exceptional transition from an epigenetic to genetic DNA element. As revealed here, this transition was driven by the co-option of Ty5 LTR retrotransposons. Our findings show that the CDEII of point and proto-point centromeres originated directly from Ty5 LTRs, evidenced by conserved genomic positioning of centromeres (Figure 4 and 6), sequence identity between CDEII regions of *Sa. ludwigii* and divergent Ty5 LTRs (Figure 6), and conservation of the CBF3 complex genes across orders (Figure 7). The proto-point centromeres of Saccharomycodales, exemplified by *Sa. ludwigii*, highlight the ancestral flexibility of centromeric DNA, which we infer to have been comprised of Ty5 clusters and LTR-related CDEII sequences. Domestication of Ty5 LTRs underscores the insight that point centromeres evolved vertically by descent with modification. The transition was thus not abrupt, but a gradual process involving Ty5 neocentromere formation and subsequent sequence refinement (Figure 7). These findings serve to highlight a complex interplay of protein complex innovation and retrotransposon sequence co-option occurring over tens to hundreds of millions of years of evolution. The evolutionary shift to genetic centromeres was likely in-part facilitated by protein adaptations, such as the emergence of the CBF3c complex via *CTF13* duplication, which we show preceded the strict genetic structuring of CDEI-II-III motifs. We therefore wonder if the ancestral function of CBF3c, specifically Cep3, was primarily transcriptional regulation of Ty5 retrotransposons rather than its modern-day role in centromere activity and kinetochore structure^12,14^. Ty5’s intrinsic preference to integrate in silent chromatin is distinct from other yeast retrotransposons which preferentially integrate at tRNAs^54^. Ty5’s integration into silent chromatin is dependent on interactions between its integrase and the protein Sir4^55^, which may have enabled its integration into ancestral centromeres, where its cis-regulatory elements were subsequently repurposed for centromere specification and kinetochore assembly. The small Ty5-rich centromeres of *Sa. ludwigii* might serve as a future model system to dissect the mechanistic roles of LTR retrotransposons in centromere function. Overall, we show that point centromere evolution represents an extreme example of LTR retrotransposon domestication shaping genomic architecture. In this case a selfish mobile element was co-opted as an indispensable genetic element for the division of its host’s chromosomes.

## Contributions

M.A.B.H. conceptualized the project, performed the formal investigation (genome assembly, HiC scaffolding, sequence and synteny analysis, experimental work), wrote the original manuscript, and prepared figures. L.L.S., conceptualized the project, performed the formal investigation (HiC experiments, HiC data analysis and interpretation, and centromere identification), prepared figures and wrote the original text. L.B., performed the formal investigation (genome assembly and HiC scaffolding). A.W., performed the formal investigation (Nanopore sequencing). X.Z., A.R., and C.T.H. provided genome annotations. A.M. provided funding. J.D.B. supervised the research and provided funding, and analyzed sequences. All authors edited the manuscript.

## Acknowledgments

We are thankful to the ARS Culture Collection (USDA) for their support in providing the type strains of the three *Hanseniaspora* species. This work was supported by a Rules of Life: Epigenetics 2 grant from the NSF (award number: 1921641) and NIH grant P01-AG051449 to J.D.B. Research in the Hittinger Lab is supported by the National Science Foundation (DEB-2110403), USDA National Institute of Food and Agriculture (Hatch Project 7005101), in part by the DOE Great Lakes Bioenergy Research Center (DOE BER Office of Science DE–SC0018409), and an H.I. Romnes Faculty Fellowship (Office of the Vice Chancellor for Research and Graduate Education with funding from the Wisconsin Alumni Research Foundation). Research in the Rokas Lab is supported by the National Science Foundation (DEB-2110404) and the National Institute of Allergy and Infectious Diseases (R01 AI153356). A.M. acknowledges funding from the Max Planck Society, the European Research Council (ERC) Synergy Grant 951430 (BIOMECANET), the DGF’s Collaborative Research Centre 1430 "Molecular Mechanisms of Cell State Transitions", and the CANTAR network under the Netzwerke-NRW program. M.A.B.H is supported by a Marie Skłodowska-Curie grant (agreement No. 101206234). L.B. is supported by the Swiss National Science Foundation (Grant no. 182632).

## Supplemental Figure legends

**Figure S1.** Genome assemblies of *Hanseniaspora* species.

(**a**) Overview of the twelve orders of Saccharomycotina and species whose centromeres have been identified or whose putative centromeres have been identified. Time calibrated phylogeny is from Opulente et al. (**b**) Pulsed field electrophoresis analysis of *Hanseniaspora* spp. (**c-d**) 2D HiC interaction maps of *H. occidentalis* and *H. vineae*, (**e-f**) to the right are the corresponding 3D projections. (**g**) Example pericentromeric interaction between *H. uvarum*’s chromosome 4 and 5. Left, HiC contact probabilities between *H. uvarum*’s chromosome 4 and 5. Right, plotted as function of intensity is the intra-chromosomal interactions between chromosome 4 and 5. (**h–i**) Corresponding 5-Kb bins with maximal intra-chromosomal interactions are shown. For each the stranded RNA sequencing data is plotted, mono-nucleosome profile, and a 50-bp rolling window average of percent GC content. (**j**) Centromeres are orthologous amongst Saccharomycodales species. The conserved synteny is shown for one centromere across all species and the inferred ancestral state of Saccharomycetales species.

**Figure S2.** Cse4^CENP-A^ MNAse-ChIP-seq.

(**a**) Browser views of 6 kb windows around the seven centromeres of *H. uvarum*. Cse4 dyads are shown in green, a mono-nucleosome map (MNAse-seq) is shown as blue line, and a plot of GC content (50-bp rolling window average) in brown. CDEII regions are highlighted as are the CAM sequences (green bars). (**b**) Mono-nucleosome composite plot near the ATG codon of coding genes. (**c**) Cse4-nucleosome composite plot centered at the midpoint of CDEII elements. The cluster of CAM motifs of *CEN1*, *CEN2*, *CEN4* is highlighted as a minor Cse4 peak.

**Figure S3.** Centromere DNA element structure of the Saccharomycodales.

(**a–c**) Species-specific Centromere DNA elements and centromere structure of the Saccharomycodales, as indicated.

**Figure S4.** CDEII and CAM mutant effects on centromere function

The centromere induced growth arrest was performed with a *CEN4* CDEII mutant and CAM mutant. The CDEII mutant was made by scrambled the sequence of the CDEII region of *CEN4.* Likewise, the CAM mutant scrambled the two nearest CAM sites (immediately flanking the CDEII). Details on the specific sequences used are found in the Supplemental text.

**Figure S5.** Centromere ancestral linkage groups between Saccharomycodales and Saccharomycetales yeasts.

(**a**) Number of genes contained within the eight *CEN* ALGs is shown. (**b-c**) example of *CEN* ALGs plotted on relative genomic coordinates is shown for (**b**) *CEN2*-ALG and (**c**) *CEN4-*ALG. (**d-g**) Expanded *CEN* ALGs based on manual annotated synteny analysis. Continued in Figure S6.

**Figure S6.** Continued, Centromere ancestral linkage groups between Saccharomycodales and Saccharomycetales yeasts.

(**a-d**) Expanded *CEN* ALGs based on manual annotated synteny analysis. Continued in Figure S6.

**Figure S7.** Conservation of Ndc10-like proteins in Saccharomycotina and beyond.

(**a**) Domain organization of Ndc10 is shown for *Kluyveromyces lactis* based on prior work. Additionally, the corresponding Interpro domains are annotated below. Percent identity to *S. cerevisiae* Ndc10 is shown. (**b**) Domain structure and percent identity to *S. cerevisiae* Ndc10 of a selection of Ndc10 homologs. Species with genetic point centromeres are highlighted.

**Figure S8.** Saccharomycodales yeasts have a Ctf13 ortholog and form a predicted Cbf3 complex.y(**a**) CBF3 complex structure (PDB: 6gys) and diagram of arrangement of CDEI-II-III and the proteins bound to each. (**b**) Synteny analysis of the CTF13 gene in Saccharomycodales yeasts and outgroup species. The ancestral (Pre-Whole Genome Duplication) gene order is shown for Saccharomycetales yeasts. (**c**) Amino acid sequence alignment of the Ctf13 F-Box domain from Saccharomycetales and Saccharomycodales yeasts. (**d**) Alphafold3 models of CTF13 from *S. cerevisiae* and *H. occidentalis.* (**e**) An unrooted gene tree of Ctf13 is shown between orthologs of the two orders, with key genera marked. (**f**) Core CBF3 complex (Cep3, Skp1, Ctf13) Alphafold3 predictions between *S. cerevisiae* and *H. occidentalis* compared to the atomic model of *S. cerevisiae* Cbf3c (PDB: 6P7X). (**g**) Overview of the models obtained for the *H. occidentalis* Core Cbf3 complex Alphafold3 predictions.

**Figures S9.** *CTF13* is related to divergent yeast F-box LLR proteins.

(**a**) Unrooted phylogeny of orthologs of *S. cerevisiae* F-box containing proteins from 1,154 yeast species. (**b**) Gene tree of Ctf13, Das1, and YDR131C. (**c**) Alphafold2 models (Ctf13, AF-P35203-F1-v4; Das1, AF-P47005-F1-v4; YDR131C, AF-Q03899-F1-v4) of the indicated protein. (**d**) Amino acid sequence alignment of the leucine-rich-repeat insert domain from Ctf13, Das1, and YDR131C. (**e**) Protein identity matrix of Ctf13, Das1, and YDR131C sequences shown in panel d.

**Figure S10.** *Saccharomycodes ludwigii* centromeres are diminutive Ty5 clusters.

(**a**) Dotplot local DNA alignments of the seven *Sa. ludwigii* centromeric regions. (**b**) Summary of local DNA alignments between each centromeric region as a network diagram. (**c**) Proportion of repeat blastn hits that are centromeric or non-centromeric. To the right are diagrams of retrotransposon elements whose LTRs match the centromeric repeats. Pair-wise, percent identity between domains are shown, as are the percent identities between the LTRs and a consensus LTR sequence. (**d–f**) Dotplot alignments of the three *Sa. ludwigii* centromeres with SaCEN-Ty5 elements. (**g**) Maximum likelihood phylogeny of SaCENTy5 elements and related yeast Ty’s. (**h**) Diagram of the *SaCEN*-Ty5-4 element with *Sa. ludwigii* initiator methionine tRNA. Highlighted is the 5’ LTR sequence corresponding to the Ty5 primer binding site complementary to the anticodon stem-loop of *S. cerevisiae* initiator methionine tRNA. Below is a dotplot alignment of the centromeric region against a consensus *SaCEN*-Ty5 LTR sequence.

**Figure S11.** Primary sequence support *SaCEN*-Ty5 as a yeast Ty5 family member.

(**a–e**) Amino acid sequence alignments of the indicated Ty protein domains. (**d**) The reverse transcriptase YVDD motif is highlighted.

**Figure S12.** Comparison of *SaCEN*-Ty5’s from *Sa. ludwigii* strains

(**a**) DNA sequence identity matrix of *SaCEN*-Ty5’s LTRs compared to Tkp5 and Ty5-6. (**b**) DNA sequence identity matrix of full length *SaCEN*-Ty5’s compared to Tkp5 and Ty5-6. (**c**) Maximum likelihood phylogeny of full length *SaCEN*-Ty5’s with Tkp5 and Ty5-6. (**d**) DNA sequence identity matrix of yeast Ty5 LTRs. (**e**) Average sequence identity of Ty5 LTRs depending on the taxonomic level of comparison.

**Figure S13.** Examples of Ty5 clusters from Pichiales, Serinales, Ascoideales, Phaffomycetales, and Saccharomycodales.

(**a**) Survey of Ty5 elements from species with complete genome assemblies, as in main Figure 6. (**b-f**) Identity heat maps between whole chromosomes and putative centromeric regions for the indicated species. Repetitive sequences correspond to Ty5 elements and Ty5 LTRs. Identity heat maps were made using the ModDotPlot visualization tool^57^ or as for *Sa. ludwigii* in the YASS DNA local alignment program^58^.

**Figure S14.** Conserved gene synteny of centromere- and Ty5 cluster linked genes from diverse yeast species.

**Figure S15.** Continued, conserved gene synteny of centromere- and Ty5 cluster linked genes from diverse yeast species.

(**a**) Conserved synteny of the indicated centromeres and Ty5 clusters. (**b–h**) Relative genomic distance analysis of the indicated CEN-ALG compared across orders. *CEN8*-ALG was not analyzed as it is not conserved between the Saccharomycodales and Saccharomycetales yeasts. Numbers in parentheses indicates the number of species analyzed. (**i**) Relative genomic distance analysis of all CEN-ALGs, with the addition of the outgroup order Dipodascales.

**Figure S16.** *Sa. ludwigii* CDEII is a divergent Ty5 LTR

(**a**) Local DNA alignment results between *Sa. ludwigii* centromeric and Ty5 cluster centromeric regions. Hits are plotted by their e-value and alignment length, those colored green were considered significant matches (e-value <0.05). (**b**) Exemplar DNA alignments of *Sa. ludwigii* CDEII *CEN2* to two CDEII-like sequences from *P. terricola* and *D. hansenii*. (**c**) Search strategy to uncover additional CDEII-like sequences from the three species which had significant matches to *Sa. ludwigii* CDEIIs. Left, ML phylogeny of *D. hansenii* Ty5 LTRs and CDEII-like sequences. (**d**) ML phylogeny of *P. terricola* Ty5 LTRs and CDEII-like sequences. (**e**) ML phylogeny of *Sch. stipitis* Ty5 LTRs and CDEII-like sequences. (**f**) ML phylogeny of the combined *Sa. ludwigii* CDEIIs, Ty5 LTRs and CDEII-like sequences from *D. hansenii*, *P. terricola*, and *Sch. stipitis*. (**g**) ML phylogeny of a profile-profile based alignment of yeast Ty5 LTRs and *Sa. ludwigii* CDEIIs. (**i**) ML phylogeny of yeast Ty5 LTRs and *Sa. ludwigii* CDEIIs. Ty5 LTRs are colored by the indicated species.

**Figure S17.** Motif enrichment analysis of Saccharomycotina Ty5 LTRs.

(**a**) Analysis overview. 975 Ty5 LTRs were used as the input to AME (Analysis of Motif Enrichment) against the YEASTRACT motif database. CDEI and CDEIII motifs are shown for reference. (**b**) Results of AME analysis broken down by the class of trans factor whose motif are enriched in Saccharomycotina Ty5 LTRs. Example motifs of zinc cluster proteins and bHLH are shown. Below each are the q-values and the percentage of yeast LTRs that have the indicated motif. (**c**) Overlap of enriched motifs on single LTRs is shown. To the right are species whose LTRs have the indicated motif enriched on single LTR sequence.

**Figure S18.** Presence and Absence analysis of the indicated proteins across 1,154 genomes.

(**a**) Detailed view of presence and absence of key genes as shown in Figure 7a. Inferred Mis18 loss events are highlighted by a blue star. (**b**) Zoomed in view of the Lipomycetales, showing the conservation of canonical RNAi-dependent heterochromatin factors. (**c**) Overview of 2µ Rep1 and Rep2 presence and absence analysis in Saccharomycotina – 2µ proteins are only found in Saccharomycetales. The most parsimonious interpretation is that *CTF13* and the ancestor of proto-point and point centromeres evolved before 2µ invasion.

**Figure S19.** Linear mitochondrial genome of *H. uvarum*

(**a**) Diagram of the assembled mitochondrial genome with terminal repeats highlighted. Each terminal repeat unit is colored according to its percent identity the RPU6 at the 5’ end. (**b**) The annotation of the mitochondrial genome is shown. Annotation was obtained using the MITOS2 de novo annotator. (**c**) Percent identity comparison to the mitochondrial genome of the *H. uvarum* strain MUCL 31704. The geographic location at date of each strain’s isolation is provided.

**Figure S20.** Characterization of the *CEN*-induced cellular arrest in *H. uvarum*

(**a**) Chromosome coverage analysis from whole genome sequencing of *H. uvarum* strains with indicated plasmids. (**b**) Correlation between transcripts per million from RNA sequencing replicates. (**c**) Chromosome-specific transcriptional changes (Log2FC compared to WT) are not observed based on the individual episomal *CEN* transformed. Black point, episomal ARS; red points, episomal *CEN1*; purple points, episomal *CEN4*; green points, episomal *CEN7*. Each point represents the average Log2FC value of each chromosome (averaged of all genes on a chromosome). Above the distribution of average Log2FC of each chromosome is shown. (**d**) Log2FC vs. average transcript abundance is shown for each strain with episomal *CEN*s. The gene *LEU2* is highlighted as its down regulation is direct a result of mitotic stabilization of the episomal vector. (**e**) Summary of differentially up regulated genes involved in meiosis progression. Below the enriched GO biological process terms are shown.

## Methods

### Data availability

All yeast strains and plasmids are available upon request to Jef D. Boeke. Raw whole genome, MNase, and RNA sequencing data were deposited to the Sequence Read Archive (SRA) and are available under the Bioproject ID PRJNA961205 with accessions SRR32695949–SRR32695985. Assembled genomes are available under the Bioproject ID PRJNA961205. Processed MNase and ChIP sequencing data are available at the Gene Expression Omnibus (GEO) accession GSE294411.

### Strains and media

Strains used are listed in Table S2 and plasmids in Table S3. Strains and plasmids are available upon request to J.D.B. *Hanseniaspora* strains were grown in standard rich medium (YPD; Yeast extract, Peptone, Dextrose) or in Synthetic complete (SC) dropout medium at 30 °C.

### DNA extraction and Nanopore sequencing

DNA for nanopore sequencing was prepared as previously described.^59^ Briefly, overnight *Hanseniaspora* spp. yeast cultures (∼5 mL YPD) were pelleted by centrifugation, washed with 1× PBS, and resuspended in 5 mL of spheroplast buffer (1 M sorbitol, 50 mM potassium phosphate, 5 mM EDTA, pH 7.5) supplemented with 5 mM DTT and 50 mg/mL zymolyase. Cultures were incubated at 30°C with shaking at 210 rpm for 1 hour. The resulting spheroplasts were collected via centrifugation at 2,500 g (4°C), gently washed with 1 M sorbitol, and treated with a proteinase K solution (final concentration: 25 mM EDTA, 0.5% SDS, 0.5 mg/mL proteinase K) for 2 hours at 65°C, with gentle inversion every 30 minutes. Genomic DNA was extracted twice using a 1:1 ratio of phenol:chloroform:isoamyl alcohol. The aqueous layer was treated with ∼10 µg of RNase A at 37°C for 30 minutes, followed by a final 1:1 extraction with chloroform:isoamyl alcohol. DNA precipitation was carried out using 0.1 volumes of 3 M sodium acetate (pH 5.2) and 2.5 volumes of ice-cold 100% ethanol, with inversion until visible DNA strands formed. High-molecular-weight DNA was spooled onto a pipette tip, washed in 70% ethanol, air-dried, and dissolved overnight in TE buffer (10 mM Tris-HCl, pH 8.0; 1 mM EDTA). DNA quantification was performed using the Qubit 1× dsDNA HS Assay reagent (Thermo, Q33231) on the Qubit Flex Fluorometer. For sequencing, genomic DNA was simultaneously tagmented and barcoded with the Oxford Nanopore Rapid Barcoding Kit (SQK-RBK004) following the manufacturer’s protocol. Barcoded libraries were pooled, purified, and concentrated using SERA-MAG beads (Cytiva, 29343052). The prepared library was immediately loaded onto a MinION R9.4.1 flow cell (SKU: FLO-MIN106.001) and sequenced on a GridION Mk1 device for 46 hours.

### HiC library generation

Cultures (YPD) were inoculated with the desired strain and grown overnight at 30°C. The following morning, each was diluted into 150 mL of YPD at a starting optical density (OD) of A600 = 0.25 and each was grown until an OD of A600 = 0.8-1.0. Cells were cross-linked in formaldehyde (3% [v/v]) for 20 minutes at room temperature and the reaction was quenched with glycine (300 mM). Cells were then collected by centrifugation and washed 2x in fresh YPD. Lastly, the cell pellet was frozen in liquid nitrogen and kept at -80°C until further processing. HiC experiments and library generation were then performed as previously described.^60–62^

### Genome sequencing and assembly

Raw fastq reads were first processed with porechop (v.0.2.4) to remove barcodes and adaptor sequences. We then generated *de novo* genome assemblies using Canu (v. 2.2; genomeSize=10 maxInputCoverage=100). Contigs were polished in the following manner: first, raw contigs were corrected using the ultrafast consensus module Racon (v. 1.4.17), followed by two sequential rounds of contig polishing with Medaka (v. 1.7). Finally, we performed three rounds of contig polishing with Pilon (v. 1.23) using publicly available Illumina sequencing datasets from each strain (SRA Accession: SRX5619117, SRX5619118, and SRX5619119). Next, the polished contigs were scaffolded using chromatin confirmation sequencing data (HiC) with the 3D-DNA pipeline (v180922).^63^ Scaffolded assemblies were manually corrected using the Hi-C map visualization and editor Juicebox (https://github.com/aidenlab/Juicebox). The final assembly statistics are reported in Table S1.

### Genome annotations

Genomes were annotated following the previously published methods of the Y1000+ consortium.^41^ Genome completeness was then assessed with BUSCO, which yielded expected results for *Hanseniaspora* species (Table S1).^42^

### HiC data analysis

The HICLib algorithm was used to generate contact maps from paired-end reads.^64^ Read-pairs were mapped independently using Bowtie2 (v.2.2.9 --very-sensitive, --rdg 500,3 --rfg 500,3)^65^ on the corresponding MboI-indexed reference sequence. Unwanted restriction fragments were filtered out (e.g., loops, non-digested fragments, as described by Cournac et al.^66^) and valid restriction fragments were binned into 5 kb bins. Contact maps were finally filtered and normalized as previously described.^67^

### Pulsed-field gel electrophoresis (PFGE)

Chromosomes from stationary yeast cultures were prepared in agar plugs using Certified Megabase Agarose (Bio-Rad, 1613108). Approximately ∼10 mg of wet cell pellet was used for preparation. Each cell pellet was then rapidly mixed with the Zymolyase solution (25 mg/mL 20T Zymolyase in 10 mM potassium phosphate, pH 7.5) and cooled (42 °C) low melting point agarose (0.5% in 100 mM EDTA, pH 7.5). This mixture was quickly mixed by pipetting and transferred to the agar plug molds. After setting (30 minutes), plugs were transferred to 50 mL Falcon tube containing 1 mL of 500 mM EDTA, 10 mM Tris, (pH 7.5) and incubated at 37 °C overnight. The following morning 400 µL of proteinase K solution (5 mg/mL proteinase K, 5% sarcosyl, 500 mM EDTA, pH 7.5) was added and sample incubated for five hours at 50 °C. Lastly, plugs were washed 1x in water, and then 3x in TE buffer. Plugs were stored at 4 °C in TE buffer until use. PFGE was then carried out using the running conditions for *Hansenula wingei* (*Wickerhamomyces canadensis*; Bio-Rad Catalog 170-3667). Chromosomes from *S. cerevisiae* (0.225–2.2 Mb, Bio-Rad; 1703605) and *W. canadensis* (1.05–3.13 Mb, Bio-Rad; 170-3667) were used as molecular weight standards. Chromosomes were then separated on a 0.8% agarose gel in 1x TAE buffer (Certified Megabase Agarose Bio-Rad, 1613108) following the program details for *Hansenula wingei* chromosomes, on a BioRad CHEF-mapper XA system.

### *H. uvarum* genetic transformations

*H. uvarum* strain HH04 was transformed by electroporation following a previously published method^44^. To integrate the mNeonGreen tagged Cse4 we first subcloned a DNA fragment encoding *H. uvarum* Cse4 (strain HH04) with mNeonGreen inserted at residue Valine 60, a region previously targeted for internal integration of GFP in *S. cerevisiae*^68^. The fragment was then cloned into a vector containing a hygromycin resistance cassette (hphMX, with the promoter and terminator of *H. uvarum TEF1*), flanked by homology arms targeting the native Cse4 locus (730 bp upstream and 997 bp downstream). 1 µg of linearized plasmid was used for the transformation. The plasmid Transformants were grown on YPD + 400 µg of hygromycin B for selection of the hphMX cassette. For episomal centromere vectors, we cloned each pericentromeric region identified by HiC into the vector pJJ3252 (*H. uvarum* ARS/Leu2 vector). Details on critical DNA constructs are shown in supplemental note 7.

### Cross-linked MNAse ChiP-seq and MNAse-seq

Cultures (YPD) were inoculated with the desired strain and grown overnight at 30 °C. The following morning, each was diluted into 150 mL of YPD at a starting optical density (OD) of A600 = 0.25 and each was grown until an OD of A600 = 0.8-1.0. Cells were cross-linked in formaldehyde (1% [v/v]) for 10 minutes at room temperature and the reaction was quenched with glycine and incubated for 5 minutes (125 mM). Cross-linked cells were then washed 2x in ice-cold PBS and frozen in liquid nitrogen at stored at -80 °C until use.

Pellets were thawed and spheroplasts generated as previously described^59^. Spheroplasts were then resuspended and washed in MNase digestion buffer (1M sorbitol, 50 mM NaCL, 10 mM TRIS-HCL (pH 7.4), 5 mM MgCL2, 0.5 mM spermidine, 0.075% NP-40, 1 mM β-mercaptoethanol). Next, chromatin was digested with MNase digestion buffer supplemented with either 10 units/mL or 1 units/mL MNase (Thermo Fisher Scientific Cat. #EN0181) and incubated at 37 °C for 45 minutes. Reactions were stopped by addition of EDTA (30 mM final). For total MNase mononucleosome maps, sample were then de-crosslinked and protein digested by adding SDS (0.5% final), proteinase K (20 mg/mL), and incubated for 1 h at 37°C and 2 h at 65°C. Digested DNA was purified by phenol-chloroform extraction, and precipitated with isopropanol. DNA was then resuspended in TE buffer with 1 mg/mL RNAse A and incubated at 37°C for 30 min. LAstly, DNA was cleaned with the Zymo DNA clean and concentrator kit according to the manufacturer’s specifications.

For Cse4-containing nucleosomes, following MNase digestions, samples were sonicated using a Branson digital sonifier (30% power, 2.5s on, 5s off, 40s total) and the lysate was clarified by centrifugation at 4 °C (5 minutes at 16,000 x g). Next the lysate was bound to ChromoTek mNeonGreen-Trap Magnetic Agarose beads at 4°C overnight. The next day beads were washed 10x times with MNase digestion buffer and DNA was directly purified by phenol chloroform extraction. Purified DNA was used as the input for the NEB Ultra II Library Prep Kit following the manufacturer’s specifications. Libraries were sequenced on an Illumina NextSeq 500 with paired-end 2 × 150 bp read chemistry.

### Cross-linked MNAse ChiP-seq and MNase-seq data analysis

Data analysis for the MNase sequencing was carried out as before.^59^ Demultiplexed reads were trimmed of adaptor sequences using Trimmomatic (v0.39). Processed reads were then aligned to the *H. uvarum* genome (HuvaT2T, this work) using the Burrows Wheeler aligner (BWA) mem algorithm (v0.7.7). For the mono-nucleosome analysis, we filtered reads with estimated insert sizes in the 120–180 bp range using SAMtools. Filtered reads were then used as input for mononucleosome analysis using the DANPOS (v2) pipeline. Pre-processing and genome alignment steps for the X-linked MNAse ChiP-seq was carried out as above. We compared the counts per million from cells with tagged Cse4-mNG and cells without tagged Cse4 (WT). The genome wide ratio of Cse4-mNG to WT were then visualized in the IGV browser and TBtools. Additionally, Cse4-mNG nucleosome dyads were determined with deepTools (v3.5.2) using the function “bamCoverage –Mnase”.

### Whole genome sequencing and RNA sequencing

For WGS, cells were grown in triplicate to saturation (two days at 30 °C) in SC-Leu medium. Cells were then collected by centrifugation and washed once in fresh medium and frozen at 80 °C. For RNA-seq cells were grown in triplicate to mid-log phase (0.6 – 0.8 OD600), placed on ice, collected by centrifugation at 4° C, and frozen in liquid nitrogen then stored at -80°C. Procedures for DNA and RNA extractions were carried out a previously described.^69^

For WGS 50 ng of purified genomic DNA was used for the input to the NEB Ultra II FS library prep kit for Illumina (NEB cat. E7805L) and libraries were sequenced using paired end 2 x 75 bp read chemistry on the NextSeq 500 platform. Analysis of ploidy levels was performed as follows. Raw reads were first processed to remove sequencing adaptors with Trimmomatic (v0.39). Reads were then aligned to a modified HuvaT2T genome file, which has the additional episomal DNA, using the BWA mem algorithm (v0.7.7). Chromosome ploidy levels and relative plasmid copy number was estimated as before.^69^ Since the strains of *H. uvarum* used in these experiments (HHO44 derivatives) are diploid, we normalized the chromosome copy number to an expected diploid genome of 2n = 14.

For RNA-seq, 100 ng of purified RNA was used to prepare total-RNA stranded libraries with the QIAseq Stranded Total RNA Lib Kit (Qiagen cat. 180745) following the manufacture’s specifications. Ribosomal rRNA was depleted with the QIAseq FastSelect-rRNA Yeast Kit (Qaigen cat. 334217). Libraries were then sequenced using paired end 2 x 75 bp read chemistry on the NextSeq 500 platform. Reads were process to remove sequencing adaptors and barcodes with Trimmomatic (v0.39). Finally, reads were aligned to the HuvaT2T genome using the Kallisto pseudoalignment program (v0.46.0) and data analyzed in the sleuth tool (v. 0.30.0) Computed differential expression values are found in Table S7.

### Motif enrichment analysis

To identity DNA sequences enriched in the centromeric regions Saccharomycodales yeasts we used the MEME suite. First, we scanned each centromeric region for the canonical CDEI and CDEIII sequences using the FIMO function. Then for each species centromeres we performed motif discovery analysis using the MEME function (options; -mod anr, -objfun classic, -nmotifs 10 -minw 6 -maxw 50). For *H. vineae*, *H. occidentalis*, and *Sa. ludwigii*, CDEII sequences were defined as the length of the AT-rich region between the two immediate flanking motifs.

### Ancestral linkage group inference

As point centromeric DNAs are the fastest-evolving DNA sequences in yeast genomes^70,71^, we determined relatedness between proto-point and point centromeres by conserved gene synteny of pericentromeric regions. To avoid confounding effects of paralogs from the WGD that occurred in a subset of Saccharomycetales species, we only used the genomes of species that did not experience WGD. Ancestral linkage groups were inferred between Saccharomycodales and Saccharomycetales yeasts using the protein-based synteny analysis software suite, odp (v0.3.2).^72^ Genome assemblies used in this analysis were downloaded from NCBI genome repository along with their annotations (Dataset S1). Centromere ancestral linkage groups (ALGs) were then manually annotated as ALGs flanking or completely spanning centromeres in extant species.

### Catalog of yeast Ty5 retrotransposons and Ty5 LTRs

Publicly available genome assemblies (as accessed on May 14^th^, 2024) were filtered based on the criteria of 1) assembled to at least “chromosome” level, 2) only “reference” strains, and 3) limited to the taxonomic group “Saccharomycotina” (NCBI Taxonomy ID: 147537). This resulted in a dataset of 77 species spanning 7 out of the 12 orders of Saccharomycotina yeasts (Dataset S2). Ty5 elements were then identified using tblastx searches with either *Saccharomyces paradoxus* Ty5 element (GenBank: U19263.1) or the *Sa. ludwigii SaCEN*-Ty5 element as the query. Significance threshold was manually established by blasting species with known Ty5 elements and comparing e-value scores for known Ty5 vs other yeast Ty retrotransposons. For example, significant matches to *SaCEN*-Ty5 from the species *D. hansenii* that corresponded to a *bona fide* full length Ty5 element had an average e-value <1e^-125^, whereas non-Ty5 elements had a minimum e-value no smaller than 1e^-68^, therefore we applied a lower cut off of <1e^-68^ as a putative Ty5 element. These cutoffs were consistent across the species examined. Ty5 and LTR sequences were then manually annotated for each species.

### Ancestral linkage group gene positioning analysis

Conserved synteny near point centromeres, proto-point centromeres, and Ty5 cluster centromeres was assessed by measuring the genomic distance from each gene to the nearest centromeric region. Genomic coordinates of putative centromeres inferred by Ty5 cluster presence are provided in Table S8. In order to estimate a null distribution of any random gene from its nearest centromere we randomly sampled without replacement 100 ORFs from the protein annotation of *Zygosaccharomyces rouxii* (GCF_000026365.1) using the *sample* function from the Python random module. We then used tblastn to obtain genomic coordinates for each query sequences (random set of genes or the genes in *CEN*-ALGs). For the conserved gene synteny maps we manually annotated centromeric regions and constructed the graphs.

### Phylogenetics of yeast LTRs

CDEII-like sequences were identified using the local DNA aligner YASS (v1.16; seed pattern = “very high sensitivity”, indels = 20%, mutations 35%) We input the centromeric regions from *Sa*. *ludwigii* as the query sequences and searched against the centromeric Ty5 clusters from Pichiales, Serinales, and Phaffomycetales species. Alignments were filter by their e-value, with alignments considered significant if e-value <0.05. LTR, CDEII, and CDEII-like sequences were aligned using MAFFT (v7), using the options G-INS-1 and --adjustdirectionaccurately. We then inferred a maximum likelihood phylogeny using IQ-Tree (v1.6.12) with the parameters -st DNA -m GTR+R4+F -bb 1000 -alrt 1000. For the profile-profile alignment of LTRs, we aligned LTRs from each species using MAFFT as above. Then each species-specific alignment was aligned to each other using the profile-profile alignment feature in Seaview (v5.05).

### Gene presence and absence in 1,154 genomes

To map the presence and absence of genes across the diversity of Saccharomycotina yeasts we downloaded the genome sequences from a recent study that sequenced the majority of all Saccharomycotina yeast species.^41^ We then conducted Hidden Markov Model-based sequence similarity searches. To build HMM profiles for each query gene we first used the *S. cerevisiae* ortholog to search the non-redundant protein sequence (nr) database using PSI-BLAST algorithm in blastp with an exclusion of the taxonomic group “Saccharomyces”. In cases where *S. cerevisiae* has no homolog of the query gene (i.e., Swi6/Clr4/Mis18) we used the appropriate ortholog from the fission yeast *Schizosaccharomyces pombe*. After 3 iterations, or reaching 500 homologs, we selected sequences based on a e-value threshold of 1 x 10^-3^. We then aligned homologs from each query gene using MAFFT (v7.520) with the “auto” option. HMM profiles were generated for each alignment using the hmmbuild function of the HMMER program (v3.3.2). Then, using each HMM profile we searched all genome assemblies for putative homologs using the hmmsearch function. To accurately call homologs, we first performed reciprocal blastp searches and removed sequences that did not best match the original query sequencd. In some cases, such as Cse4 and Skp1 we recovered a significant hit from nearly all species and therefore we stopped at this step. In the other cases, we then realigned all passing sequences and created a second HMM profile, “HMMrecip”. We then performed a second and final hmmsearch search using the HMMrecip profile. In general, this strategy resulted in an increased recovery of sequences that pass the e-value and score threshold cut offs of 0.01 and 50, respectively (e.g., for Cep3, the first HMM profile recovered orthologs from 421 species, whereas the Cep3 HMMrecip profile recovered orthologs from 1084 species).

For Clr4 orthologs, our HMM profile also recovered *SET2* homologs as significant hits. However, Clr4 is a much shorter peptide (*Sch. pombe* Clr4 is 496 AA and the average observed Saccharomycotina Clr4 is 426 AA) than Set2 (the average observed Saccharomycotina Set2 is 796 AA) and we used this feature to filter out the Set2 sequences. The average score of passing Clr4 homologs was 318 vs an average score of 97 for Set2 sequences.

For RNAi proteins we searched for Argonaute (Ago), Dicer (Dcr), and RNA-dependent RNA polymerase (RdRP) homologs, using *Sch. pombe* sequences as the initial queries. We also searched for homologs of the yeast-specific Dicer-like gene (Dcr*), derived from the Rtn1 RNaseIII, using the *C. albicans* sequence as the initial query. Presence of a Dicer-like homolog was based on the observation, for the species in question, if it encoded two genes which significantly matched the HMM-profile constructed from the *C. albicans* Dicer-like gene. For example, for *S. cerevisiae*, which only encodes the Rnt1 gene, we recovered only a single significant hit. Whereas, for *Naumovozyma castellii*, which encodes both genes, we recovered two significant hits. Therefore, we considered species with two significant hits as encoding both Rnt1 and Dicer-like and species with one significant hit encoding only Rnt1.

For the 2µ proteins Rep1 and Rep2 we applied a similar search strategy. Sequences were sourced from PSI-blast searches of both *S. cerevisiae* sequences. Sequential HMMprofile searches expanded recovery of Rep1 homologs, 34 sequences from the initial search and 59 sequences the second search. We recovered 11 significant Rep1 homologs from non-Saccharomycetales species. However, in all cases the sequences shared >90% amino acid identity to Saccharomycetales Rep1, indicating these sequences are most likely contaminating sequences in each genome assembly and not real homologs (Table S9). It should be noted that Rep1 homologs between *S. cerevisiae* to other *Saccharomyces spp.* are highly divergent (e.g., Rep1 between *S. cerevisiae* and *S. uvarum* shares 69% amino acid identity and Rep1 between *S. cerevisiae* and *S. kudriavzevii* shares 75% amino acid identity), so such extreme sequence conservation of Rep1 is not realistic between species of different orders. In sum, we identified 48 homologs of Rep1 all of which are only found in Saccharomycetales species. Searches for Rep2 homologs were less successful with the majority (17/26) of hits being contaminating sequences which 16/17 were >99% identical to *S. cerevisiae* Rep2 (Table S10). An additional Rep2 match from the species *Botryozyma americana* was <20% identical to *S. cerevisiae* Rep2. However, this sequence was 100% identical to the Rep2 sequence of *Zygosaccharomyces rouxii* (ASW33869.1). The remaining ten hits came from the genus *Zygosaccharomyces* (*Zs*. *pseudorouxii*, *Zs*. *parabailii, Zs*. *rouxii, Zs*. *sapae, Zs*. *gambellarensis*), and the species *Kluyveromyces lactis, Torulaspora* spp. yHMJ407*, Saccharomyces mikatae, and Zygotorulaspora danielsina* – all Saccharomycetales species.

We additionally performed a targeted approach to find Rep1 and Rep2 homologs in Saccharomycodales species. We downloaded all published genomes from NCBI (as accessed on March 24^th^, 2025) that belong to the order Saccharomycodales (n = 76 assemblies, representing 24 species; Dataset S3). We performed tblastn searches of each assembly using the diverse Rep1 and Rep2 homologs identified above. For Rep2 this yielded 16 hits that passed a lenient significance threshold of an e-value <0.05 (Table S11). Nine of these hits (all e-value <1.0E-09) corresponded to small scaffold (3.1 kb) from the assembly of *Hanseniaspora taiwanica* that when reversed searched against the ‘nr’ database was revealed to be a segment of the vector pUTD2 (CABRI collection number: LMBP 2903), whose replicon was derived from *S. cerevisiae* 2µ. The remaining hits were borderline matches (e-values between 0.047 and 0.012). Reciprocal blasts back to *S. cerevisiae* genome found these hits were significant matches to the nuclear genes *IDH1* (1 hit) and *MBA1* (6 hits). Therefore, we failed to detect any evidence of 2µ in Saccharomycodales – despite using a diverse set of 2µ Rep1/2 orthologs and multiple strains. For future genome assemblies we recommend preprocessing steps that remove reads mapping perfectly to known 2µ sequences, as they appear to be a frequent lab contaminates.

