## Supplemental Notes for "Ancient co-option of LTR retrotransposons as yeast centromeres"

**Supplementary Note 1** Genome assemblies and descriptions.

Chromosome level genome assemblies for *Hanseniaspora uvarum*, *Hanseniaspora* vineae, and *Hanseniaspora occidentalis* var. *occidentalis* were assembled from Oxford Nanopore reads with contigs scaffolded to chromosome level assemblies using conformation capture sequencing (HiC). We describe the assembly results for each species individually. For *H. uvarum*, we generated 22,795 Oxford Nanopore reads with a read N50 of 9,906 bp. These reads were then assembled using the canu assembler (v2.0; genomesize=8.8m, maxInputCoverage=100, canuIterations=1). The resulting draft genome consisted of 14 contigs with an contig N50 of 1,082,821 bp. Draft contig sequences were then polished with the ultrafast consensus module Racon (v1.4.17). To generate the alignments between raw nanopore reads and the draft assembly, we mapped reads using the minimap2 program (v2.24). After racon polishing, we performed two rounds of contig polishing with medaka (v5.2; -m r941_min_high_g303, -t 16, -b 25). Lastly, we performed three rounds of contig polishing with Pilon (v1.23). Illumina reads used for Pilon polishing for *H. uvarum* were sourced from pubically available data (SRX5619117) and reads were aligned to the draft assembly using bwa mem (v0.7.17). The final polished assembly consisted of 14 contigs with a contig N50 of 1,082,821 bp and a GC content of 32.19%. We then scafolded the final assembly to chromosomes using HiC or chromatin conformation capture sequencing. Scaffolding was done using the 3D-DNA pipeline (v180922) and the output was manaully verified in the HiC map viewer Juicebox. The final assembly consisted of 8 contigs with a contig N50 of 1,512,396 bp.

The same genome assembly method was done for both *H.* vineae and *H. occidentalis* var. *occidentalis*. For *H. vineae* the initial draft assembly from canu consisted of 38 contigs with a contig N50 of 846,583 bp and the final assembly consisted of 8 contigs with a contig N50 of 1,705,107 bp and a GC content of 37.56%. For *H. occidentalis* var. *occidentalis* the initial draft assembly from canu consisted of 41 contigs with a contig N50 of 715,189 bp and the final assembly consisted of 8 contigs with a contig N50 of 1,893,144 bp and a GC content of 34.91%.

For the three species each genome assembly contained seven chromosomes plus a mitocondrial assembly. For *H.* vineae and *H. occidentalis* var. *occidentalis* their mitochondrial genomes were assembled as 26,805 bp and 36,927 bp circlular pieces of DNAs, respectively. Whereas, for *H. uvarum* its mitochondrial genome was assembled as a 26,226 bp linear piece of DNA, consistent with the fact that *H. uvarum* has linear mitochrondrial genome.[1] The ends of *H. uvarum*'s linear mitochrondrial genome is capped by an inverted repeat (IR) of 7,755 bp and 7,754 bp, on the 5' and 3' ends respectively (Figure S19a). Each IR is composed of two segments. Nearest the ends are six tadem ~840 bp sub-repeats, with repeat-units nearest the end being most divergent (Figure S19a). These sub-repeats make up 5013 bp and 4683 bp, on the 5' and 3' ends respectively. The second segment of the IR are terminal sequences of approximately 2754 bp that are 98.73 percent identical. Excluding inverted repeats, *H. uvarum*'s mitochondrial genome is 11,126 bp (Figure S19b). Comparison to the previosuly published *H. uvarum* mitochrondrial genome revealed the mitochrondrial genomes are well conserved (Figure S19c).

For each species we obtained partial assemblies of their ribosomal DNA clusters. For *H. uvarum*, its rDNA cluster sits on the long arm of chromosome 2 and is assembled to 12 repeat units. Assembled within a portion of its rDNA cluster are Ty3-like retrotransposon elements. We estimated the total size of the ribosomal DNA cluster to be 1.67 Mb (we assembled only 0.11 Mb) with a total number of rDNA repeats to be at least 213 units, signifcntly higher than prior reports based on gel electrophoresis methods.[2] Our rDNA size estimates were based on the porportion of reads that map to the rDNA-containing chromosome relative to the reads that map to the assembled rDNA cluster. For example, 10,728 reads mapped to *H. uvarum* chromosome 2 and 5,640 of which mapped to the rDNA cluster, or ~52% of all reads from chromosome 2 are rDNA. We then used the ratio of rDNA-reads to chromosome 2 to infer the size of the rDNA cluster (length of chromosome 2 minus the assembled rDNA length is 1.5 Mb, so a ratio of 1.1 rDNA to chromosome 2 DNA equals 1.67 Mb of rDNA), with an explicited assumption that there is no isolation, nor sequencing bais of DNA from chromosome 2. For *H. vineae*, its rDNA cluster sits on the long arm of chromosome 3 and is assembled to four repeat units. We estimate that *H. vineae* rDNA cluster is at least 0.82 Mb (we assembled only 0.03 Mb), containing 101 repeat units. For *H. occidentalis* var. *occidentalis*, its rDNA cluster sits on the long arm of chromosome 3 and is assembled to ten repeat units. We estimate that *H. occidentalis* var. *occidentalis* rDNA cluster is at least 1.44 Mb (we assembled only 0.09 Mb), containing 186 repeat units. Our estimates are within the range of the number of rDNA repeats observed for *S. cerevisiae* (100 – 200 repreat units).[3]

Our assemblies also contained sub-telomeric regions and, for some chromosomes, were assembled from telomeric to telomeric repeat. Telomeric single stranded G-tails were non-canonical in *H. uvarum*, with a repeat-unit sequence of CAGGT, rather than the cononical repeat-unit sequence of GGGT, which was observed in *H. occidentalis* var. *occidentalis* and *H. vineae*.

Lastly, we were able to identify the MAT locus from each genome assembly based on tblastn searches of the *SLA2* gene, which immediately flanks MATalpha or MATa genes in orther *Hanseniaspora* and yeast species.[4] In both *H. uvarum* and *H. vineae* we observed a single MATa gene on chromosome four of both species. In *H. occidentalis* var. *occidentalis*, we instead observed that its MAT locus encodes a homolog of the MATalpha gene on its chromosome four. These results are consistent with these strains of *Hanseniaspora* species being heterothallic as only a single MATing loci is observed.

**Supplementary Note 2** CAM in relation to CDEIII

CAM sequences from *Sa. ludwigii* and *H. uvarum* contain a “GCG” motif (Figure 2c, Figure S3c). Both of these motifs could be approximated as “WWWWWGCGWWWWW” and similarly CDEIII could be approximated as “WWTCCGAA”. As both “A” and “T” are components of “W” the flanking bases nearest the GC-containing triplets in CAM and CDEIII are similar in base composition. While *H. vineae* centromeres contain a well defined “CCG” triplet, *H. occidentalis* is less clear, with equal propoation of either “CCG” or “GCG” triplets (Figure S3a, b). These obseravtions suggests that CAM and CDEIII may be evolutionary releated, as a single nucleotide substition is needed to transistion from one to the other. CDEIII and specifically the “CCG” triplet is recognized and bound by the Zinc-cluster domain of Cep3. Zinc-cluster cluster containing proteins recognize a range of consensus binding sites, including those that contain “CCG” (e.g., Cep3 or

Gal4) or “GCG” (e.g., Haa1 or Ume6) (Figure S17b). Given the significant enrichment of yeast Ty5-LTRs with Zinc-cluster binding sequences  (Figure S17b) and the simple nucleotide changes needed to transistion between the two GC-containing triplets we propose CDEIII and CAM are evolutionary related to a common ancestor that was most likely a cis-regulatory site in yeast Ty5 LTR. It will be important for furture work to test Saccharomycodales homologs of Cep3 binding substrate specifity on targets with either GC-containg triplets.

**Supplementary Note 3** Characterization of the *CEN*-induced cellular arrest in *H. uvarum*

We observed that in *H. uvarum* encoding its centromeres on an episomal DNA plasmid resulted in a paradoxical reduction to fitness, while leading to mitotic stability of the resulting episomal *CEN*s (Figure 3). To understand this phenotype better we characterized episomal *CEN*s transformed strains of *H. uvarum* to the ARS plasmid counterpart (i.e., the plasmid with a *CEN*). We first examined whole genome sequencing data from strains with *CEN1*, *CEN4*, or *CEN7* plasmid. This analysis revealed that strains transformed with episomal-*CEN*s presented aneuploidy of at least one or two chromosomes (Figure S20a). However, these aneuploids were uninfluenced by the particular episomal *CEN* transformed (e.g., we did not observe that *CEN1* induces Chr1 aneuploidy). Instead, strains, regardless of which episomal *CEN* was transformed, displayed a similar set of aneuploids (e.g. loss of Chr7 and gains of Chr1, Chr3, and Chr5). This is consistent with the idea that episomal-*CEN*s are inducing a common pathway that leads to chromosome division erorrs and therefore aneuploidy. To further investigate this phenomenon, we performed RNA sequencing on strains transformed with episomal *CEN*s (Figure S20b). In strains transformed with episomal *CEN*s, we observed a shared upregulation of genes whose orthologs in *S. cerevisiae* are involved in meiotic progression (sporulation) and spore cell wall biogenesis (Figure S20e). These global transcriptomic effects were independent of any specific aneuploidy (Figure S20b–e). We conclude that induction of a meiotic-like state by episomal *CEN*s DNA imposes a negative selective pressure against the maintenance of the episomal centromeric plasmid, thus explaining the paradoxical fitness defect and elevated rates of plasmid loss of centromeric plasmids in *H. uvarum*. The episomal *CEN* phenotypic effect may be the reason centromeres from *H. uvarum* were not isolated from episomal genomic libraries.[5] Episomal-*CEN* DNA influencing cellular physiology is not without precedent in the yeasts, as episomal-*CEN* DNA induces pseudohyphal cell growth in *Candida maltosa*[6]. However, the molecular mechanism(s) by which episomal-*CEN* DNA can induce these cellular state changes remain to be investigated.

**Supplementary Note 4** Ancestral gene linkage analysis

From the perspective of *H. uvarum*, 3/7 of its centromeres were fully contained within a single ancestral linkage group (ALGs *CEN2*, *CEN3*, *CEN7*). For example, *CEN2*-ALG contains *CEN4* from *H. uvarum*, which is linked between the genes *MET14* and *VPS1* (Figure 4b), and this linkage is preserved in the majority of species examined (Figure S5b). We therefore conclude that the ancestral *CEN2* was most likely linked between the *MET14* and *VPS1* genes in the common ancestor of Saccharomycodales and Saccharomycetales. The other centromeres (from *H. uvarum*'s perspective 4/7) were mapped between 2-4 non-contiguous flanking ALGs. Given that our approach is stringent (using only protein annotations) and conservative (e.g., one-to-one reciprocal blast hits), many ALGs can be artifactually fragmented due to species-specific genome arrangements or gene annotation inaccuracies. For example, the *CEN4*-ALGs comprise two separate linkage groups flanking either side of *CEN2* in *H. uvarum*, which is linked between the genes *ERG26* and *RPN9* (Figure S5c). However, in the rest of the species examined, extant ancestral *CEN4* sequences are linked between the genes *ERG26* and *EFM5*, and the ALG containing *RPN9* is not linked to any centromere (Figure S5c). Complicating the matter further is the fact that the *EFM5* gene was not included in our annotations of *H. uvarum* and was involved in an inter-centromere translocation event in *K. lactis* (Figure S6). Thus, *EFM5* was overlooked in the ALG reconstructions, thereby resulting in *CEN4*-ALG being split into two groups. Additionally, we note that the centromere of the *CEN8*-ALG is not conserved in Saccharomycodales, although the local gene order is partially preserved. This could be explained by centromere loss in the Saccharomycodales or centromere gain in the Saccharomycetales (Figure S5g). These caveats notwithstanding, our ALGs reconstructions support the evolutionary relatedness between extant point and point-like centromeres and their common origin (Figure S5d–g, S6a–d, Table 1).

**Supplementary Note 5** *CTF13* homolog identification

The hallmark of Saccharomycetales species with canonical point centromeres is the presence of the yeast-specific CBF3 complex (CBF3c; Cep3, Ndc10, Ctf13, and Skp1), which binds to CDEIII. Two subunits of the CBF3c, Skp1 and Cep3, have homologs outside of the Saccharomycotina yeasts, indicating that their origins predate the emergence of point centromeres.[7] However, the precise evolutionary origin of the CBF3c remains enigmatic as no other order of yeast has been shown to encode the subunits Ctf13 and Ndc10. The absence of Ndc10 and Ctf13 subunits in other yeast species was previously taken as evidence that these two proteins, and the origins of the CBF3c, are descended from 2μ plasmid's DNA-adaptor proteins Rep1 and Rep2[7]. Furthermore, in this model, the point centromeric DNA itself is descended from the 2μ's STB locus, which is bound by REP1/2. Nevertheless, recent structural insights have shown that Ctf13 is divergent F-box containing leucine rich repeat protein that bridges the Skp1 subunit to a Cep3 dimer.[8,9] In addition, the subunit Ndc10 bares structurally similarities to tyrosine recombinases[10], therefore the claim that Ndc10 and Ctf13 are of 2μ origin is not clear.

We therefore investigated this hypothesis by examining the presence and absence of Ndc10 and Ctf13 across all 1,154 sequenced budding yeasts genomes from a recent comprehensive study[11]. We were surprised to find that Ndc10 is broadly conserved across the yeast subphylum and searches of the InterPro database revealed proteins with similar domain structure in distantly related fungal lineages (Figure 4c; Figure S7a–b, Table S4). For Ctf13, we observed that high-confidence matches were only found in species from the Saccharomycetales and, surprisingly, we also found that species from the Saccharomycodales have low-scoring matches to Ctf13 (Figure 4c). Additionally, we find evidence that Saccharomycodales and Saccharomycetales Ctf13 are evolutionary related to two conserved yeast F-box containing leucine rich repeat proteins, Das2 and Ydr131c. Phylogenetic analysis supported the conclusion that *CTF13* is a member of the broader *DAS1*/*YDR131C* family (Figure S9a–b). These insights suggests that *CTF13* is an ancient paralog of *DAS1/YDR131C*, which underwent gene duplication deep in the evolutionary history of the Saccharomycotina yeast subphylum (Figure 4c; Figure S9).

### **Supplementary Note 6** Ty5 cluster analysis

Ty5 clusters are thought to function as centromeres in diverse yeast species across multiple orders, including Pichiales (*Ogataea polymorpha*), Serinales (*Scheffersomyces stipitis*, *Debaryomyces hansenii*), and Ascoideales (*Saccharomycopsis fibuligera*).[12–14] We found that Ty5 LTR retrotransposons are relatively uncommon in Saccharomycotina yeast (present in ~37% of species examined), though they appear in all major orders (Figure S10a). However, we cannot rule out the possibility that Ty5 elements are common and strain-to-strain variation explains our observations. We classified species by Ty5 distribution: (1) telomeric enrichment (5/30 species, as in *Saccharomyces*), (2) single-locus per chromosome enrichment (11/30 species, as in Ty5 cluster centromeres), and (3) no specific enrichment (14/30 species). Notably, two species from Phaffomycetales, *Cyberlindnera sargentensis* and *Cyberlindnera fabanii*, the sister order to Saccharomycodales and Saccharomycetales, have small, single-per-chromosome Ty5 clusters (Figure S10). Additionally, the Pichiales species *Pichia terricola* and two other *Saccharomycopsis* species also exhibit this pattern (Figure S10). We propose that these Ty5 clusters represent their centromeric regions (Table S8).

### **Supplementary Note 7** Key DNA constructs

Cse4(1-60)::mNeonGreen::Cse4(61-222)-HphMX

Homology Arms Cse4 coding sequence mNeonGreen Linkers HphMX

ATAAAAATGAATGTATATAAAATATAGGGTTTTCTTTGGGCTGTTAATTACTTAGATTAAATAAATATTATACAGCAAAAATTTAAAAAAAACAGAAAAA
AAGAATCGCATATTCTGCGTTAAGCAGCGCAAAAAAAAAAAAAGTAAGGGATAAATTAGAAAAGATTGAGGATATAGCTAATGATAAGATCAGAGGGTATA
TGTAACGGAGATAAGACAACTATACATATGAATATCCAATTCAAGCAACGTTATTCACATATTGAAGAAACAGCAATGATGAACAATTGATTCACTATTT
CAAAATGGTGATCCTATTTAGTTTTTTTTTTTGTTTTTCATAATTGAAATAGCTGAAACGTCTTTTTATCAGACGTTTAATTCACTGGTGAACACAAACA
TCTTGACAAGAAAAAAACTTAAAACTATCTTAAGCAATCCTACCGCTATGTTGTAACCACAATTTTTTACTTTCAAAGTGCTAAAATACGGTTGATTTTG
CTTAGTGGTGCTTACTTTCAATGATGATCACGTGGTTGCTTTTCACACGTTCCTGATATTTCTTTTTTTTCTTTTTTTTTCGCGTTAAGCATTGAACTTCTA
AGGTCTAGTTGAGCGACTTTTGTTTATTTAGATTATTTTCTTTTTTAAATCTATAATCCCAAGGAAATCATACACACCTCCACCTAAACACATACTCTCA
TCAAACGAAGAAAAAAAAAAAAAAATTAAAAATGGAACAAGAACATGATTTTAATGATTTGAACAAAAGGGCTTTAAATTTACTAGGACAGAACGATAGTC
TGATAAGTGACCGTAAATCCTTAATTTCTTCTGAAAAAAACAAGCATCATGAAGGTATGATGGACTATCTAAAAAGACAAAGATTTAAGATCAACAGAAA
CAGAGCTGTTTCCTGCAGCCCGGGGGATCCACCGGTCGCCACCGTGAGCAAGGGCGAGGAGGATAACATGGCCTCTCTCCCAGCGACACATGAGTTACAC
ATCTTTGGCTCCATCAACGGTGTGGACTTTGACATGGTGGGTCAGGGCACCGGCAATCCAAATGATGGTTATGAGGAGTTAAACCTGAAGTCCACCAAGG
GTGACCTCCAGTTCTCCCCCTGGATTCTGGTCCCTCATATCGGGTATGGCTTCCATCAGTACCTGCCCTACCCTGACGGGATGTCGCCTTTCCAGGCCGC

CATGGTAGATGGCTCCGGCTACCAAGTCCATCGCACAATGCAGTTTGAAGATGGTGCCTCCCTTACTGTTAACTACCGCTACACCTACGAGGGAAGCCAC
ATCAAAGGAGAGGCCCAGGTGAAGGGGACTGGTTTCCCTGCTGACGGTCCTGTGATGACCAACTCGCTGACCGCTGCGGACTGGTGCAGGTCGAAGAAGA
CTTACCCCAACGACAAAACCATCATCAGTACCTTTAAGTGGAGTTACACCACTGGAAATGGCAAGCGCTACCGGAGCACTGCGCGGACCACCTACACCTT
TGCCAAGCCAATGGCGGCTAACTATCTGAAGAACCAGCCGATGTACGTGTTCCGTAAGACGGAGCTCAAGCACTCCAAGACCGAGCTCAACTTCAAGGAG
TGGCAAAAGGCCTTTACCGATGTGATGGGCATGGACGAGCTGTACAAGTTACCGCGGCCGCCCACCGCTCTAGAAGTTGATACTCTATCCAATCAAATAT
CCCAAAAAACAGGGCTTAACCTTAATTCCAAAAAATTATACTCGGTTAGAAGGAAAAAAAGAAGATCAAGTCAATACTCACCTTCTTCAAGCAGAAGAAA
CTCCAAGTATGGCAACAGCAGAGAATCCAGCATATATGAAAAAAAATCACATGCATCTGATATGGGTGCTTCAGACAAAGCATTGCGTGAAATCATGAAA
TATCAATCAACAACGGATTTGCTTGTGGCCAAAATTCCCTTTGCAAAAATAGTCAAACAAATTACAGATAGATATACCATAGCCGACGGGTCTTCGGAAC
CTTACAAATGGCAGAGTATGGCTTTGTTGGCTTTGCAAGAGGCAAGTGAAGCTTATATAGTTGGTCTTTTAGAGCACACAAATTTATTAGCTATACACGC
TAAAAGAAGTACGGTTATGAAAAAAGACTTGCAATTGGCAAGAAGAATTAGAGGTTTTCAATAGTATATATGAAGTTACCCGTTTAGGAAATACCAACAA
TTTCATCACACAAGCCCTGAAAAAAAAAAAAAATATTCATTCTTTTTTTTTCTCCTATGATCGCTTTAGGAAAAAAATCAAAACTTCTACCTAATTTTAA
ACAGTTCATAAGGGGTAAACGAAAACAAAAAATGCAAGCAAAATGATGCAATATAAGCGGTTATTCATTCTTCTCCTAAACAAGCCACTTCTGCGTACTT
TTCTAAGAACTTCTTATCTTCTCCACTTGGCCCTCTCGTTTCTTTTTCTAATTTCTATCATCAACACTAAAATTTGTCTAGCAATTGCTAGAAATCAAGG
TGGCGAAAATTTTTTTTCATTTTCTTTTTCAATGAATTGCTCATTAGGTTGTCGATATAAAAGAAAAAGGAAGAAAATAGAAATATTTCATTTGTGTCAG
ATGTTTTACTTTCATCAGGGTTTTTCTTAAACATTATGAAGTTCTTGTTTAATTTTCTTATTGATTACTAGAGAGTAAGAAAAAACCAGTCATTAACAAT
AAATTTAAAAAAATGGGTAAAAAGCCTGAACTCACCGCGACGTCTGTCGAGAAGTTTCTGATCGAAAAGTTCGACAGCGTCTCCGACCTGATGCAGCTCT
CGGAGGGCGAAGAATCTCGTGCTTTCAGCTTCGATGTAGGAGGGCGTGGATATGTCCTGCGGGTAAATAGCTGCGCCGATGGTTTCTACAAAGATCGTTA
TGTTTATCGGCACTTTGCATCGGCCGCGCTCCCGATTCCGGAAGTGCTTGACATTGGGGAATTCAGCGAGAGCCTGACCTATTGCATCTCCCGCCGTGCA
CAGGGTGTCACGTTGCAAGACCTGCCTGAAACCGAACTGCCCGCTGTTCTGCAGCCGGTCGCGGAGGCCATGGATGCGATCGCTGCGGCCGATCTTAGCC
AGACGAGCGGGTTCGGCCCATTCGGACCGCAAGGAATCGGTCAATACACTACATGGCGTGATTTCATATGCGCGATTGCTGATCCCCATGTGTATCACTG
GCAAACTGTGATGGACGACACCGTCAGTGCGTCCGTCGCGCAGGCTCTCGATGAGCTGATGCTTTGGGCCGAGGACTGCCCCGAAGTCCGGCACCTCATG
CACGCGGATTTCGGCTCCAACAATGTCCTGACGGACAATGGCCGCATAACAGCGGTCATTGACTGGAGCGAGGCGATGTTCGGGGATTCCCAATACGAGG
TCGCCAACATCTTCTTCTGGAGGCCGTGGTTGGCTTGTATGGAGCAGCAGACGCGCTACTTCGAGCGGAGGCATCCGGAGCTTGCAGGATCGCCGCGGCT
CCGGGCGTATATGCTCCGCATTGGTCTTGACCAACTCTATCAGAGCTTGGTTGACGGCAATTTCGATGATGCAGCTTGGGCGCAGGGTCGATGCGACGCA
ATCGTCCGATCCGGAGCCGGGACTGTCGGGCGTACACAAATCGCCCGCAGAAGCGCGGCCGTCTGGACCGATGGCTGTGTAGAAGTACTCGCCGATAGTG
GAAACCGACGCCCCAGCACTCGTCCGAGGGCAAAGGAATAAATTTGACTTTTAATATAAAAAAATCTTTTTCTATTCAATAAATGTCATTTACAAACTCT
ATGCTAGTAAACCCAGTATAAAATATATAATCCTATATAATTATTTATATGATTAAAAATTAAAAAAAAAAAAAAATAAAACCAAAAAAAAAAAAACACAA
TCGATTATAGCTTTTTGTTTTCGTTTTAGATATCTTGAAGTGCAATGTTGTTTAAACTCTGTTATTAGAGTTGTTATAGTATTGATTACGTTAAATGTTA
ATATATAAATATTTTTAAGTATTATGGTTACATTAGTAGATTTGTCTTCTTTCTTTTTTTTTTTTTAACGTTGATGGGGCACAAATACTTTCAAAGATTTG
ATTGTGGATTGAGATGGCTCATCTAATGTTTGCATTTTGCCCAAATCGTTCAAAATGGATTTTCTTTTATACTGCCTTTCTTTCATTTTCATGACAGCAT
TTGTAAGCATAGTCAAGTCACCTTCTTCAGAATCGTCATCTTGAGGGTCAATAGCAGGTTTATTGCTTGTGTGGATCATTTCTGATTTTTCATATGATAT
TTCATCATCTTCAATATCATCTTTGACATAGCTTTGTGGTAAATCATCTGATATATTTGCAGCTTTCAATTCTTGTGTTGTTTTTTTTACTGTAACATCT
ATAGAAGAATTTTCACTGACTCGATCGTCGTCTTGTTCTTCTTCAGACCAAAATAAATTCTTTTCTCTTTCAGGAACTTCTGGAGCCTTGAGTTTTTGTT
GACCAATTTTTTGTTTTCTGTTTCTCCAATTTACACCTTCTATGCTTTCAACACTCTTCATCATCAATCTCATTTGTTCTTTATGTGACAAGTTTTCACC
ATCACTGGTATTCATCGATCTATGTCCACTAGATTTAAATGAGTTTTTATCTGTGTGTTCAATCGAGCTTAAATGACTGTCATCGTTTAAAATTGCATGG
GAAGATAAAGAACTGTATCTCATTGGGCCCAAATGTACTTCTTCAAGACCGCTGGGTATATCAAACTCTGATAAGTTACCTAATGTGACTTTGCTTTGGT
TAAATTGAGAAGGATTGTTTAATGCAGGATTTTTTAATGGCTGTTTACTCATGTATGTAGTATTATTTGTTATGTTTGGGTTTTTAAAGGAGGTTAGATC
ATCATAGTCAAGACCAGTTTCTTCATAACTTTCATCATGAGTTGTTTGGTTATTAAATCCCATGGTAATGTGCTTGTCAATGTAAT

pJJ3252-*CEN4*

CDEII Nearest CAM Plasmid backbone

GATCTACTAGTCATATGGCATGCCTGCAGGTCGACTCTAGAGGATCCCCGCTAAGCTTGTGATATACATAAGCCACAAGACTTGAATTAAAAAAAAAAAA
GAATCTCATAATGTGCAATTTATTATATGTAACAAGTGAATCTCTCATGTGAATTACTGTTTCAATGCTTTGATGGACTTTAGTGTTAACACATAAGTTG
CAACAATACTTCTATAAATAACCACCGCTACCAAAATTTAATACAAAATCACATTAAGAAGATCTTTCAGAAAAAATCAGTTTCATTTTGATCACAATAA
TTTTTTGCAAAGCATTTTCTGCTTTAGATGCAACTTTTTATGAAAACATGCAAAAAAAATTGTCAAATACTAAGCTTATGCGCAATATATGCGTTAATTCA
GCTTTTAACGACATAAAACTTGCCAATAAAAATTGTCCTTGCAAGCATATACTGCAATAAAACCTAATAAAAACATTTAAAAAAAAAAATTAAAAAAAAA
GCAAAGCCAACTAAGTTTACATTTGCATTAATTAATATACATACACACATATAGCAAAATATAGTAATGTTTAGTTTGTAAACAAGATCAAAAGAAATAA
TTTATCAACAACTATATATCAATTTTATAACCGTTTTGATGTCATCAAGTTGGTTGAACTTTACAATTGTGTTTACGTTTAGCACTATAAAATATATATA
TGACATTCAAAATGTACCAACAACAAAACTAT**GCG**TTAAAA**ATTTAAATTAAATTAATTTTCACAATATATTTATATATTTTTTTAATATAATAAAACAC**
**ACAAACGTATTATTTTTATTTAAACATTTTTAATAAGGTAAAAATTGTATATAAAATATTGTTTATTTTTTTTGCAAAACAATTATTTTTAATTTAGTTA**

GACGCATATTTTTGAAAATATTGTAACACCAAGTATTACACCTAACGCTTATAAATATTTTTACACGCAATAAATGCGTTGACATTGTTAAAAAATACTT
TCAAGCGCAAATGTGCAATGTTTTAACGCAGAATATATACTGTGTTTCATTGTAAATTTCAAAAAGTTGTTGTTTTACGACGAAACCATAAAGAATTGCT
AAGGTTTTTGATATCTTAGGATATTATGGTTGTTTTAAAATTATATAGACAATTTTGAACAATTATAAGCGTAATAGTTTTACTTTTGGTGTTTTTCATC
TTTATGCCAGAAAATCATATAATTTAATAGTTTTTTGAAGAAAAATTAAAAAAATTCGGATATAATATTTTAATTGTGAATAAACAGTCATGGCGAACTT
ATACATCAAAGCTACAGTACAATGCAACAAAAGAAAAATACATGGCCATTACATGCTTGAATAAGCGTTTTAGTATTATTAATCACTTTTGTTTTCAGAT
AACATTTAAAATTGATATATAAATTAAGTTATTGTAATGTGACTATGAAAAAAATAAATTTCTAGTATACTTAAATAGTCATCAGTCAATAAAACAAAGA
AATCTCATTTGAAATGTCAAAATTCAGAAACAAATCGGTTAAAAATGAGTGGCAACCAAATGAACCACTAAAGAATTATAAAATTTTTTTGACGAGAAGA
AGCAATAACAATTGGCCAGACAAGCTTGTGTAGTAAACCACTTCCTTAAACTTAAATTACTCAATACACATCTTAAAAGTGGTGGTGGGTGAGCTCGAAT
TCACTGGCCGTCGTTTTACAACGTCGTGACTGGGAAA

pJJ3252-*CEN4*-Δleft

CDEII Nearest CAM Plasmid backbone

GATCTACTAGTCATATGGCATGCCTGCAGGTCGACTCTAGAGGATCCCCGCAAGATCAAAAGAAATAATTTATCAACAACTATATATCAATTTTATAACC
GTTTTGATGTCATCAAGTTGGTTGAACTTTACAATTGTGTTTACGTTTAGCACTATAAAATATATATATGACATTCAAAATGTACCAACAACAAAACTAT
GCGTTAAAAATTTAAATTAAATTAATTTTCACAATATATTTATATATTTTTTTAATATAATAAAACACACAAACGTATTATTTTTATTTAAACATTTTTA
ATAAGGTAAAAATTGTATATAAAATATTGTTTATTTTTTTTGCAAAACAATTATTTTTAATTTAGTTAGACGCATATTTTTGAAAATATTGTAACACCAA
GTATTACACCTAACGCTTATAAATATTTTTACACGCAATAAATGCGTTGACATTGTTAAAAAATACTTTCAAGCGCAAATGTGCAATGTTTTAACGCAGA
ATATATACTGTGTTTCATTGTAAATTTCAAAAAGTTGTTGTTTTACGACGAAACCATAAAGAATTGCTAAGGTTTTTGATATCTTAGGATATTATGGTTG
TTTTAAAATTATATAGACAATTTTGAACAATTATAAGCGTAATAGTTTTACTTTTGGTGTTTTTCATCTTTATGCCAGAAAATCATATAATTTAATAGTT
TTTTGAAGAAAAATTAAAAAAATTCGGATATAATATTTTAATTGTGAATAAACAGTCATGGCGAACTTATACATCAAAGCTACAGTACAATGCAACAAAA
GAAAAATACATGGCCATTACATGCTTGAATAAGCGTTTTAGTATTATTAATCACTTTTGTTTTCAGATAACATTTAAAATTGATATATAAATTAAGTTAT
TGTAATGTGACTATGAAAAAAATAAATTTCTAGTATACTTAAATAGTCATCAGTCAATAAAACAAAGAAATCTCATTTGAAATGTCAAAATTCAGAAACA
AATCGGTTAAAAATGAGTGGCAACCAAATGAACCACTAAAGAATTATAAAATTTTTTTGACGAGAAGAAGCAATAACAATTGGCCAGACAAGCTTGTGTA
GTAAACCACTTCCTTAAACTTAAATTACTCAATACACATCTTAAAAGTGGTGGTGGGTGAGCTCGAATTCACTGGCCGTCGTTTTACAACGTCGTGACTG
GGAAA

pJJ3252-*CEN4*-Δright

CDEII Nearest CAM Plasmid backbone

GATCTACTAGTCATATGGCATGCCTGCAGGTCGACTCTAGAGGATCCCCGCTAAGCTTGTGATATACATAAGCCACAAGACTTGAATTAAAAAAAAAAAA
GAATCTCATAATGTGCAATTTATTATATGTAACAAGTGAATCTCTCATGTGAATTACTGTTTCAATGCTTTGATGGACTTTAGTGTTAACACATAAGTTG
CAACAATACTTCTATAAATAACCACCGCTACCAAAATTTAATACAAAATCACATTAAGAAGATCTTTCAGAAAAAATCAGTTTCATTTTGATCACAATAA
TTTTTTGCAAAGCATTTTCTGCTTTAGATGCAACTTTTTATGAAAACATGCAAAAAAATTGTCAAATACTAAGCTTATGCGCAATATATGCGTTAATTCA
GCTTTTAACGACATAAAACTTGCCAATAAAAATTGTCCTTGCAAGCATATACTGCAATAAAACCTAATAAAAACATTTAAAAAAAAAAATTAAAAAAAAA
GCAAAGCCAACTAAGTTTACATTTGCATTAATTAATATACATACACACATATAGCAAAATATAGTAATGTTTAGTTTGTAAACAAGATCAAAAGAAATAA
TTTATCAACAACTATATATCAATTTTATAACCGTTTTGATGTCATCAAGTTGGTTGAACTTTACAATTGTGTTTACGTTTAGCACTATAAAATATATATA
TGACATTCAAAATGTACCAACAACAAAACTATGCGTTAAAAATTTAAATTAAATTAATTTTCACAATATATTTATATATTTTTTTAATATAATAAAACAC
ACAAACGTATTATTTTTATTTAAACATTTTTAATAAGGTAAAAATTGTATATAAAATATTGTTTATTTTTTTTGCAAAACAATTATTTTTAATTTAGTTA
GACGCATATTTTTGAAAATATTGTAACACCAAGTATTACACCTAACGCTTATAAATATTTTTACACGCAATAAATGCGTTGACATTGTTAAAAAATACTT
TCAAGCGCAAATGTGCAATGTTTTAACGCAGAGGTGAGCTCGAATTCACTGGCCGTCGTTTTACAACGTCGTGACTGGGAAA

pJJ3252-*CEN4*-Δleft-right (m*CEN4*)

CDEII Nearest CAM Plasmid backbone

GATCTACTAGTCATATGGCATGCCTGCAGGTCGACTCTAGAGGATCCCCGCAAGATCAAAAGAAATAATTTATCAACAACTATATATCAATTTTATAACC
GTTTTGATGTCATCAAGTTGGTTGAACTTTACAATTGTGTTTACGTTTAGCACTATAAAATATATATATGACATTCAAAATGTACCAACAACAAAACTAT
**GCG**TTAAAA**ATTTAAATTAAATTAATTTTCACAATATATTTATATATTTTTTTAATATAATAAAACACACAAACGTATTATTTTTATTTAAACATTTTTA**
**ATAAGGTAAAAATTGTATATAAAATATTGTTTATTTTTTTTGCAAAACAATTATTTTTAATTTAGTTAGACGCATATTTTTGAAAATATTGTAACACCAA**
**GTATTACACCTAACGCTTATAAATATT**TTTACA**CGC**AATAAAT**GCG**TTGACATTGTTAAAAAATACTTTCAAGCGCAAATGTGCAATGTTTTAACGCAGA
GGTGAGCTCGAATTCACTGGCCGTCGTTTTACAACGTCGTGACTGGGAAA

### pJJ3252-*CEN4*-ΔCDEII

CDEII Nearest CAM Plasmid backbone

GATCTACTAGTCATATGGCATGCCTGCAGGTCGACTCTAGAGGATCCCCGCTAAGCTTGTGATATACATAAGCCACAAGACTTGAATTAAAAAAAAAAAA
GAATCTCATAATGTGCAATTTATTATATGTAACAAGTGAATCTCTCATGTGAATTACTGTTTCAATGCTTTGATGGACTTTAGTGTTAACACATAAGTTG
CAACAATACTTCTATAAATAACCACCGCTACCAAAATTTAATACAAAATCACATTAAGAAGATCTTTCAGAAAAAATCAGTTTCATTTTGATCACAATAA
TTTTTTGCAAAGCATTTTCTGCTTTAGATGCAACTTTTTATGAAAACATGCAAAAAAATTGTCAAATACTAAGCTTATGCGCAATATATGCGTTAATTCA
GCTTTTAACGACATAAAACTTGCCAATAAAAATTGTCCTTGCAAGCATATACTGCAATAAAACCTAATAAAAACATTTAAAAAAAAAAATTAAAAAAAAA
GCAAAGCCAACTAAGTTTACATTTGCATTAATTAATATACATACACACATATAGCAAAATATAGTAATGTTTAGTTTGTAAACAAGATCAAAAGAAATAA
TTTATCAACAACTATATATCAATTTTATAACCGTTTTGATGTCATCAAGTTGGTTGAACTTTACAATTGTGTTTACGTTTAGCACTATAAAATATATATA
TGACATTCAAAATGTACCAACAACAAAACTAT**GCG**TTAAAATTACA**CGC**AATAAAT**GCG**TTGACATTGTTAAAAAATACTTTCAAGCGCAAATGTGCAAT
GTTTTAACGCAGAATATATACTGTGTTTCATTGTAAATTTCAAAAAGTTGTTGTTTTACGACGAAACCATAAAGAATTGCTAAGGTTTTTGATATCTTAG
GATATTATGGTTGTTTTAAAATTATATAGACAATTTTGAACAATTATAAGCGTAATAGTTTTACTTTTGGTGTTTTTCATCTTTATGCCAGAAAATCATA
TAATTTAATAGTTTTTTGAAGAAAAATTAAAAAAATTCGGATATAATATTTTAATTGTGAATAAACAGTCATGGCGAACTTATACATCAAAGCTACAGTA
CAATGCAACAAAAGAAAAATACATGGCCATTACATGCTTGAATAAGCGTTTTAGTATTATTAATCACTTTTGTTTTCAGATAACATTTAAAATTGATATA
TAAATTAAGTTATTGTAATGTGACTATGAAAAAAATAAATTTCTAGTATACTTAAATAGTCATCAGTCAATAAAACAAAGAAATCTCATTTGAAATGTCA
AAATTCAGAAACAAATCGGTTAAAAATGAGTGGCAACCAAATGAACCACTAAAGAATTATAAAATTTTTTTGACGAGAAGAAGCAATAACAATTGGCCAG
ACAAGCTTGTGTAGTAAACCACTTCCTTAAACTTAAATTACTCAATACACATCTTAAAAGTGGTGGTGGGTGAGCTCGAATTCACTGGCCGTCGTTTTAC
AACGTCGTGACTGGGAAA

### pJJ3252-m*CEN4*-CDEII-scrambled

CDEII scrambled Nearest CAM Plasmid backbone

GATCTACTAGTCATATGGCATGCCTGCAGGTCGACTCTAGAGGATCCCCGCAAGATCAAAAGAAATAATTTATCAACAACTATATATCAATTTTATAACC
GTTTTGATGTCATCAAGTTGGTTGAACTTTACAATTGTGTTTACGTTTAGCACTATAAAATATATATATGACATTCAAAATGTACCAACAACAAAACTAT
**GCG**TTAAAA**ATTAAATATTTAATAATAATAATGTATGAAATAATTTTCACTTGGTAATTATTCACATTTCTAAAATTTATTTTTCATATCGAATCATATA**
**CATTATAATTTAACTCAGGAAAATACTAATTTATATTTGAAATAACTATATTTAATTTAAATGTGTAATATCTTTATAATATAATTTTAGCTTCTGCCTC**
**TATTTTAAAAGAAATATTATAAAATT**TTTACA**CGC**AATAAAT**GCG**TTGACATTGTTAAAAAATACTTTCAAGCGCAAATGTGCAATGTTTTAACGCAGAG
GTGAGCTCGAATTCACTGGCCGTCGTTTTACAACGTCGTGACTGGGAAA

### pJJ3252-m*CEN4*-CAM-scrambled

CDEII Nearest CAM scrambled Plasmid backbone

GATCTACTAGTCATATGGCATGCCTGCAGGTCGACTCTAGAGGATCCCCGCAAGATCAAAAGAAATAATTTATCAACAACTATATATCAATTTTATAACC
GTTTTGATGTCATCAAGTTGGTTGAACTTTACAATTGTGTTTACGTTTAGCACTATAAAATATATATATGACATTCAAAATGTACCAAAACTAAGACCTA
AAGACATTA**ATTTAAATTAAATTAATTTTCACAATATATTTATATATTTTTTTAATATAATAAAACACACAAACGTATTATTTTTATTTAAACATTTTTA**
**ATAAGGTAAAAATTGTATATAAAATATTGTTTATTTTTTTTGCAAAACAATTATTTTTAATTTAGTTAGACGCATATTTTTGAAAATATTGTAACACCAA**
**GTATTACACCTAACGCTTATAAATATT**TTTACACGCAATAAATGCGTTGACATTGTTAAAAAATACTTTCAAGCGCAAATGTGCAATGTTTTAACGCAGA
GGTGAGCTCGAATTCACTGGCCGTCGTTTTACAACGTCGTGACTGGGAAA

### Supplementary Discussion

The evolutionary transition from ancestral epigenetic centromeres to genetic point centromeres involved complex genome changes (Figure 7a). While the precise macroevolutionary processes remain unclear, our findings reveal a previously unknown evolutionary innovation: point and proto-point centromeres are descended from Ty5 LTR retrotransposons. This unexpected evolutionary link only emerged after a detailed study of centromeres of the Saccharomycodales. The evidence for this conclusion includes: (1) Ty5 elements and LTRs surround *Sa. ludwigii* centromeres, (2) Orthologous centromeric locations and a conserved CBF3 complex, and synteny of genes encoding CBF3 subunits, between Saccharomycodales and Saccharomycetales, (3) point and proto-point centromeres occupy conserved genomic positions where Ty5 clusters are found in diverse orders, such as Phaffomycetales and Serinales, (4) the CDEII sequences of *Sa. ludwigii* centromeres show stromg sequence identity to divergent Ty5 LTRs, and (5) yeast Ty5 LTRs broadly show enrichment with DNA-binding motifs resembling those found in point and proto-point centromeres.

Discovery of Saccharomycodales proto-point centromeres further resolves uncertainties about point centromere evolution. We can infer that the single Cse4-containing structure per chromosome and consequently the single-kinetochore-to-microtubule axis evolved before the strict genetic arrangements of CDE-I, -II, and -III seen in classic point centromeres. Additionally, our findings demonstrate that the proto-point and point centromeres originated from divergent Ty5 LTRs, a state that is exceptionally well preserved in *Sa. ludwigii*. Parsimony would reason that the ancestral centromeres of the Saccharomycodales and Saccharomycetales were sequence-flexible, Ty5 LTR-rich, proto-point centromeres.

Protein mapping associated with epigenetic and genetic centromeres clarified the tempo of point centromere evolution (Figure 7a). Critically we show that Ctf13 and Ndc10 originated from native yeast genes, not the 2μ plasmid.[7] This unambiguously dates the emergence of the CDEIII-binding CBF3c millions of years before the emergence of canonical point centromere DNA – suggesting  the protein coding changes needed to give rise to the CBF3c emerged millions of years before the emergence of the cis-regulatory changes at centromeres. Protein-coding changes evolving before cis-regulatory gains is not without precedent, as has been observed in the evolution of the yeast mating type transcriptional circuit.[15] Innovation of the CBF3c in the ancestor of the Saccharomycetales and Saccharomycodales may have poised the evolution of the strictly structured point centromere. The transition from epigenetic centromere to genetic centromere was thus not a sudden leap as suggested by the 2μ model, but instead a gradual process of Ty5 neocentromere formation and later refinement to a strict genetic specification (Figure 7a).

We propose an alternative hypothesis for point centromere evolution (Figure 7b). In this model, genome-wide relaxation of heterochromatin regulation led to the loss of epigenetic centromeres in early Saccharomycotina evolution. However, selective pressure to regulate transposable elements and centromeric DNA facilitated the co-option of Ty5 regulatory elements at centromeric DNA. Ancient Ty5 trans-regulators, perhaps Cbf1 and Cep3, therefore likely were transcriptional regulators of retrotransposons rather than centromeres. Future evolutionary biochemical and genetic studies will clarify this possibility.

Ty5 elements were originally discovered as inhabitants of telomeres and other silenced regions in *S. paradoxus*.[16] This preference for silent chromatin is dependent on a interaction between its integrase and the Silent Information Regulator 4 protein.[17] Thus, ancient centromeres perhaps were ideal targets for Ty5 integration due to their intrinsic preference for silent chromatin.[18] This contrasts with yeast Ty1-Ty4 retrotransposons, which integrate preferentially near RNA Pol III promoters.[19–21] Centromere targeting may still be retained by the Ty5 elements of *Sa. ludwigii*, as well-conserved, full-length Ty5 elements are found within its centromeric regions, with variation among strains (Table S5). However, the extent to which the Ty5 elements are involved in centromere function is not clear. LTR retrotransposons in eukaryotes are often enriched at centromere sites, suggesting that their role in centromere formation and identity is either deeply conserved or has independently emerged multiple times in evolution. Notably, minor mutations near the integrase domain can shift retrotransposon target specificity, as seen in *Arabidopsis* and *Saccharomyces paradoxus*.[22,23] These findings suggest that LTR retrotransposons target specificity can evolve rapidly while the underlying mechanisms that determine target specificity remain deeply conserved. The small size of *Sa. ludwigii* centromeres and its potential genetic tractability make it a compelling model for future mechanistic studies on retrotransposon function at centromeres.

Our model also predicts that the centromeres of the orders Pichales, Serinales, Alaninales, Ascoideales, Phaffomycetales, Saccharomycetales, and Saccharomycodales are all derived from common ancestral centromeres. In this sense, the evolution of centromeres across these many orders might be thought of a massive natural evolution experiment following heterochromatic centromere loss over millions of years. It is clear many distinct solutions have been found as with many convergent solutions (Figure S1a). It may come to light as more species centromeres are described that point centromeres are no wholely unique and that they may have evolved multiple time independently. Reportly, one species of Serinales, *Candida maltosa* have small centromeres with conserved CDEI-like and CDEII-like sequences[24]. Additionally, as mentioned in supplemental note 3 this species' centromeres also alters its cellular physiology when encoded on episomal centromeres[6]. However, *C. maltosa* does not encode the *CTF13* subunit of the CBF3 complex, thus we expect its reported small centromeres to be of a independent origin to those of the Saccharomycetales and Saccharomycodales, although perhaps also derived from Ty5 LTRs. Nonetheless, small centromeres like *C. maltosa* not only fit our model, but further support its general conclusions since the Ty5 LTR was the element responsible for CDEI, CDEII, and CDEIII.

Ultimately, our work shows that point centromeres emerged through vertical descent, involving key innovations such as the *CTF13* gene duplication. We hypothesize that the many diverse types of centromeres observed in Saccharomycotina yeasts lacking H3K9me and Mis18 are derived from common ancestral Ty5 cluster centromeres. This inference is supported by conserved micro-syntenies near Ty5 clusters from different orders (Figure 6). The evolutionary transition from epigenetic to genetic centromeres represents an extreme example of LTR retrotransposon domestication, as LTRs themselves were repurposed as an essential genetic element responsible for nucleating kinetochore assembly.

**Supplemental Text References**
