## Supplemental Figures for "Ancient co-option of LTR retrotransposons as yeast centromeres"

Supplemental Figure 1. Genome assemblies of Hanseniaspora species.

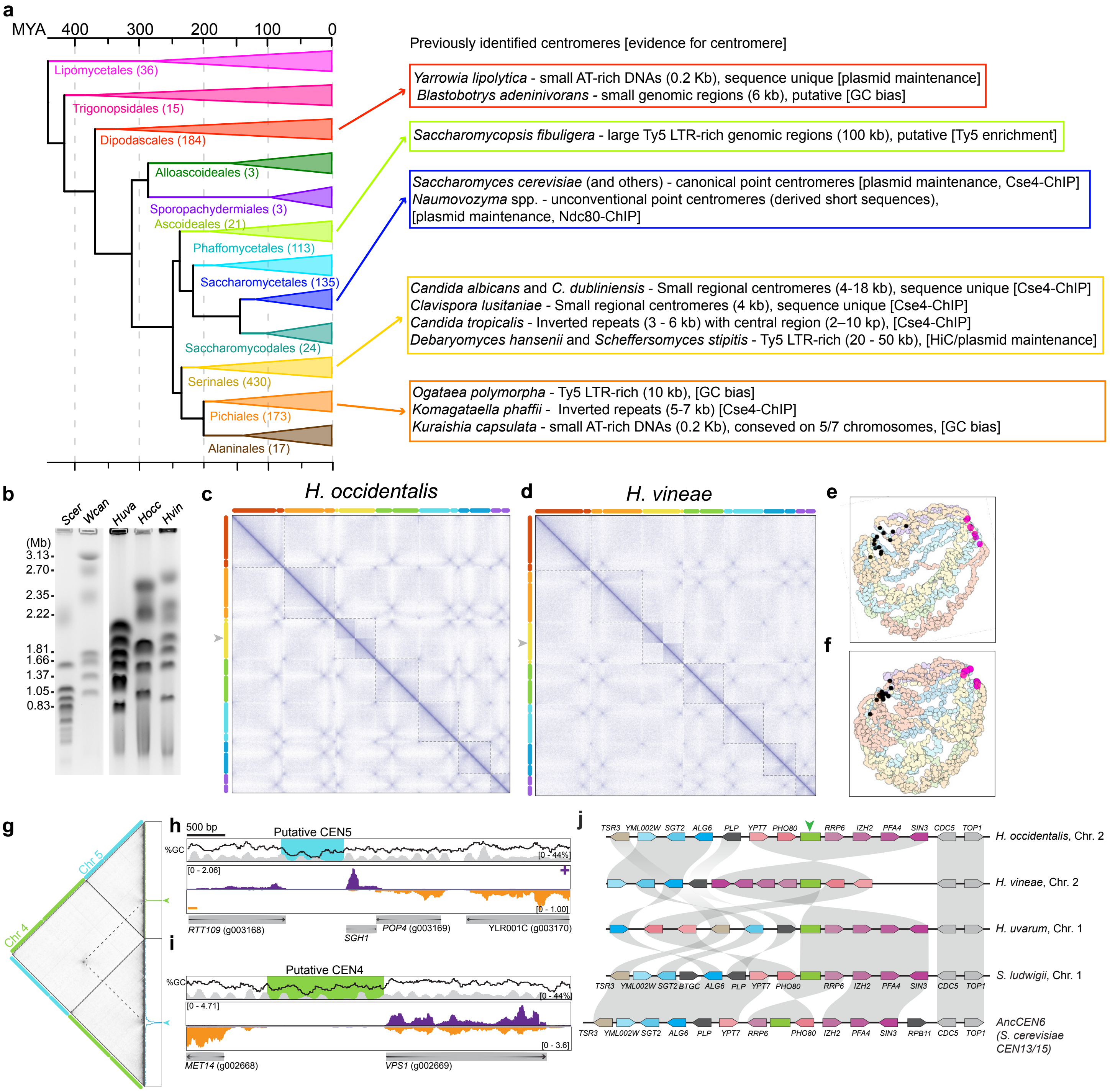

Supplemental Figure 2. Cse4-mNG MNase-ChIP in *H. uvarum*

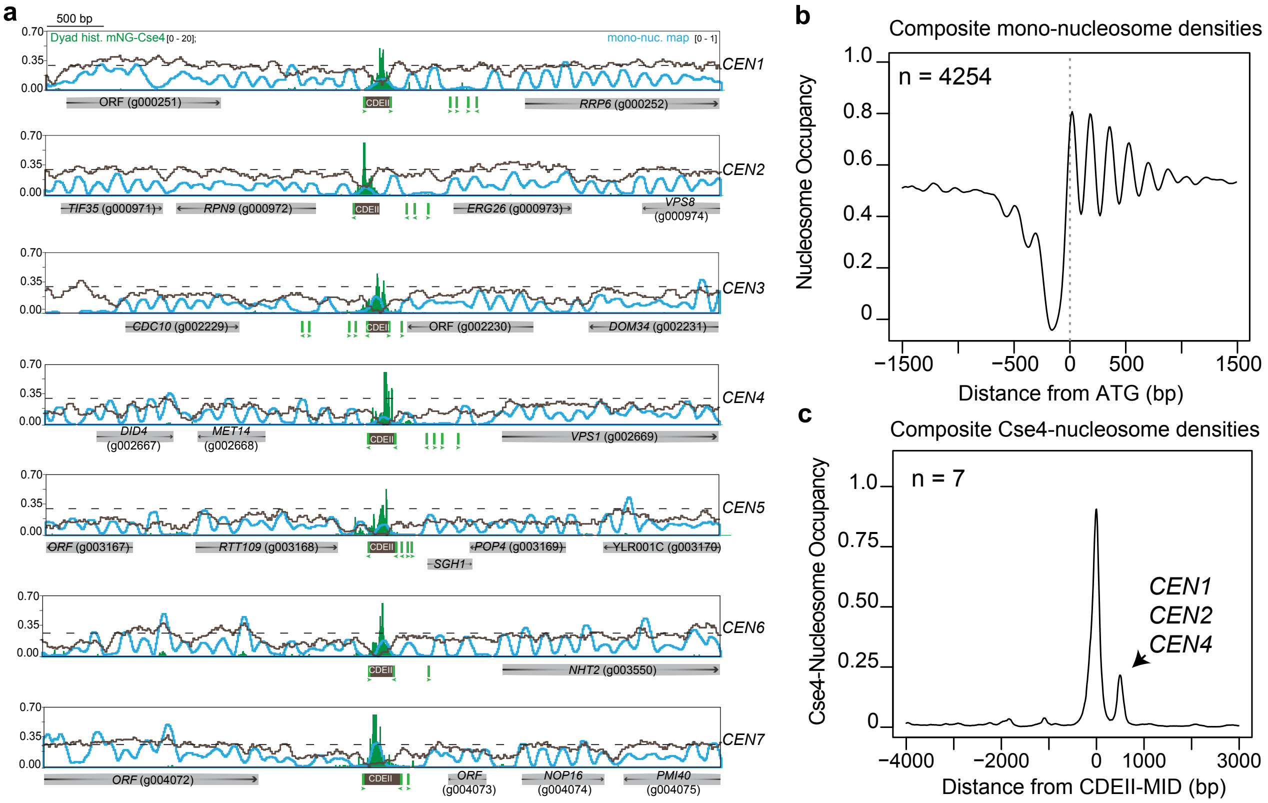

Supplemental Figure 3. Centromeric associated motifs of the Saccharomycodales yeasts

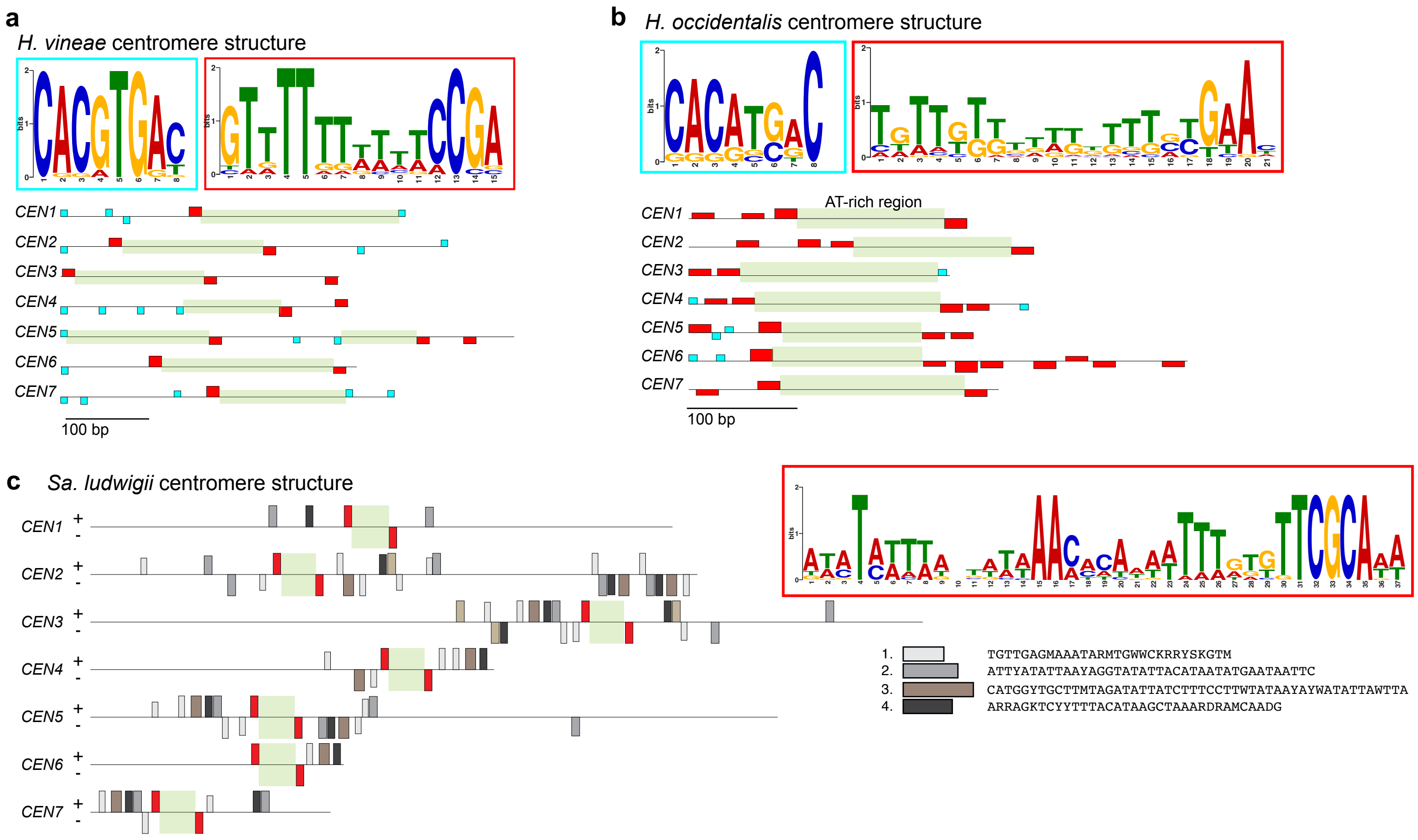

Supplemental Figure 4. CDEII is necessary and sufficient for point-like centromere function

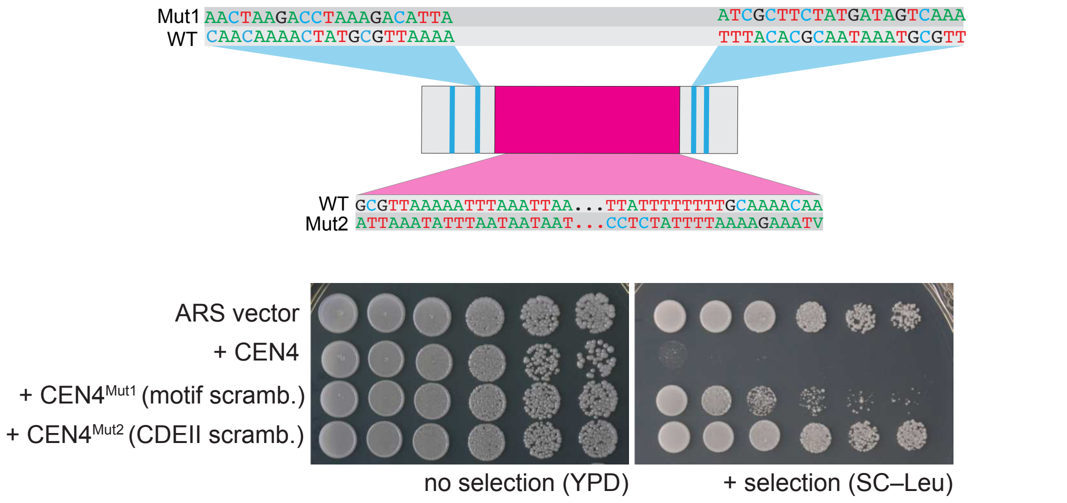

Supplemental Figure 5.

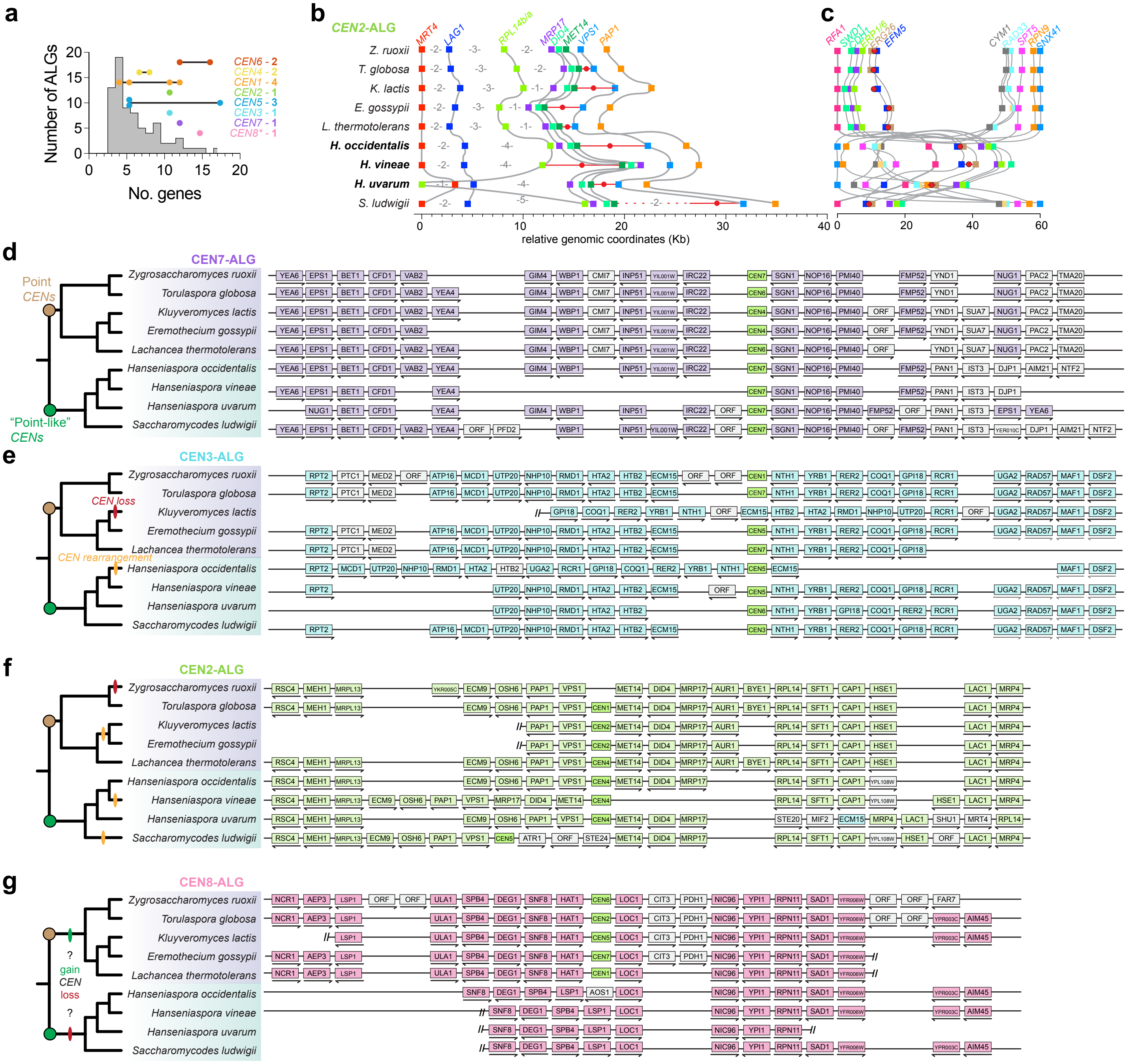

Supplemental Figure 6.

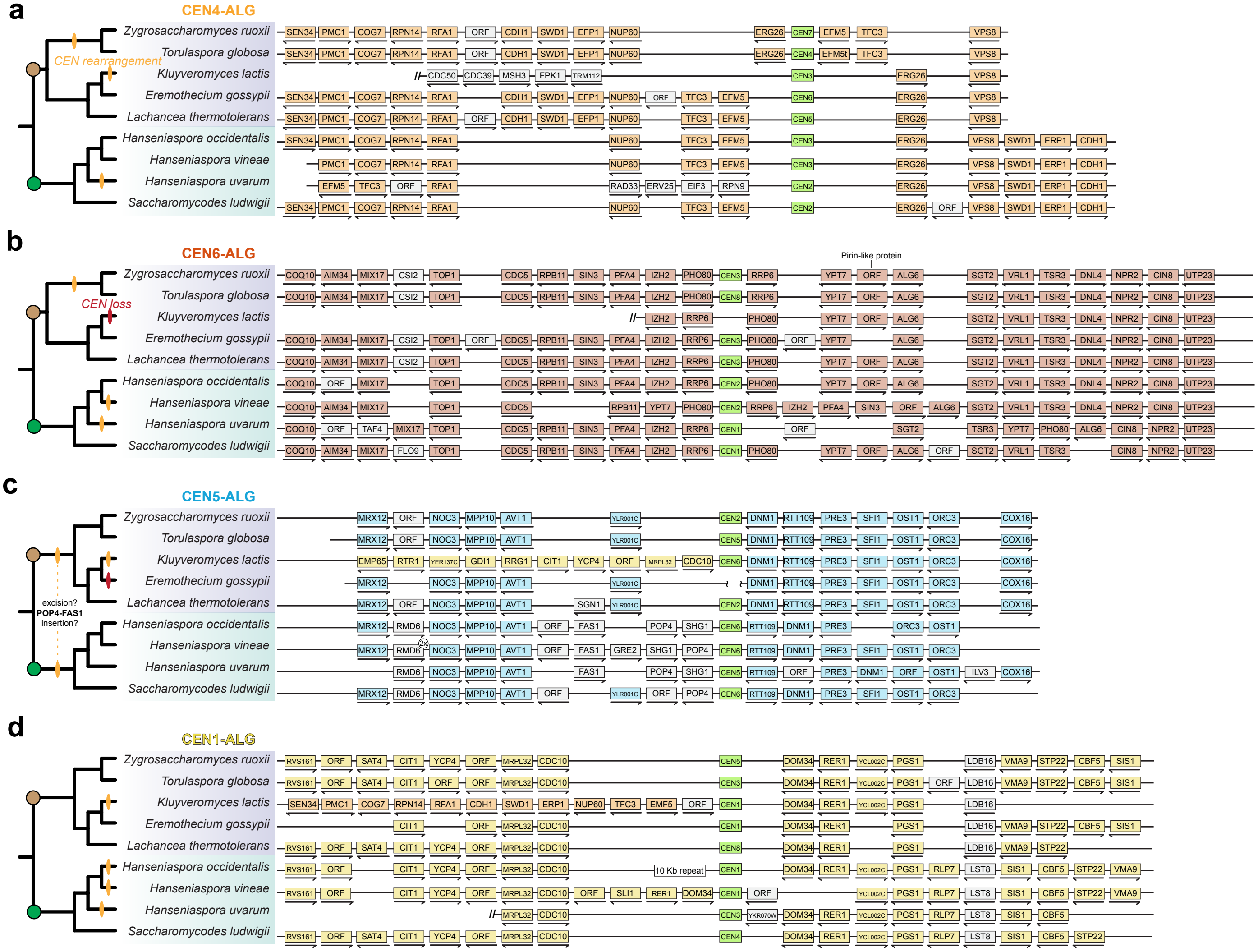

Supplemental Figure 7.

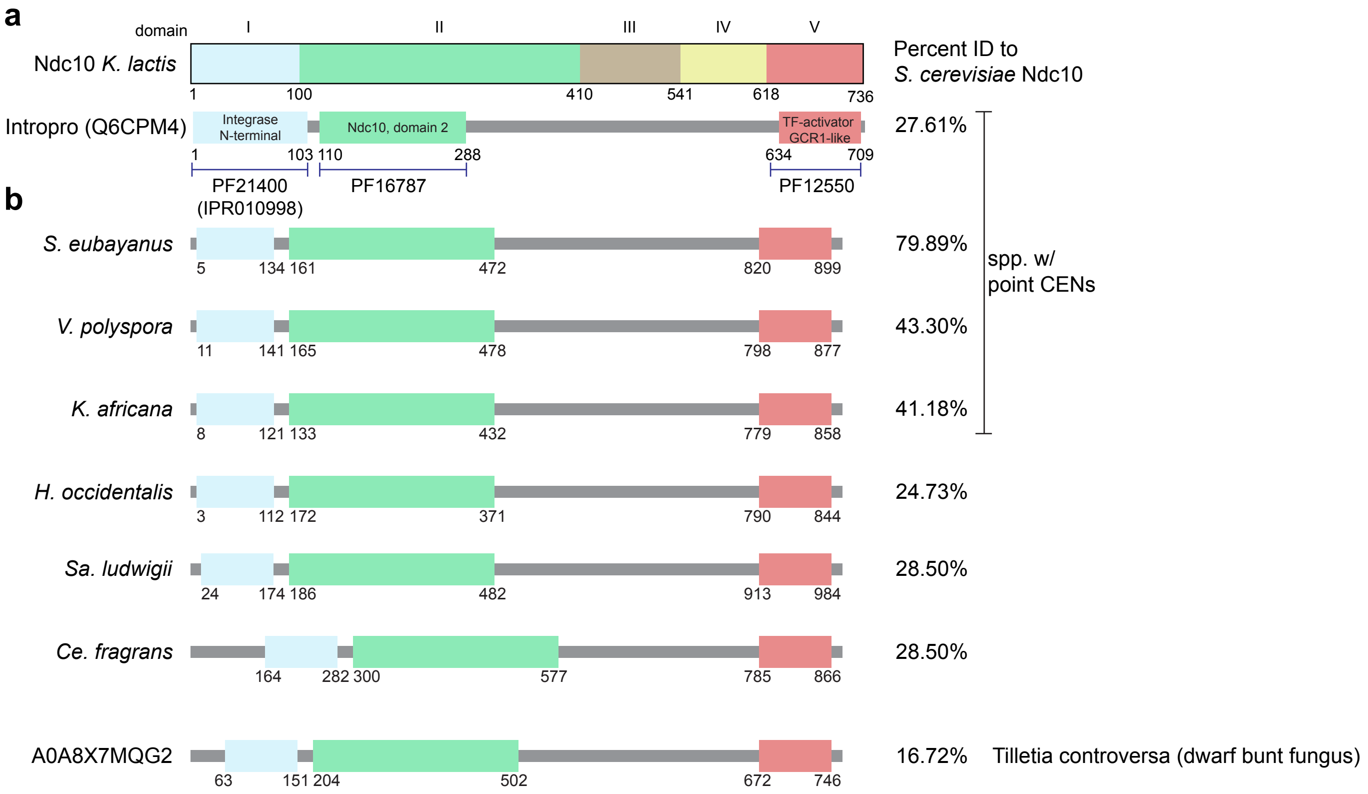

Supplemental Figure 8. Saccharomycodales yeasts have a Ctf13 ortholog and form a predicted Cbf3 complex

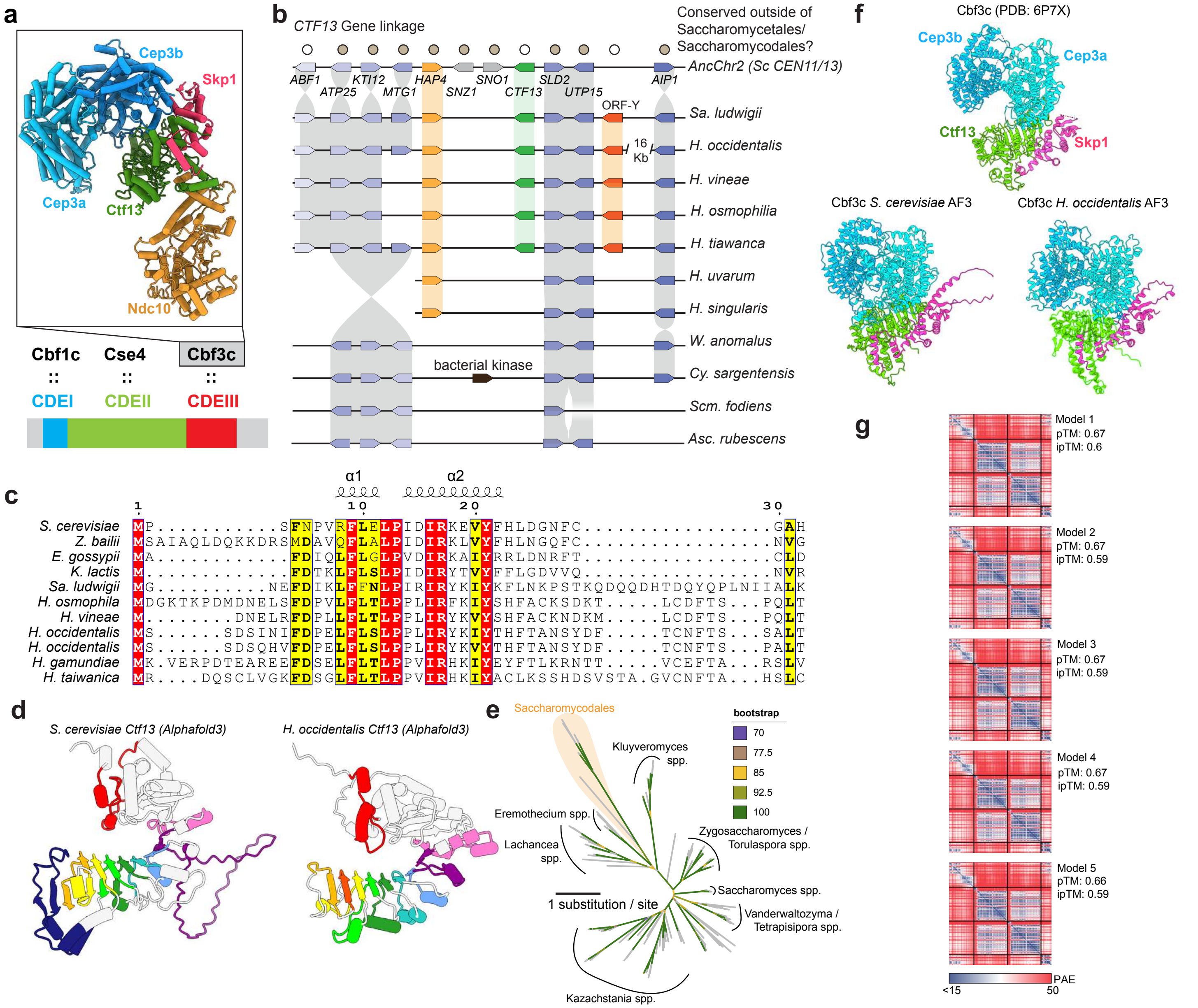

Supplemental Figure 9.  CTF13 is related to divergent yeast F-box LLR protein

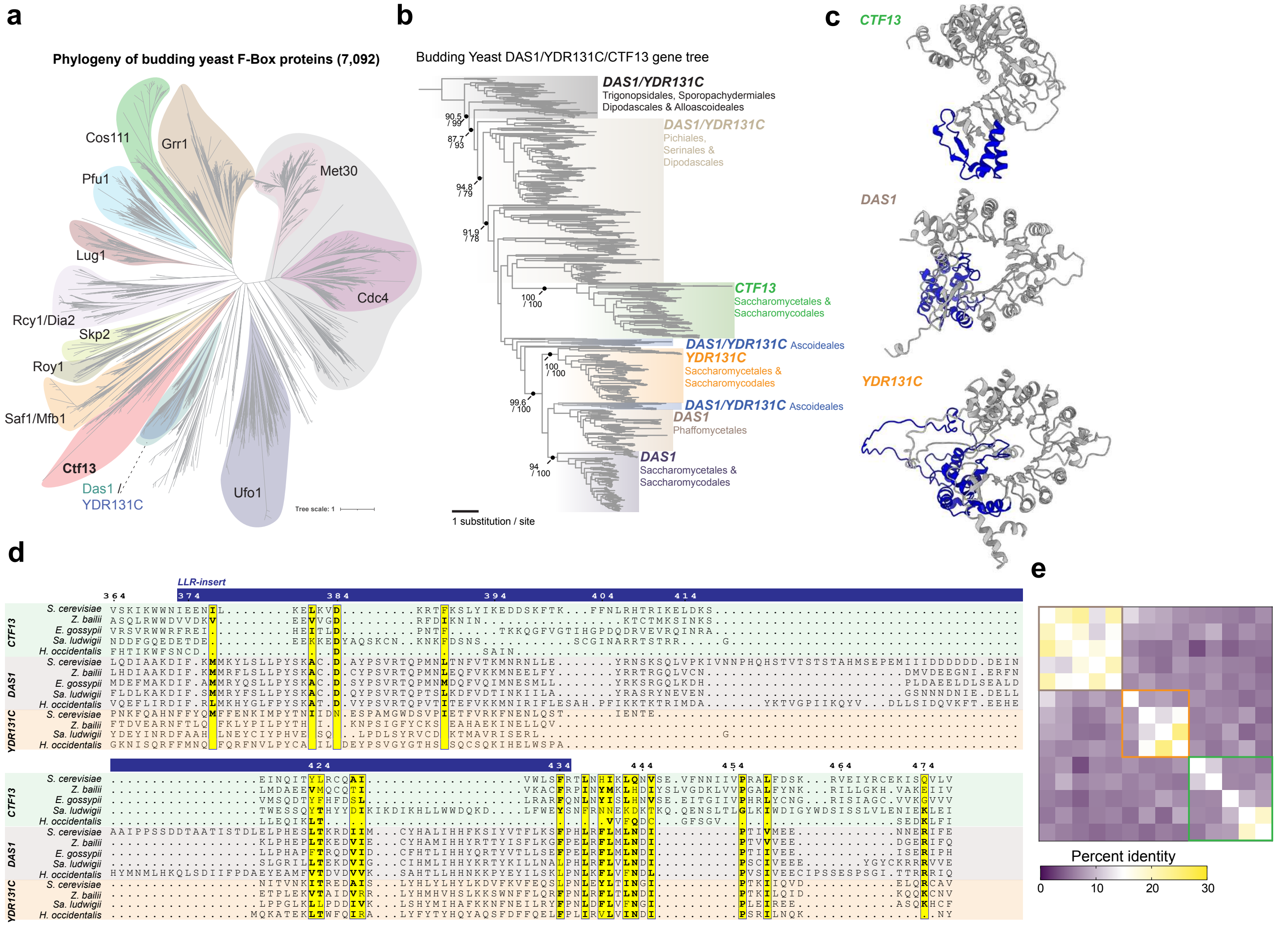

Supplemental Figure 10. Saccharomycodes ludwigii centromeres are diminutive Ty5 clusters

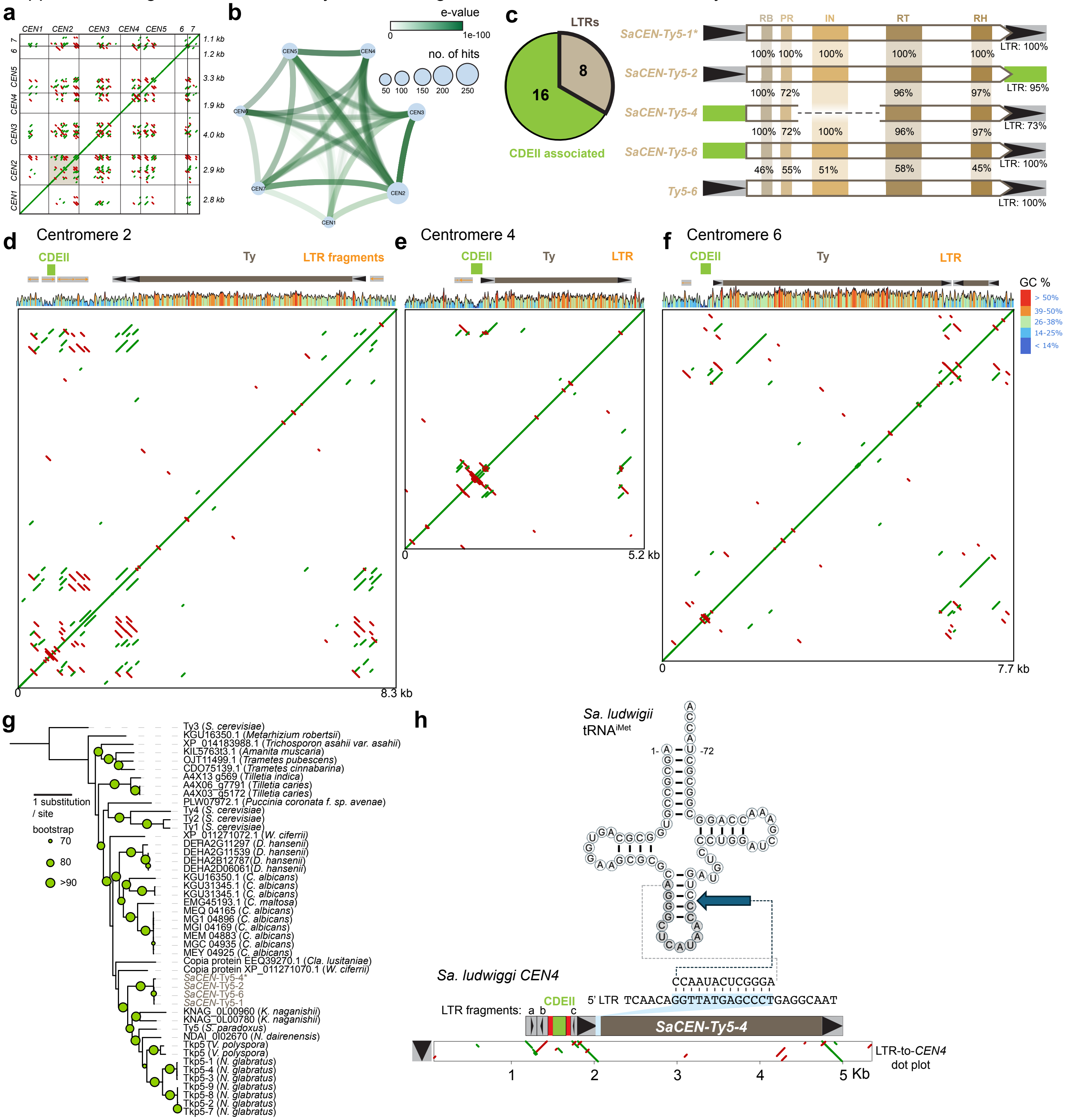

Supplemental Figure 11. Primary sequence support SaCEN-Ty5 as a yeast Ty5 family member

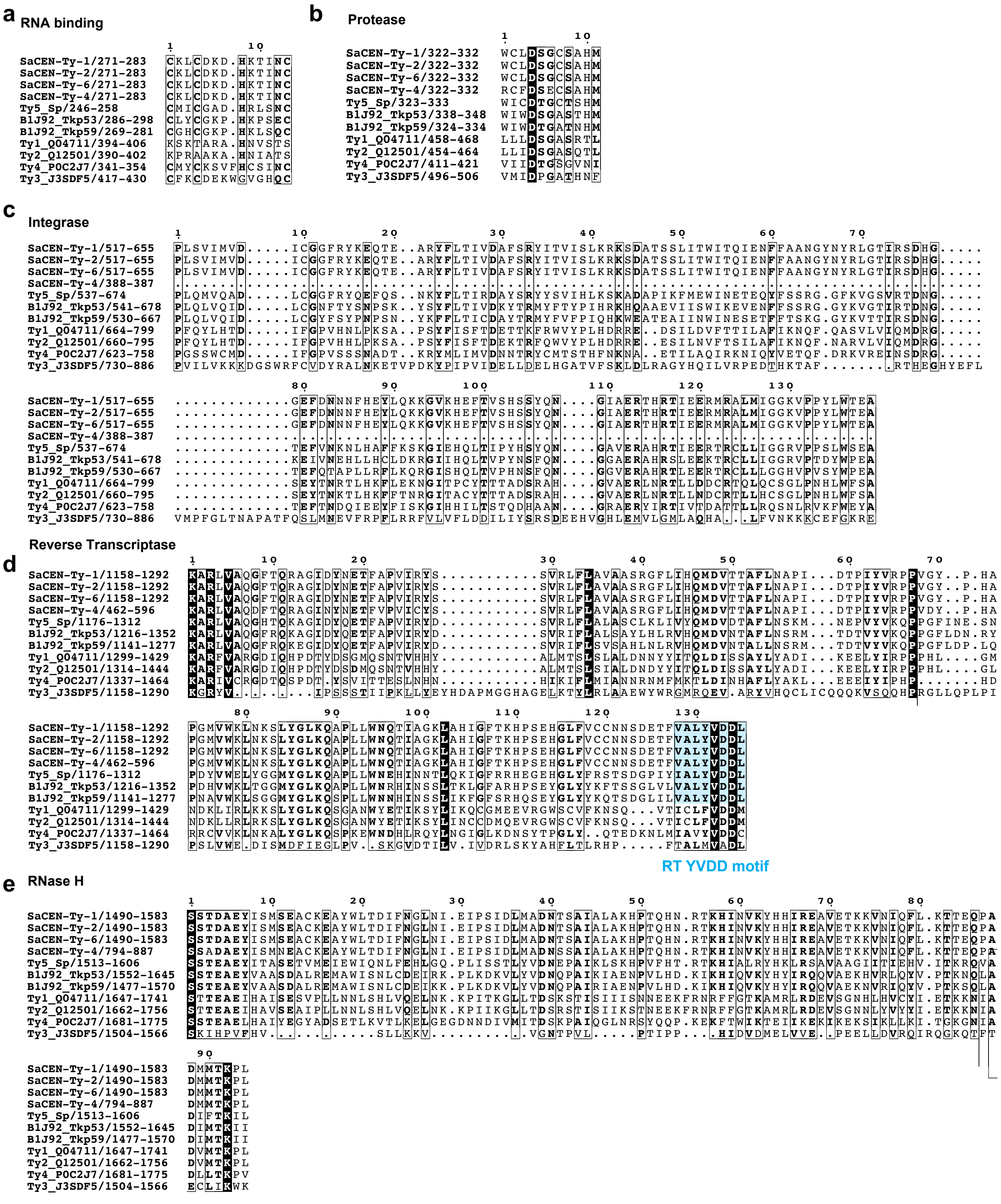

Supplemental Figure 12. Comparison of SaCEN-Ty5's from *Sa. ludwigii* strains

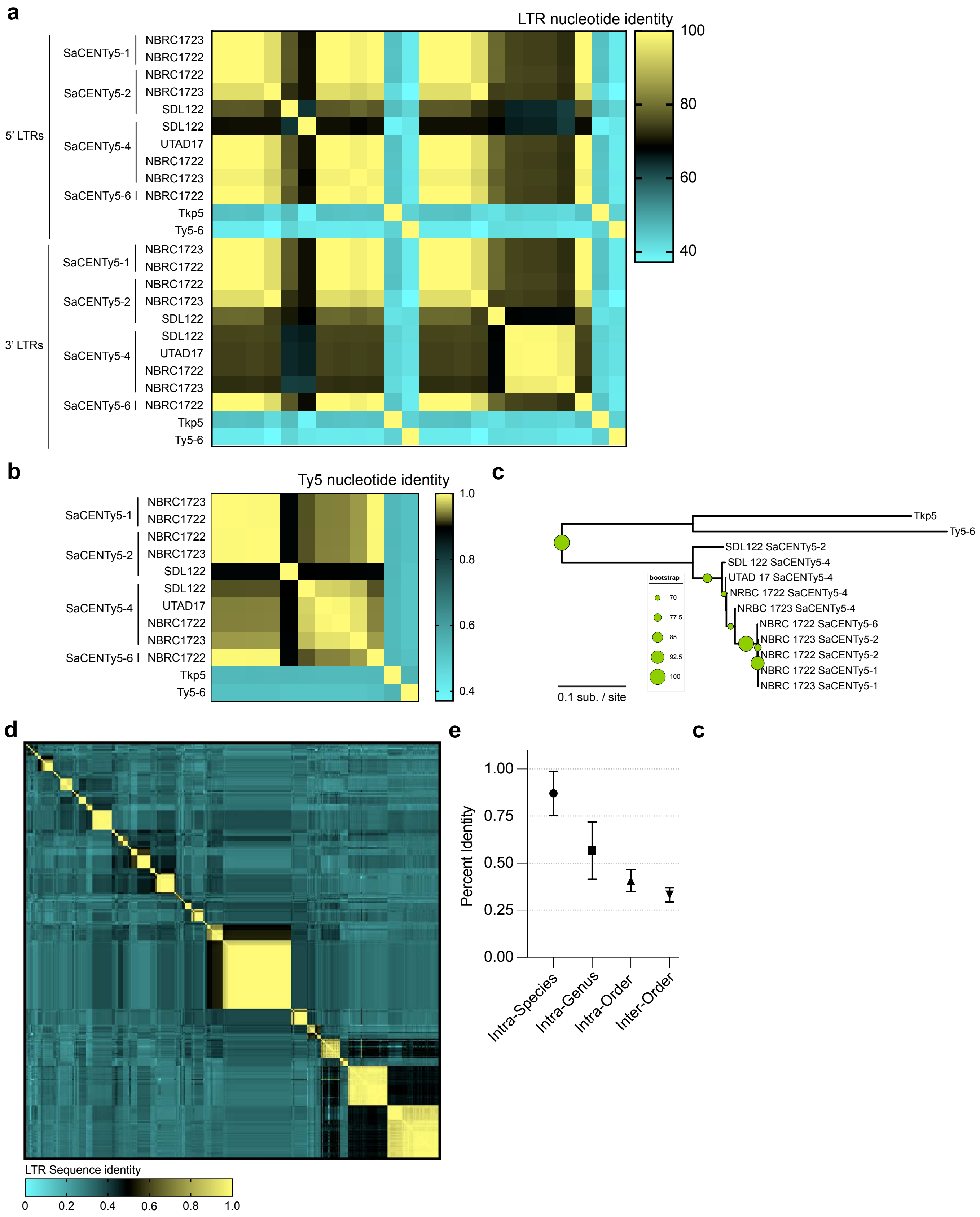

Supplemental Figure 13. Examples of Ty5 clusters from Pichiales, Serinales, Ascoideales, Phaffomycetales, and Saccharomycodales

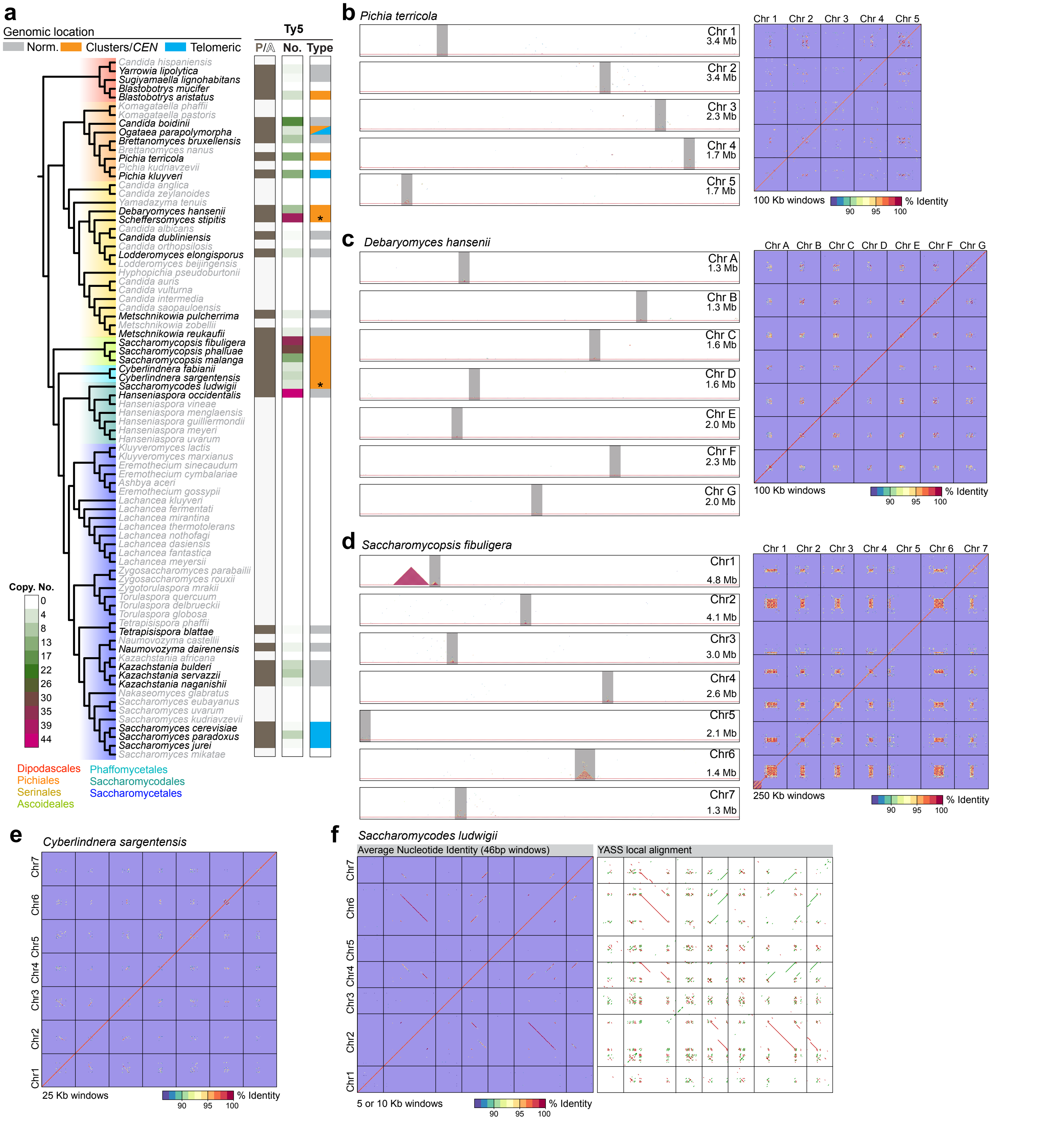

Supplemental Figure 14. Conserved gene synteny of centromere- and Ty5 cluster linked genes from diverse yeast species

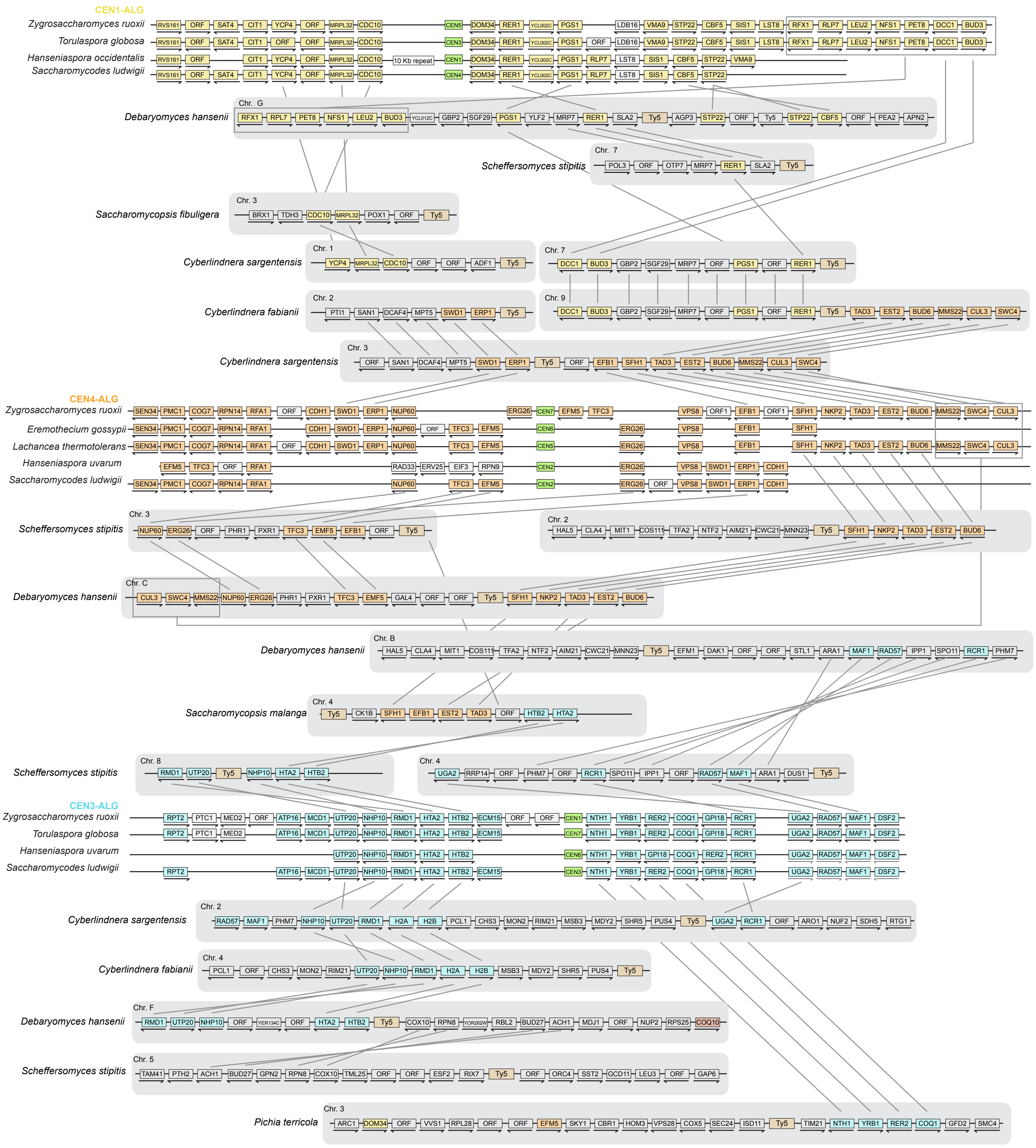

Supplemental Figure 15. Continued, conserved gene synteny of centromere- and Ty5 cluster linked genes from diverse yeast species.

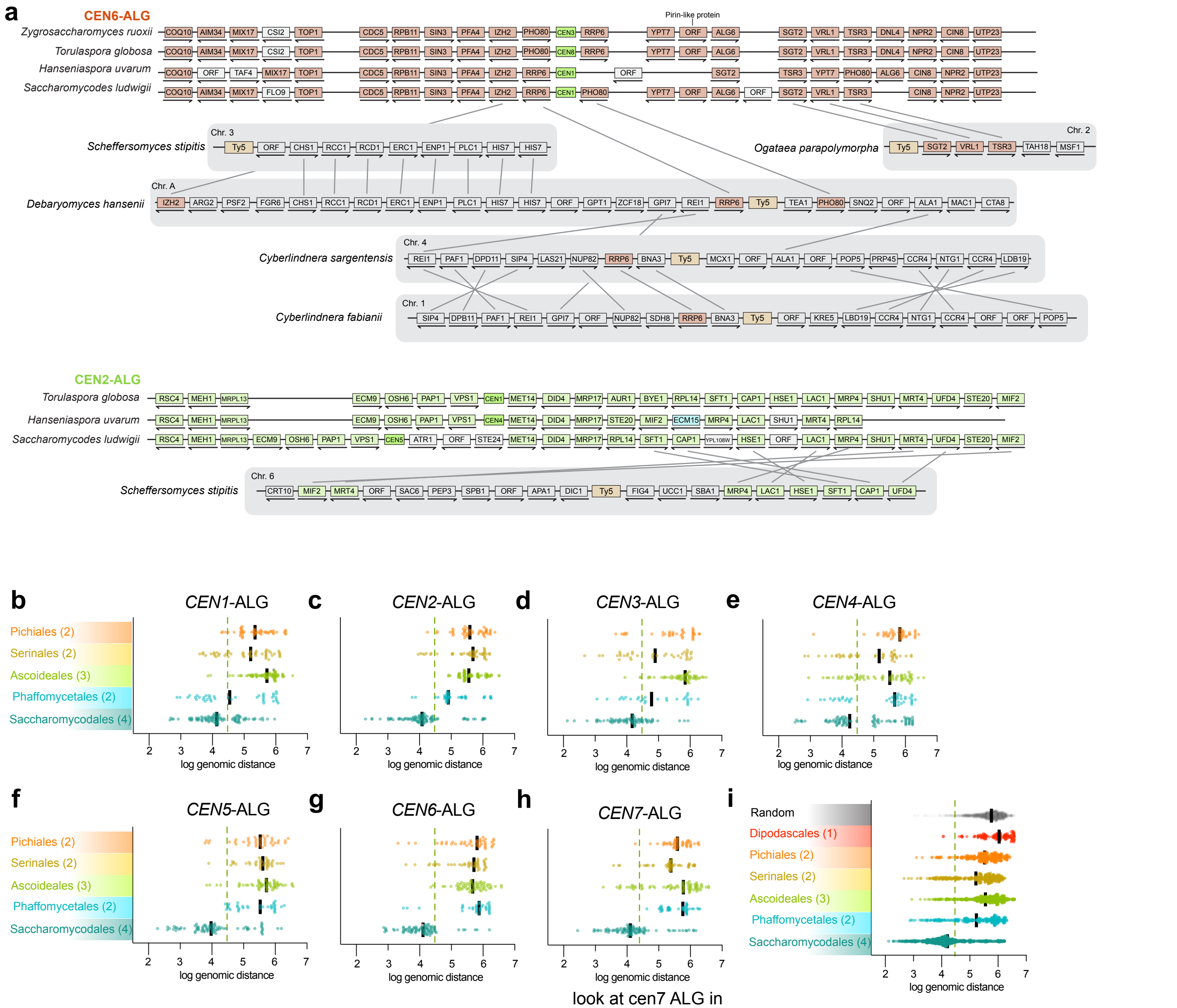


look at cen7 ALG in

Supplemental Figure 16. *Sa. ludwigii* CDEII is a divergent Ty5 LTR

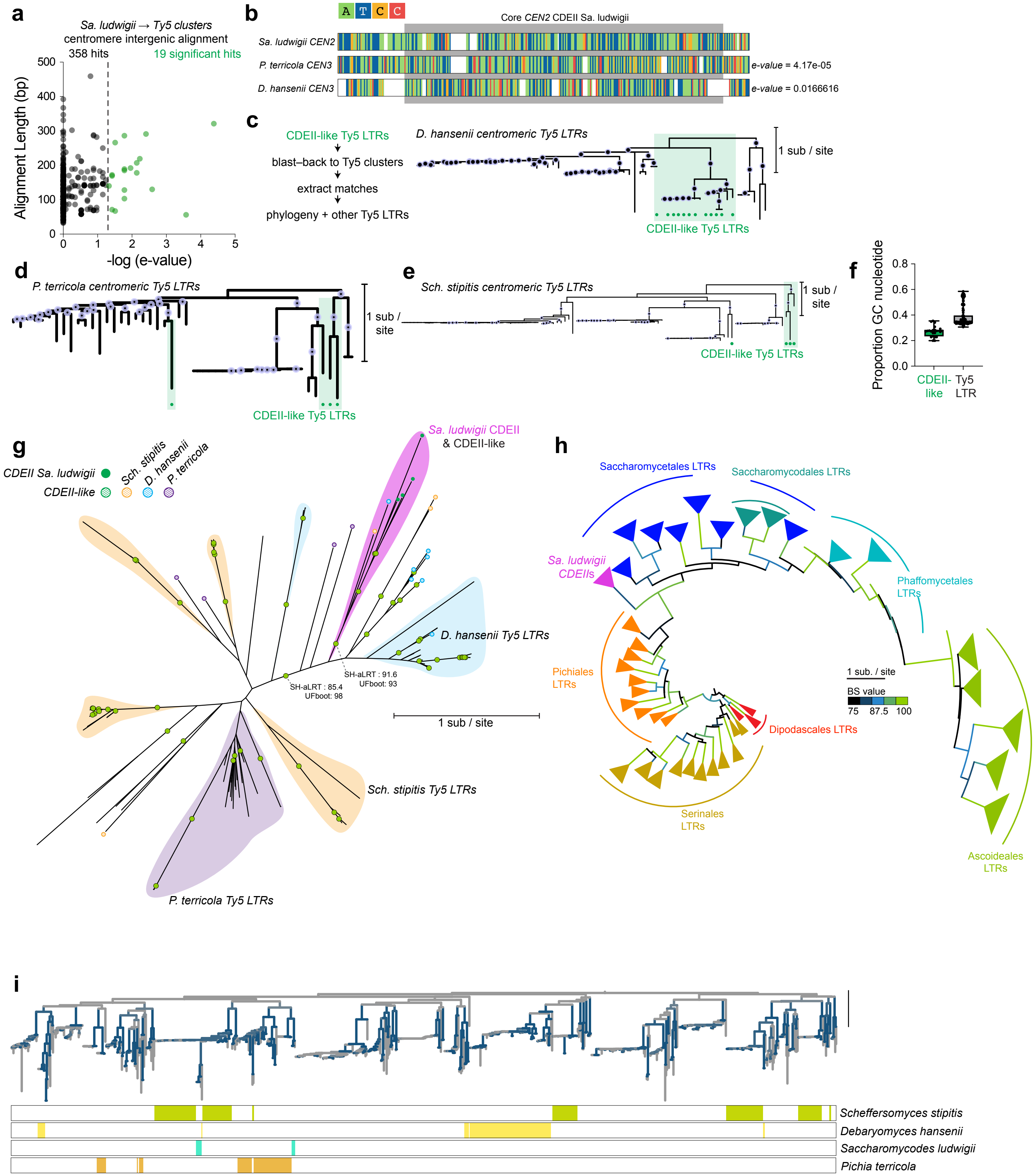

Supplemental Figure 17. Motif enrichment analysis of Saccharomycotina Ty5 LTRs

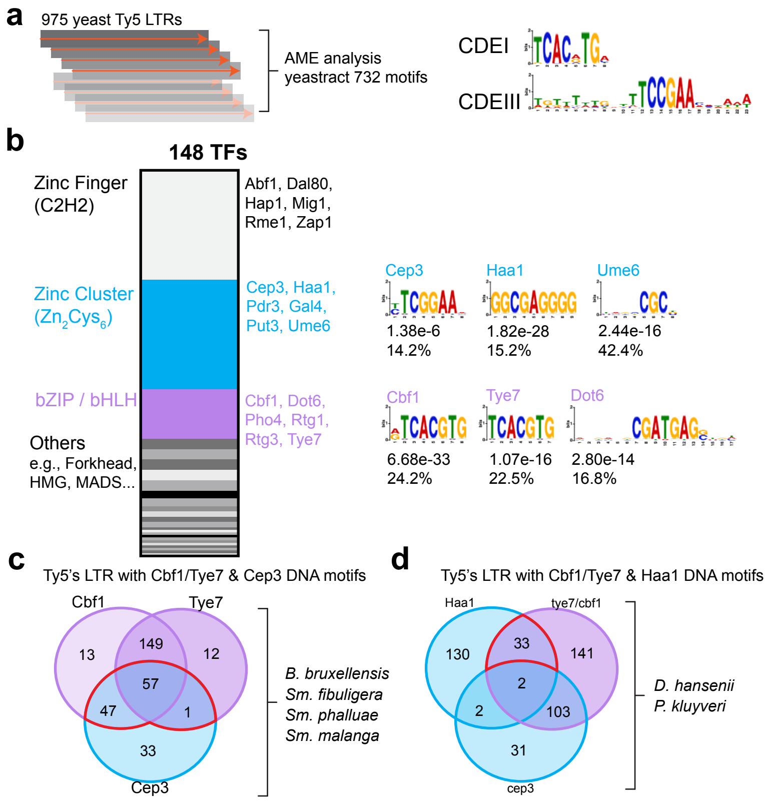

Supplemental Figure 18. Presence and Absence analysis of the indicated proteins across 1,154 genomes.

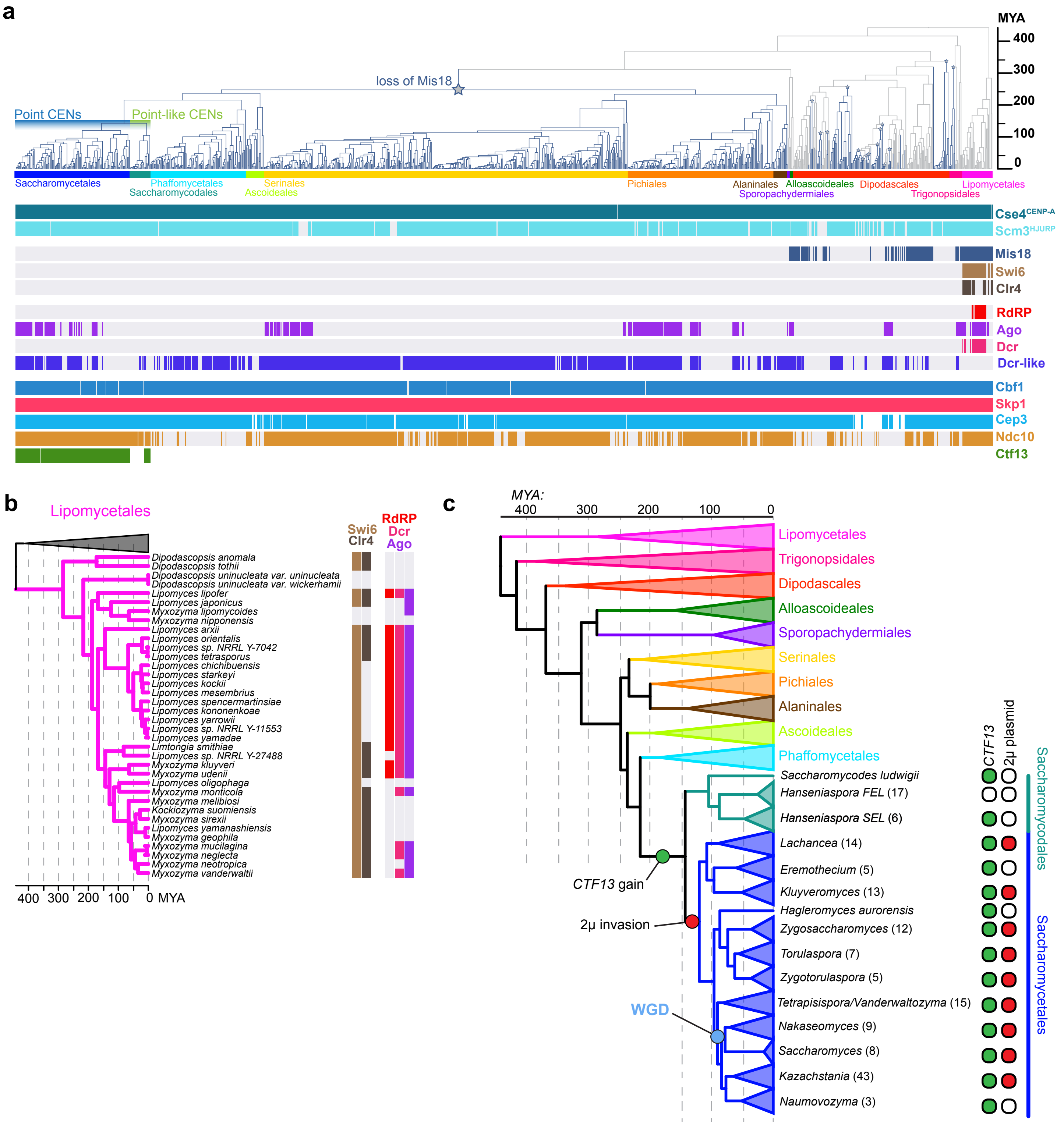

Figure S19. Linear mitochondrial genome of *H. uvarum*

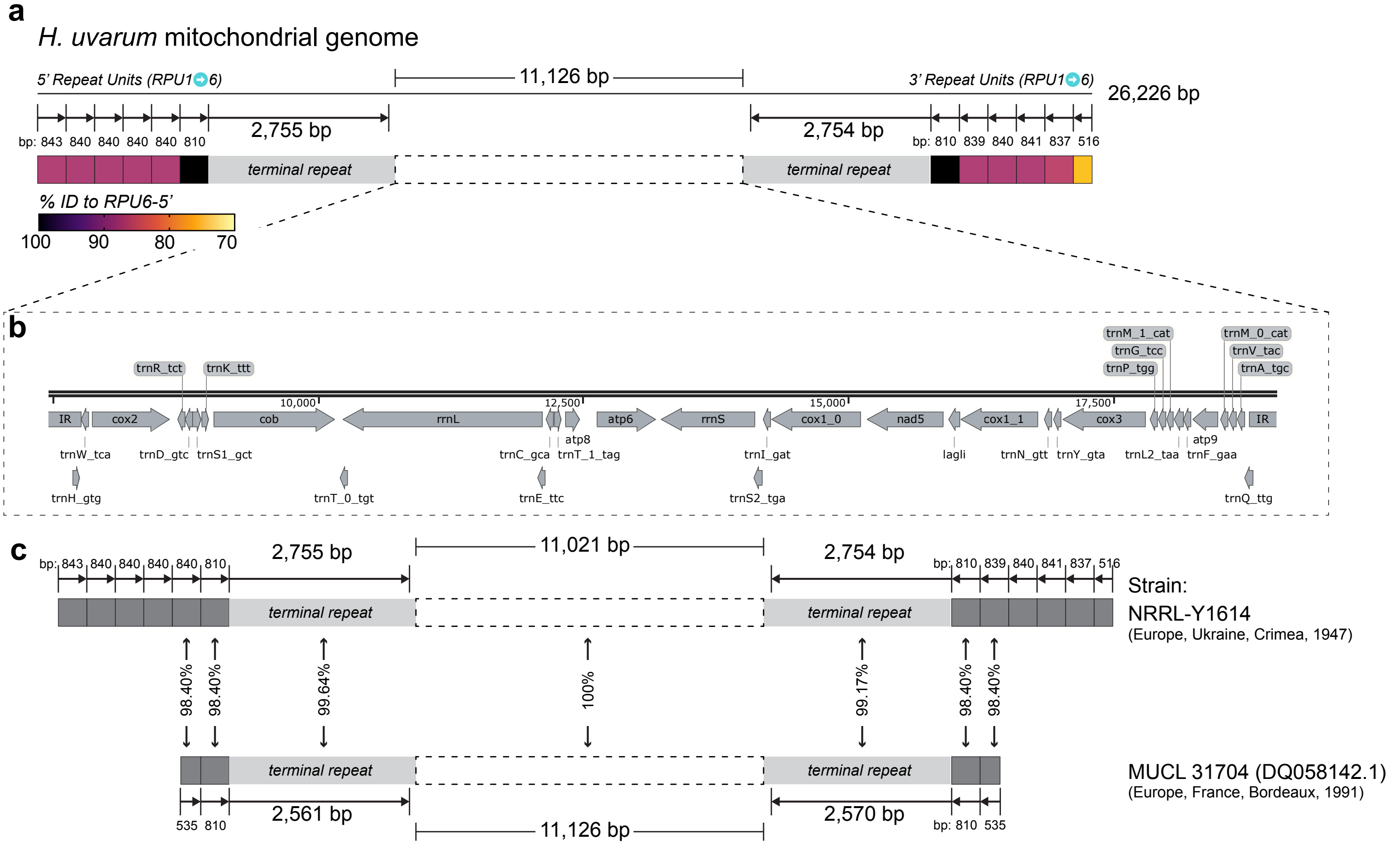

-log(evalue)

Figure S20. Characterization of the CEN-induced cellular arrest in *H. uvarum*

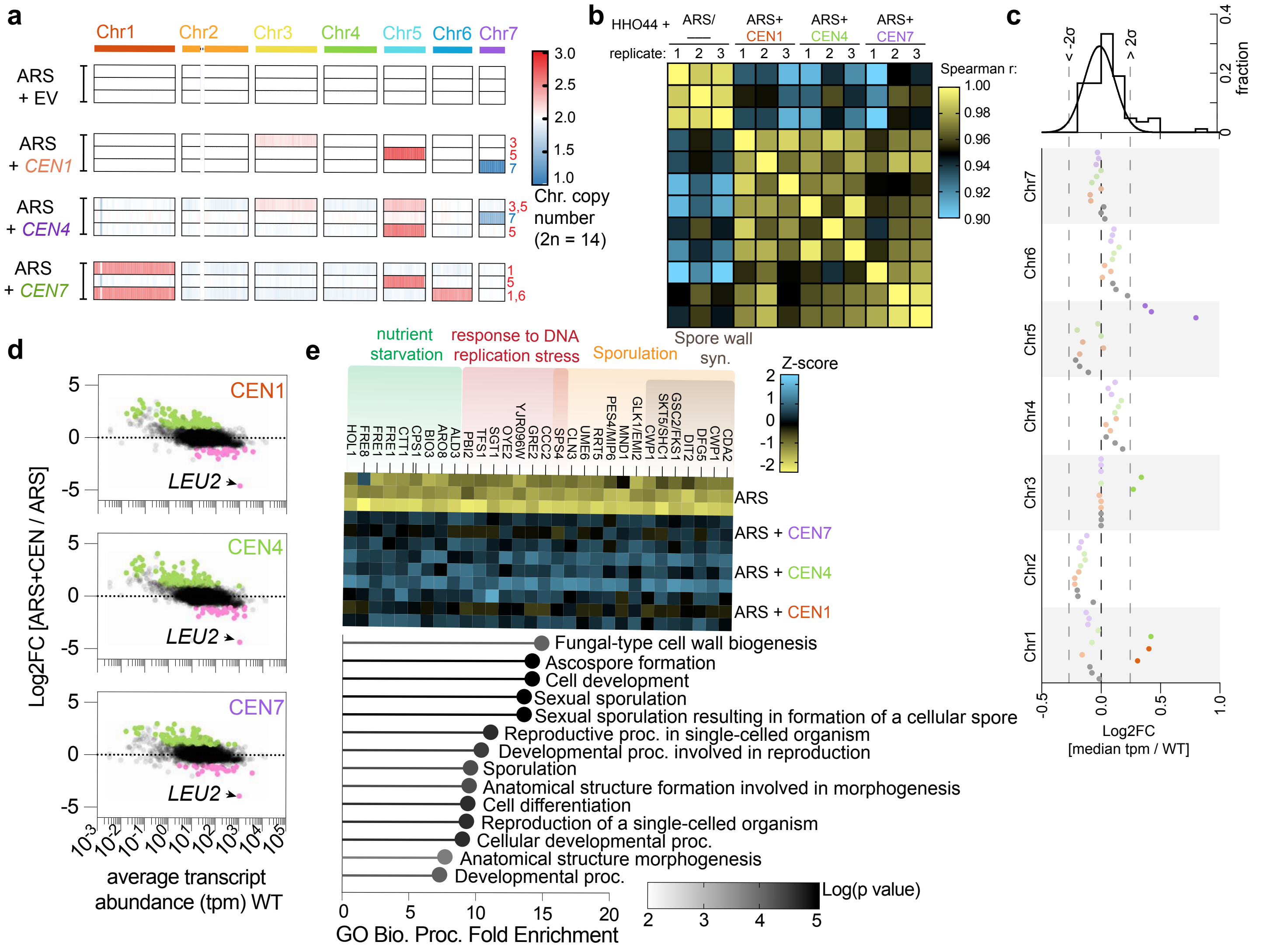
